# ATG2A interacts with RAB1A and ARFGAP1 positive membranes during autophagosome biogenesis

**DOI:** 10.1101/2025.03.24.645038

**Authors:** Devin M. Fuller, Yumei Wu, Florian Schueder, Burha Rasool, Shanta Nag, Justin L. Korfhage, Rolando Garcia-Milian, Katerina D. Melnyk, Joerg Bewersdorf, Pietro De Camilli, Thomas J. Melia

## Abstract

Autophagosomes form from seed membranes that expand through bulk-lipid transport via the bridge-like lipid transporter ATG2. The origins of the seed membranes and their relationship to the lipid transport machinery are poorly understood. Using proximity labeling and a variety of fluorescence microscopy techniques, we show that ATG2A localizes to extra-Golgi ARFGAP1 puncta during autophagosome biogenesis. ARFGAP1 itself is dispensable during macroautophagy, but among other proteins associating to these membranes, we find that RAB1 is essential. ATG2A co-immunoprecipitates strongly, albeit indirectly, with RAB1A, and siRNA-mediated depletion of RAB1A/B blocks autophagy downstream of LC3B lipidation, similar to ATG2A depletion. Further, when either autophagosome formation or the early secretory pathway is perturbed, ARFGAP1 and RAB1A accumulate at ectopic locations with autophagic machinery. Our results indicate that ATG2A engages a RAB1A complex on select early secretory membranes in support of autophagosome biogenesis.

**Significance Statement:** This study expounds upon the role of early secretory membranes in autophagosome biogenesis. The authors demonstrate that RAB1/ARFGAP1-positive membranes are essential to autophagy and are recruited to the phagophore assembly site at an early step of autophagosome biogenesis. These membranes interact with the bridge-like lipid transport protein ATG2A and are positive for LC3B and WIPI2, suggesting that RAB1 membranes are a direct source for autophagosome formation.

## Introduction

Macroautophagy is a cellular degradative pathway in which a seed vesicle grows to encapsulate a target cargo, closes to form the complete autophagosome, and subsequently fuses with the lysosome. This transient organelle forms *de novo* in approximately ten minutes and requires the delivery of millions of lipids^1^. ATG2 proteins function as bridge-like bulk lipid transport proteins^2–5^ and are essential during the membrane expansion step of autophagosome biogenesis. We propose that ATG2 bridges the ER and the developing phagophore, thereby allowing lipids to cross the cytosol through a long hydrophobic groove^3^. At the phagophore, lipids are equilibrated between both leaflets of the bilayer via the ATG9 scramblases^6–8^. As such, ATG9 is essential to macroautophagy and is a key component of the seed membranes that ultimately expand during ATG2-mediated lipid transport^9, 10^.

In addition to ATG9 vesicles, several other membrane sources have been proposed to be involved in autophagosome biogenesis. In particular, both genetics and biochemistry implicate an essential role for the early secretory pathway. In yeast, some COP-II vesicle and ER exit site (ERES) proteins contribute to starvation-mediated autophagy at a stage that is downstream of formation of the pre-autophagosome structure (PAS)^11, 12^. Furthermore, yeast ERES naturally colocalize with the autophagic cargo Ape1^13^, and that colocalization increases when autophagosome growth is stalled by depletion of essential factors^13, 14^. Autophagy proteins themselves collect at ERES; both imaging and cross-linking studies indicate a close association between the two membranes and disruption of ERES activity through depletion of Sec12 dramatically reduces Atg8 puncta formation^15^. Finally, the direct integration of yeast COPII-derived membranes into developing phagophores is suggested by the detection of the COP-II cargo Axl2-GFP, a transmembrane protein, at both phagophores and mature autophagosomes^16^.

In human cell lines, genetic or pharmacological disruption of the COPII-related small GTPase SAR1A reduces the frequency of autophagic events^17^. Following biochemical fractionation of cellular membranes, the ER-to-Golgi Intermediate Compartment (ERGIC) is especially competent for supporting in vitro lipidation by LC3B, the primary marker of autophagic structures^17, 18^ and clusters of COPII positive membranes can be seen accumulating adjacent to phagophores^19^. Like in yeast, these studies can be interpreted as a key role for ERES or ERGIC membranes in forming or expanding autophagosome membranes. As starvation-mediated macroautophagy drives substantial remodeling of both the ERES and ERGIC membranes^18, 20^, including the formation of subdomains with unique protein-dependent organelle-organelle contact sites^21^, it is possible that the ERES/ERGIC contribution to autophagy represents a distinct subcompartment of these membrane structures.

Presumably, any contribution of an early secretory membrane must ultimately coordinate with recruitment of ATG9 vesicles, in order to ensure that the lipid transport machinery is competent to expand the phagophore membrane. Several studies have postulated an early fusion intermediate in phagophore formation, such as the description of the HyPas compartment where vesicles bearing distinct autophagy proteins (ATG16L1 and FIP200) were observed to coalesce early in autophagosome biogenesis^22^. Intriguingly, when the lipid transporter ATG2 is genetically depleted from cells, all of the essential autophagy machinery including ATG9 concentrates into large compartments^9, 23–25^. Correlative light and electron microscopy of these structures indicate they are large assemblies of ATG9 vesicles surrounded by more complex membranes which harbor FIP200 and WIPI2, among other proteins^9, 26^. These complex membranes often adopt a cup-like shape, leading us to name them “proto-phagophores”^9^. The origins of the proto-phagophores have been unknown.

In this study, we demonstrate that ATG2A is enriched during autophagosome biogenesis at RAB1/ARFGAP1-positive membranes. Although both proteins can be found within the ERGIC compartment, proximity data and colocalization analyses in secretory pathway disrupted cells suggest that RAB1 and ARFGAP1 membranes involved in ATG2 recruitment are distinct from generic ERGIC membranes. Genetic depletion experiments indicate that RAB1 and ATG2 function at a similar step in autophagosome formation and further these proteins can be isolated together in a complex. Finally, both ARFGAP1 and RAB1 co-localize with proto-phagophores, implying these membranes can recruit much of the key early autophagy machinery and may be dynamically remodeled into cup-like morphologies.

## Results

### ATG2A is proximal to early secretory membranes

To determine the local proteome adjacent to ATG2A, we used a proximity labeling approach. We fused the biotinylation enzyme APEX2 and a green fluorescent marker to the amino-terminus of ATG2A (APEX2-EGFP-ATG2A) and expressed this construct in ATG2A/B double knockout cells (ATG2 DKO cells). In this way, the APEX2 enzyme will biotinylate cellular proteins in close proximity to the expressed ATG2A fusion protein^27^. To ensure that the APEX2-EGFP-ATG2A construct is active and correctly localized, we measured the restoration of macroautophagy in these cells by following LC3B flux and p62 clearance (Fig. S1A-C). While both LC3B-II and p62 dramatically accumulate in the ATG2 DKO cells, this accumulation is almost entirely reversed upon re-expression of APEX2-EGFP-ATG2A.

We also validated that biotinylation in these cells depends upon the presence of APEX-EGFP-ATG2A and a full reaction. In cells expressing APEX2-EGFP-ATG2A, biotin is only observed when both the biotin substrate (biotin phenol) and the reaction promoter (hydrogen peroxide) are included (Fig. S1D,E). Likewise, streptavidin-mediated immunoprecipitation revealed a strong enrichment of biotinylated material only when the full reaction conditions were employed (Fig. S1F). Furthermore, the anti-biotin and GFP signals colocalized establishing that biotinylation is favored at sites proximal to ATG2A (Fig. S1D).

Mass spectrometry of proteins isolated by streptavidin-mediated immunoprecipitation was performed using a three state quantitative SILAC experiment^27^ with light, medium, and heavy isotopes as follows: no hydrogen peroxide as a negative control, a full reaction in complete media, and a full reaction in starvation media and 100 nM bafilomycin. When comparing both the heavy and medium isotope states to the light state, mass spectrometry analysis revealed an abundance of autophagy related proteins in both full media and under starvation (Figs. 1A, S1G, respectively). The majority of these proteins were cargo adaptors such as NBR1, p62 (SQSTM1), and TAX1BP1, suggesting that the target cargo is immediately adjacent to the biogenesis machinery during autophagosome maturation. We also observed a clear enrichment of ERES or COPII proteins including SEC16A, SEC23A, SEC23IP, and SEC24B/C. Additional ERES proteins such as TFG and SEC31A were also enriched in some, but not all, of the three replicates. In addition, ARFGAP1, a COPI effector that is primarily localized to the Golgi but also found in earlier secretory membranes, was one of the two most enriched proteins in both full media and starvation. Both ARFGAP1 and the COPII proteins suggest ATG2 is localizing to the early secretory pathway.

**Figure 1.**
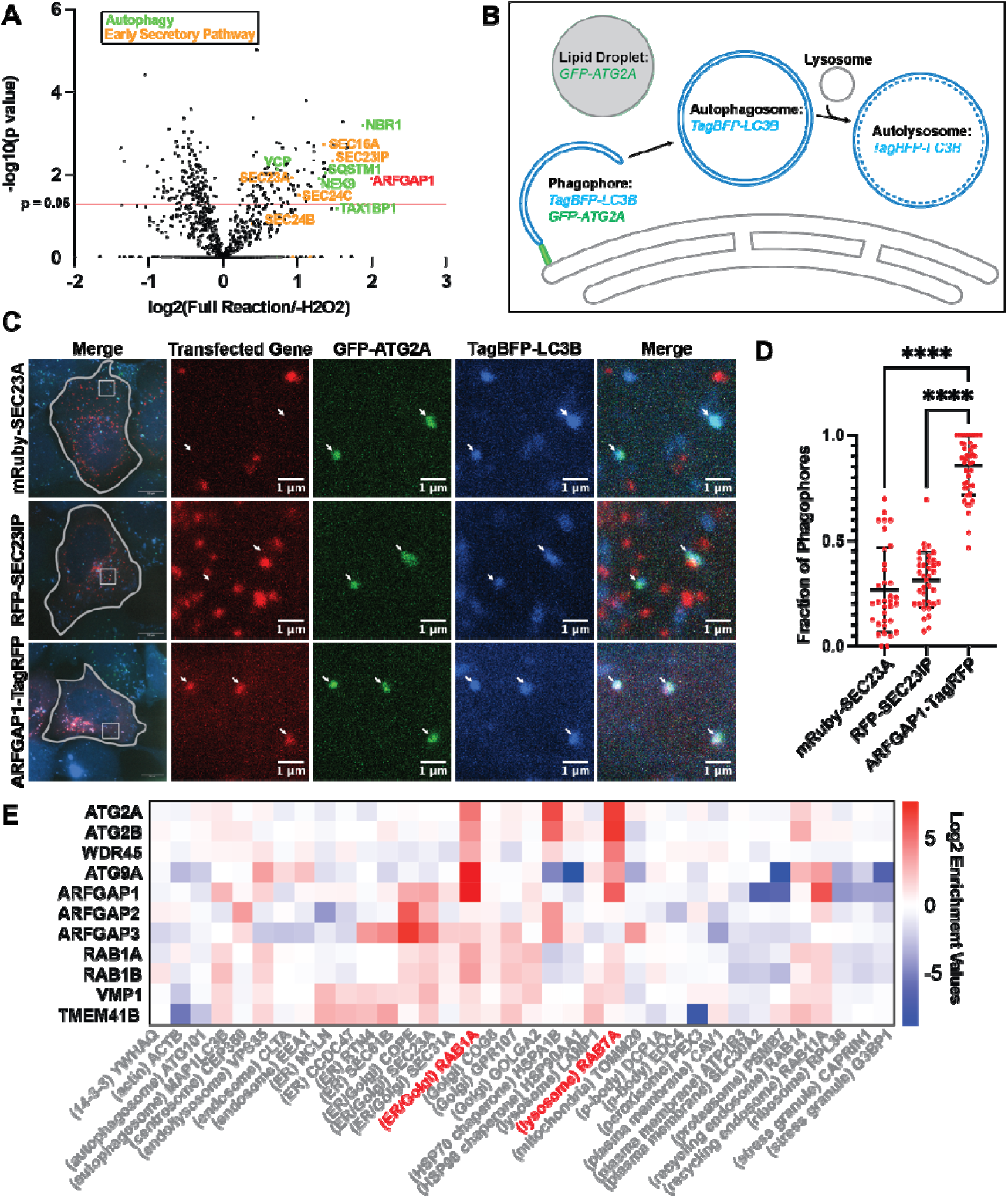
ATG2A is proximal to ARFGAP1-positive membranes. **(A)** Proximity labeling of HEK293 ATG2 DKO cells stably expressing APEX2-GFP-ATG2A was performed as previously described^27^. Three biological replicates were averaged and plotted as the log2 ratio between the full labeling reaction in complete media over cells that were not treated with hydrogen peroxide on the x axis, and -log10 of the p-value on the y axis. The red line demonstrates the p-value cutoff of 0.05. The proteomics revealed an enrichment of autophagy (labeled in green) and early secretory proteins (COPII/ER Exit Site proteins are labeled in orange, ARFGAP1 is labeled in red). **(B)** Strategy to identify phagophores by live cell imaging. Phagophores are defined as sites where both GFP-ATG2A and TagBFP-LC3B co-localize. **(C-D)** Live cell imaging of ATG2 DKO cells stably expressing GFP-ATG2A and TagBFP-LC3B and transfected with mRuby-SEC23A or RFP-SEC23IP/p125a or ARFGAP1-TagRFP. The cells were starved for 2-4 hours prior to imaging. The fraction of phagophores (white arrows) that colocalized with the third fluorescent protein (transfected gene) was quantified and plotted in (D). The gray lines show the cell periphery and the insets focus on representative phagophores. Statistical significance was assessed by one way ANOVA. *, adjusted P value <0.05. **, adjusted P value <0.01. ***, adjusted P value <0.001. ****, adjusted P value <0.0001. Data from three biological replicates were pooled for each condition. Maximum intensity projections of confocal images. **(E)** Global organelle profiling data from https://organelles.sf.czbiohub.org^32^. Whole organelle profiles were established by pulling down endogenously tagged proteins and performing quantitative proteomics. The enrichment value for each protein was calculated by taking the difference in median log2 LFQ values between the replicates and the null distribution.

### ATG2A is recruited to ARFGAP1 membranes during autophagosome biogenesis

As an orthogonal method to assess the proximity of ATG2A to early secretory components, we performed FLASH-PAINT, a super resolution technique with high multiplexing capabilities^28^. FLASH-PAINT offers sub-organelle resolution while simultaneously allowing us to screen >10 proteins. ATG2 DKO cells stably expressing GFP-ATG2A were starved and then immunostained for autophagic proteins (GFP-ATG2A, p62, and LC3B), ERES proteins (Tango1, SEC24C, SEC31A, SEC16A, TFG), and markers of COPI-positive membranes (ARFGAP1 and COPI). All proteins were concurrently labeled with primary antibodies that were in turn identified by their association with secondary antibodies with unique DNA barcodes (Fig. S1I). A fluorophore conjugated imager DNA strand and adaptor strands were added to target the imager to the correct secondary antibody. Eraser strands were added in between rounds of imaging to outcompete the adaptor and allow for sequential labeling. In this way, the sub-cellular localization of all ten proteins could be established within the same image.

In these images, the Golgi can easily be identified as a perinuclear ribbon staining strongly for both ARFGAP1 and COP1 (magenta and purple respectively, Fig. S1J). In addition, there are bright puncta throughout the cell (pink arrow heads) stained with TFG and SEC31A, indicating that they are ERES. ARFGAP1 and COPI frequently also accumulated at ERES (made more obvious in the top-right inset, where TFG and SEC31 are removed), consistent with previous reports that COPI vesicles facilitate ER to Golgi transport in human cells^29–31^. GFP-ATG2A is mostly cytosolic, however small accumulations of ATG2 could be found at ERES and were generally observed to be proximal to ARFGAP1/COP1 (inset; brackets). Note that clusters of LC3B and p62 are also present. Occasionally, non-ERES accumulations of p62 could be observed in close proximity to LC3B and ATG2A (lower insets), and here again ATG2 is proximal to ARFGAP1.

Critically, we can directly measure the nominal proximity of each protein relative to the others under investigation by quantifying the median distance between all ten tested proteins within 200 nm of each individual protein (Fig. S1K). All early secretory pathway proteins closely associate with each other, including the COPI proteins ARFGAP1 and β-COP. As the majority of ATG2A was cytosolic, its average distances from other proteins were greater, however, within this cohort of candidate proteins, we found that ATG2A is most closely associated with p62 and ARFGAP1 (15nm and 16nm respectively; top row of Fig. S1K), exactly consistent with the APEX screen.

The APEX and FLASH-PAINT datasets provided a clear but static result that ATG2A is recruited to ARFGAP1 positive structures. To determine whether this recruitment was coincident with phagophore formation, we stably expressed GFP-ATG2A and TagBFP-LC3B in ATG2 DKO cells in order to identify phagophores. ATG2A primarily localizes to phagophores and lipid droplets, whereas LC3B primarily localizes to phagophores, autophagosomes and autolysosomes. Where these two proteins colocalize suggests a growing phagophore in the process of membrane expansion (Figs. 1B). To assess co-localization with phagophores, we expressed candidate early secretory proteins labeled with red fluorophores (Fig. 1C,D; see Fig. S1H for analysis strategy). ARFGAP1-TagRFP colocalized with ∼90% of actively forming phagophores, whereas only ∼30% of phagophores were coincident with the ERES markers mRuby-SEC23A or RFP-p125a/SEC23IP. Further, while the ARFGAP1 signal was largely concentrated in the Golgi, phagophores most frequently colocalized with extra-Golgi ARFGAP1 puncta.

Finally, we compare our results to the subcellular localization of ARFGAP1 established by global organelle profiling available at the database at https://organelles.sf.czbiohub.org^32^. ARFGAP1 is enriched on autophagic membranes positive for LC3B as well as endolysosomal and early secretory membranes (Fig. 1E). Strikingly, ARFGAP1 was most highly enriched on membranes isolated via RAB1A immunoprecipitation. The autophagic proteins ATG2A, ATG2B, WDR45, and ATG9A, all of which are involved in membrane expansion of the autophagosome, were also highly enriched on RAB1A membranes. These collective results establish that both endogenous ARFGAP1 (organelle isolation, APEX, FLASH PAINT) and overexpressed ARFGAP1 (live imaging) naturally associate with ATG2A, including during phagophore biogenesis.

### Autophagic flux depends upon RAB1 but not ARFGAP1

To look for a potential functional role for ARFGAP1 in autophagy, we measured autophagic flux using the HaloTag dropout assay in which liberation of a free HaloTag (HT) is indicative of lysosome-mediated proteolysis of the full-length (FL) protein^33^. We stably expressed HaloTag7-mGFP-LC3B or HaloTag7-mGFP in WT HEK293 cells and then used siRNA to knockdown various regulators or potential regulators of macroautophagy initiation (Fig. 2A,B, S2A). Despite efficient knockdown, siRNA against ARFGAP1 had no effect on flux; the ratio of free HaloTag to full-length HaloTag7-mGFP-LC3B following ARFGAP1 knockdown was similar to both nontransfected (NT) and nontargeting siRNA controls. In contrast, knockdown of the autophagy initiation factor FIP200 dramatically suppressed dropout of the HaloTag. We also looked for an interaction between ARFGAP1 and ATG2A, but even when both proteins were overexpressed, we could not detect significant interaction via co-immunoprecipitation (co-IP; Fig. 3A,B). Thus, it is likely that ARFGAP1 localizes to a key membrane involved in autophagosome biogenesis but is not itself a player in initiating autophagosome formation. To examine whether the other isoforms of ARFGAP1 could redundantly contribute to autophagosome biogenesis, we knocked ARFGAP1, ARFGAP2, and ARFGAP3 down individually and collectively. Knockdown of ARFGAP2, ARFGAP3 and ARFGAP1-3 marginally increased autophagic flux (Fig. S2B,C), suggesting either no role or a minor role as negative regulators of autophagy.

**Figure 2.**
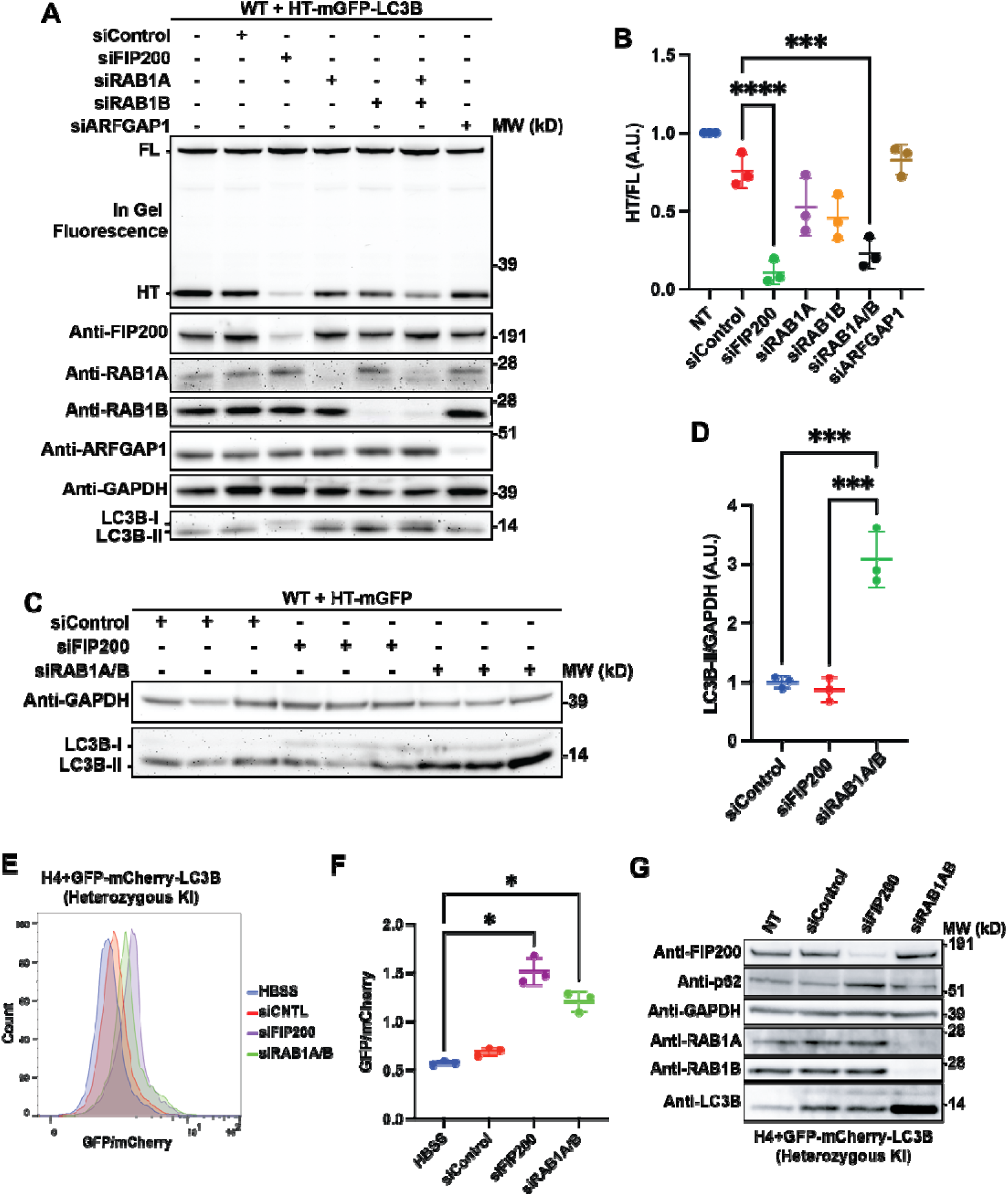
Autophagic flux depends upon RAB1 but not ARFGAP1. **(A-B)** In gel fluorescence image (top panel) and immunoblots demonstrating that siRNA KD of RAB1 inhibits autophagic flux in cells starved in EBSS for four hours. In gel fluorescence reveals HaloTag7s moiety bound to the TMR ligand. Indicated bands are full length (FL) HT-mGFP-LC3B protein and HaloTag7 after cleavage (HT) in the lysosome. The ratio between FL and HT demonstrates autophagic flux, quantified in (B). Statistical significance was assessed by one way ANOVA. *, adjusted P value <0.05. **, adjusted P value <0.01. ***, adjusted P value <0.001. ****, adjusted P value <0.0001. Data from three biological replicates were collected for each condition**. (C-D)** Immunoblot of lysates from three biological replicates demonstrating that siRNA KD of RAB1 results in an accumulation of LC3B-II in cells starved in EBSS for four hours. The band intensity of LC3B-II was normalized against the intensity of GAPDH for each lane. To normalize each replicate, each individual value was divided by value of the non-transfected sample. Statistical significance was assessed by one way ANOVA. *, adjusted P value <0.05. **, adjusted P value <0.01. ***, adjusted P value <0.001. ****, adjusted P value <0.0001. **(E-G)** FACS data demonstrating that RAB1A/B KD results in a block in autophagic flux. H4 cells with a heterozygous GFP-mCherry-LC3B endogenous tag were transfected with siRNA for three days, following which they were starved in HBSS for four hours and analyzed by FACS. The ratio between GFP and mCherry is an indicator of autophagic flux, as the GFP signal is quenched in the lysosome. This ratio is displayed in histograms in (E) for one replicate and as averages in (F). The KDs of each protein can be seen in (G).

**Figure 3.**
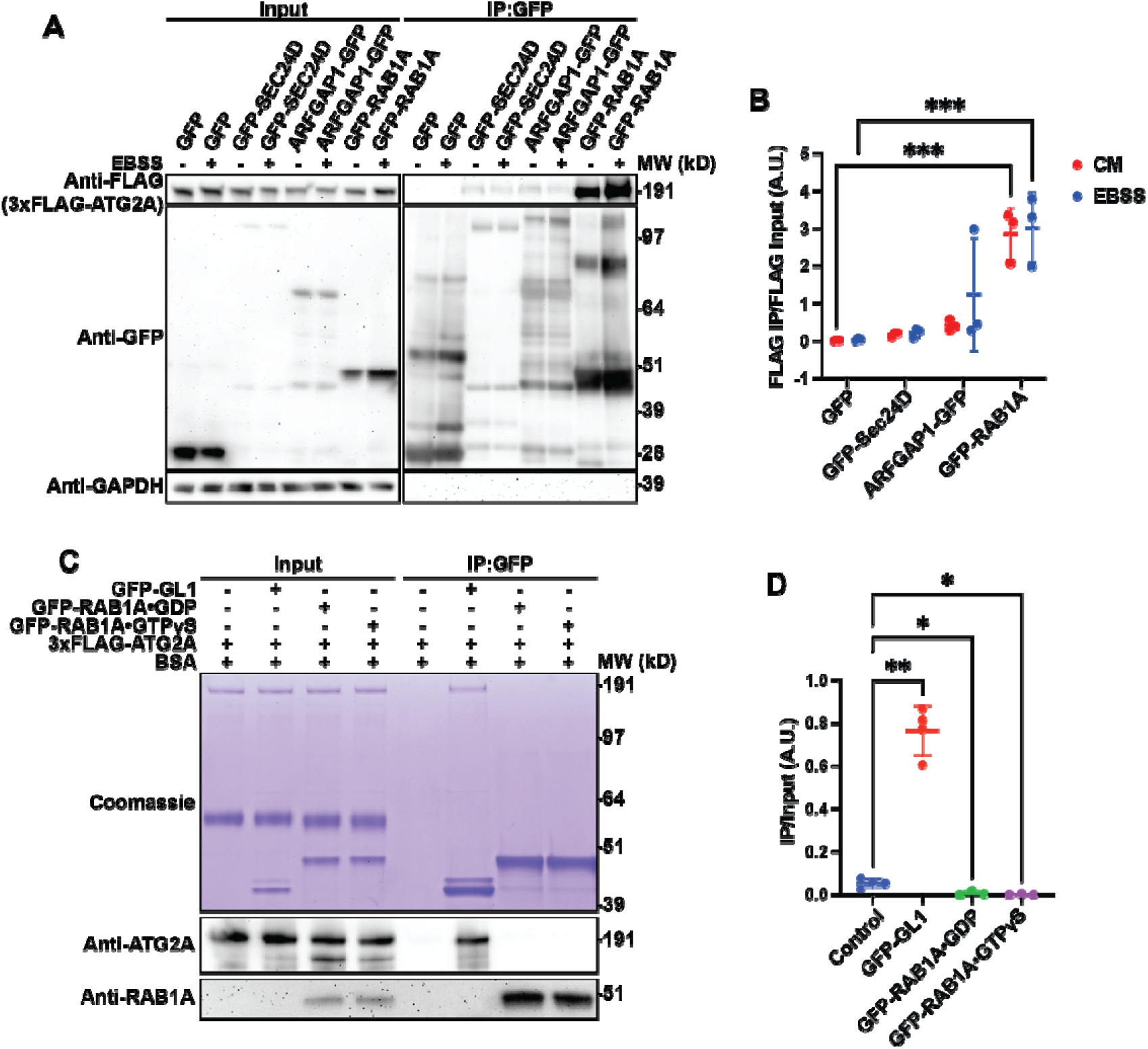
ATG2A interacts indirectly with RAB1A. **(A-B)** Immunoblot showing CoIP between GFP-RAB1A and 3xFLAG-ATG2A compared to the same reaction with other GFP-tagged early secretory proteins. The cells were incubated with either complete media or EBSS for four hours prior to harvesting. For the CoIP, ∼3 mg of cell lysate was used per condition (slight deviations in total amount between replicates, but not between lanes). To quantify the results, the intensity of the 3xFLAG-ATG2A IP signal was ratioed against the input signal. To normalize each replicate, each individual value was divided by the average of the replicate. Statistical significance was assessed by two way ANOVA. *, adjusted P value <0.05. **, adjusted P value <0.01. ***, adjusted P value <0.001. ****, adjusted P value <0.0001. The media was not a significant source of variation (p = 0.3535). **(C-D)** A fully reconstituted CoIP reaction between GFP tagged bait proteins and 3xFLAG-ATG2A reveals no direct interaction between RAB1A and ATG2A. GST tagged GFP-GABARAPL1 (GL1) and GFP-RAB1A were purified from *E. Coli*, during which process the GST tag was proteolytically removed. The purified proteins were then used in a CoIP reaction using GFP trap beads as in (A), but with no cell lysate present in the reaction. BSA was used to block the beads and as a negative control. To quantify the results, the intensity of the 3xFLAG-ATG2A IP signal was ratioed against the input signal. To normalize each replicate, each individual value was divided by the average of the replicate. Statistical significance was assessed by one way ANOVA. *, adjusted P value <0.05. **, adjusted P value <0.01. ***, adjusted P value <0.001. ****, adjusted P value <0.0001.

As shown in (Fig. 1E), both the ATG2/ATG9 lipid transport apparatus as well as ARFGAP1 are associated to RAB1A-positive membranes. RAB1 proteins mediate ER to Golgi trafficking in the early secretory pathway. The yeast RAB1 homologue, Ypt1, is essential to autophagy^34, 35^, while in mammals, genetic depletion of either RAB1A or RAB1B modestly reduces autophagic flux^36, 37^. Thus, we tested the functional and physical interaction of RAB1 proteins with ATG2. Consistent with previous results, siRNA knockdown of either RAB1A or RAB1B in our system partially inhibited flux in the Halo dropout assay (Fig. 2A,B). Strikingly, knockdown of both isoforms simultaneously was nearly as effective as knocking down FIP200, establishing that RAB1 proteins play an essential role in supporting starvation induced autophagic flux. This result was consistent with but more pronounced than previous work examining LC3B flux following RAB1A/B double knock down^38^. As an orthogonal method to examine the role of RAB1 in LC3B flux, we used H4 neuroglioma cells with a GFP-mCherry-LC3B heterozygous knock in reporter^39^.

Following knock down and starvation, we examined the ratio between GFP and mCherry by FACS. The low pH of the lysosome quenches the fluorescence of GFP at a higher rate than mCherry, resulting in a low ratio under starvation conditions as seen in the HBSS and siCNTL conditions (Fig. 2 E-G). In contrast, knock down of FIP200 or RAB1A/B resulted in an increased ratio, confirming that these proteins are necessary for proper autophagic flux. Of note, the RAB1A/B knockdown did not block flux as efficiently as the FIP200 KD. If RAB1 is essential for autophagy, this result suggests that either very few copies of RAB1 are needed to support autophagy or that another gene could be compensating for the lack of the two RAB1 isoforms.

In addition to following tagged LC3B, we also immunoblotted for endogenous LC3B in each of our siRNA experiments. We found that knockdown of either RAB1 isoform, and especially knockdown of both together leads to a clear accumulation of the membrane conjugated form of LC3B, LC3B-II (Fig. 2A,G). As LC3B overexpression can alter endogenous LC3B patterns, we confirmed this increase in a cell line expressing only HaloTag7-mGFP (Fig. 2C,D). Knockdown of RAB1A and RAB1B drove a threefold increase of LC3B-II (Fig. 2D). We also confirmed this phenotype in WT HEK293 cells in a classical LC3B flux assay with bafilomycin (Fig. S2D), where we found elevated LC3B-II levels in all conditions.

Increasing LC3B-II levels without autophagic flux is indicative of a block to macroautophagy at a late stage, downstream of initiation, and is therefore easily distinguished from the block induced by depleting FIP200 (where LC3B-II levels go down). Intriguingly, LC3B-II levels also increase when ATG2 proteins are genetically depleted (Fig. S1A,B), suggesting ATG2A and RAB1A might function concurrently. Indeed, overexpressed GFP-RAB1A could strongly immunoprecipitate flag-tagged ATG2A in our cells, indicating these proteins are in a shared complex (Fig. 2E,F). However, ATG2A and RAB1A (bound either to GDP or GTPγS) do not interact in a fully reconstituted system (Fig. 3C,D), suggesting that the interaction is indirect. In contrast, ATG2A and GABARAPL1 cleanly interact in the *in vitro* CoIP.

We also tested several other proximity hits from the biotinylation analysis for interaction with flag-tagged ATG2A, including GFP-tagged copies of ERES resident proteins SEC23A, SEC31A, SEC16A, Sar1 and all four homologs of Sec24 (Fig. S3 A-D). We found that ATG2A co-immunoprecipitated weakly with many of them, the strongest of which was SEC24D. Note that this SEC24D interaction was barely above the background of pull-down with GFP alone (Fig. 3A,B). Thus, consistent with proximity and imaging analyses, ATG2 proteins are in the general proximity of the ERES machinery, but stable protein-protein interactions appear to be limited to RAB1.

### Disruption of the early secretory pathway results in the recruitment of autophagic proteins to RAB1A-positive tubular membranes

These results suggest that ATG2 engages an early secretory membrane, distinct from the ERES, in a RAB1-dependent manner. We next tested whether disruption of the early secretory system would impact the localization of ATG2A. To this end, we overexpressed SAR1BH79G, the catalytic dead mutant of the small GTPase SAR1B that recruits the COPII coat. SAR1BH79G binds the membrane and recruits the COPII coat, but is unable to release, thereby trapping COPII onto the membrane. Overexpression of this construct has been shown to reduce autophagy in starved cells^17, 36^. Consistent with those results, overexpression of SAR1BH79G-mOrange2 diminished autophagic flux of both a bulk cargo (HaloTag7-mGFP reduced ∼20%) and of LC3B (HaloTag7-mGFP-LC3B reduced by ∼30%) by the HaloTag degradation assay^33^ (Figs. 4A,B). In a parallel assay following autophagic flux instead by the changing fluorescence of tandem tagged GFP-mCherry-LC3B, we detect a similar, but statistically insignificant trend. As both assays are likely sensitive to SAR1BH79G expression levels, which vary in our transient transfection system, we also tested how LC3 puncta accumulation correlates with SAR1BH79G expression on a per cell basis. LC3B puncta number were lowest in cells expressing high amounts of SAR1BH79G (Figs. S4A,B), consistent with a block to autophagosome formation when the secretory pathway is disrupted rather than a later impact on autophagosome clearance.

**Figure 4.**
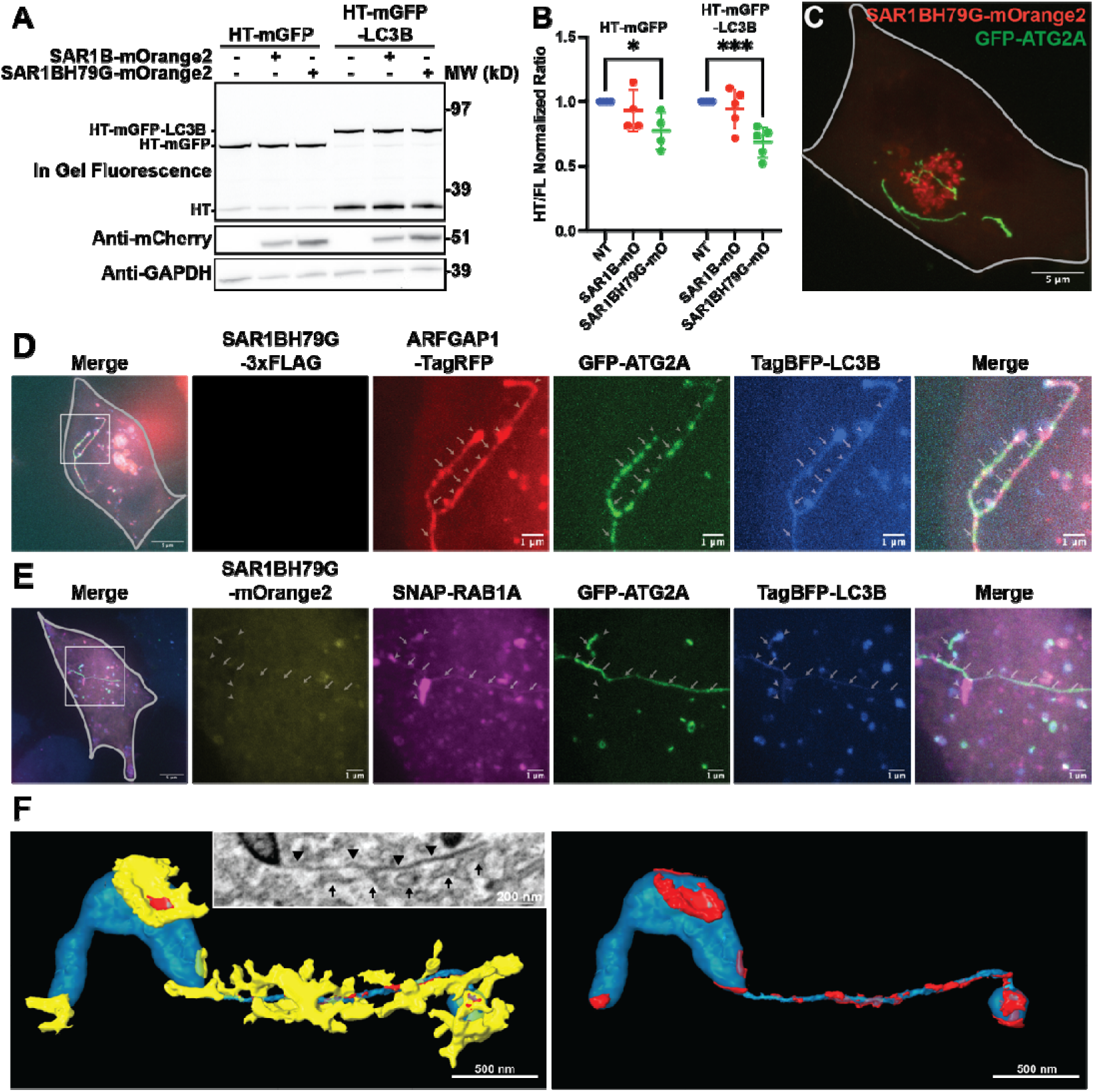
ATG2A, LC3B, ARFGAP1, and RAB1A localize to tubules following disruption of the early secretory pathway. **(A-B)** In gel fluorescence image (top panel) and immunoblots demonstrating that SAR1BH79G-mOrange2 (mO) overexpression negatively affects autophagic flux in cells starved in EBSS for four hours. The ratio between the FL and the free HT represents the autophagic flux in each condition (as in Fig. 2 A/B) and is plotted in (B). Statistical significance was assessed by two way ANOVA. *, adjusted P value <0.05. **, adjusted P value <0.01. ***, adjusted P value <0.001. ****, adjusted P value <0.0001. Data from four biological replicates were collected for each of the HT-mGFP conditions and five replicates were collected for each of the HT-mGFP-LC3B conditions. **(C)** Live cell imaging of ATG2 DKO cells stably expressing GFP-ATG2 that were transfected with SAR1BH79G-mOrange2 and starved for two hours. Note that GFP-ATG2A forms long tubules that do not colocalize with the SAR1BH79G-mOrange2 signal. Maximum intensity projection of a confocal image. Gray line was added to show cell outline. **(D-E)** Live cell imaging demonstrates the localization of ARFGAP1-TagRFP (D) and SNAP-RAB1A (E) on GFP-ATG2A/TagBFP-LC3B positive tubules following SAR1BH79G overexpression. Arrows denote thin GFP-ATG2A positive tubules whereas arrow heads denote thicker tubules that lack most GFP-ATG2A but contain high levels of LC3B, ARFGAP1, and RAB1A. Maximum intensity projections of confocal images. **(F)** CLEM-FIBSEM of cells prepared as in (C). GFP-ATG2A fluorescence corresponds to a thin tubular membrane that is connected to a thicker tubule. The small inset in the left image shows a single SEM slice. The arrowheads are pointing to the tubule which is segmented and displayed in turquoise in the larger image, whereas the arrows are pointing at adjacent ER tubules which are segmented and displayed in the larger image in yellow. At many points, these ER structures come into very close apposition with the tubule which is denoted in red. The ER structures are removed in the right image for better visualization of the ATG2A positive tubule.

Unexpectedly, in a small subset of transfected cells (1.5±0.8%; calculated from the images in Fig. S4A,B), overexpression of SAR1BH79G under starvation conditions shifted the localization of GFP-ATG2A from a largely punctate distribution to long, tubular objects that were SAR1BH79G negative (Fig. 4C). GFP-ATG2A decorated these tubules with varying intensities approximating a beads-on-a-string pattern (e.g. arrows in Fig. 4D,E). Fluorescently tagged forms of LC3B, ARFGAP1 and RAB1A also decorated these tubules. However, these three proteins displayed the brightest intensities where ATG2A was dimmer (e.g. arrowheads in Fig. 4D,E). These three proteins, but not ATG2A, also decorated larger structures that appeared to be contiguous with the tubule, including bulbous regions at the tubule termini.

To determine the ultrastructure of these regions, we subjected GFP-ATG2A-expressing cells to correlative light and focused ion beam scanning electron microscopy (CLEM-FIB-SEM). The GFP-ATG2A fluorescence signal corresponded to thin, tubular membranes (Fig. 4F and S4D), which is consistent with the finding that ATG2A preferentially binds highly curved membranes^40^. The tubule is surrounded by ER elements, many of which appear to be in very close contact (Fig. 4F, S4C). GFP-ATG2A signal was more intense in regions with closely adjacent ER, consistent with a possible contact site localization. The tubule continued into larger structures at either end, which were GFP-ATG2-negative, and morphologically similar with the bulbous structures harboring ARFGAP1, RAB1A and LC3B in fluorescence images.

Thus, although unusual, these rare tubules afforded an opportunity to demonstrate that ATG2, ARFGAP1 and RAB1A target the same membrane, easily resolvable even at confocal resolution. ATG2 collects where ER is closely adjacent, and LC3B accumulates with RAB1A, implying a topological organization with LC3B and RAB1A decorating the same structure, suggesting that RAB1A membranes may be a component or progenitor of the phagophore membrane.

### ARFGAP1 and RAB1A accumulate around pre-autophagic structures in ATG2 DKO cells

We previously published that very small, ATG9- and LC3B-positive vesicles accumulate in large clusters with the cargo-adaptor p62 in ATG2 DKO cells^9^. This cluster is surrounded by a shell of more complicated membranes which we termed “protophagophores” because they colocalize with other autophagy proteins such as WIPI2 and because many of them appear to adopt a cup-like shape in TEM images. Beyond this protophagophore shell, the entire structure is surrounded by ER (illustrated in Fig. 5A). Our work and others have shown that essentially all proteins involved in autophagosome biogenesis accumulate with these clusters, including FIP200, ATG14, ATG9A, ATG16L, and p62^9, 23, 24^, suggesting they are repositories of autophagosome biogenesis material awaiting the arrival of ATG2 proteins. If early secretory membranes contribute directly in any way to the biogenesis event, they may also concentrate at these sites.

**Figure 5.**
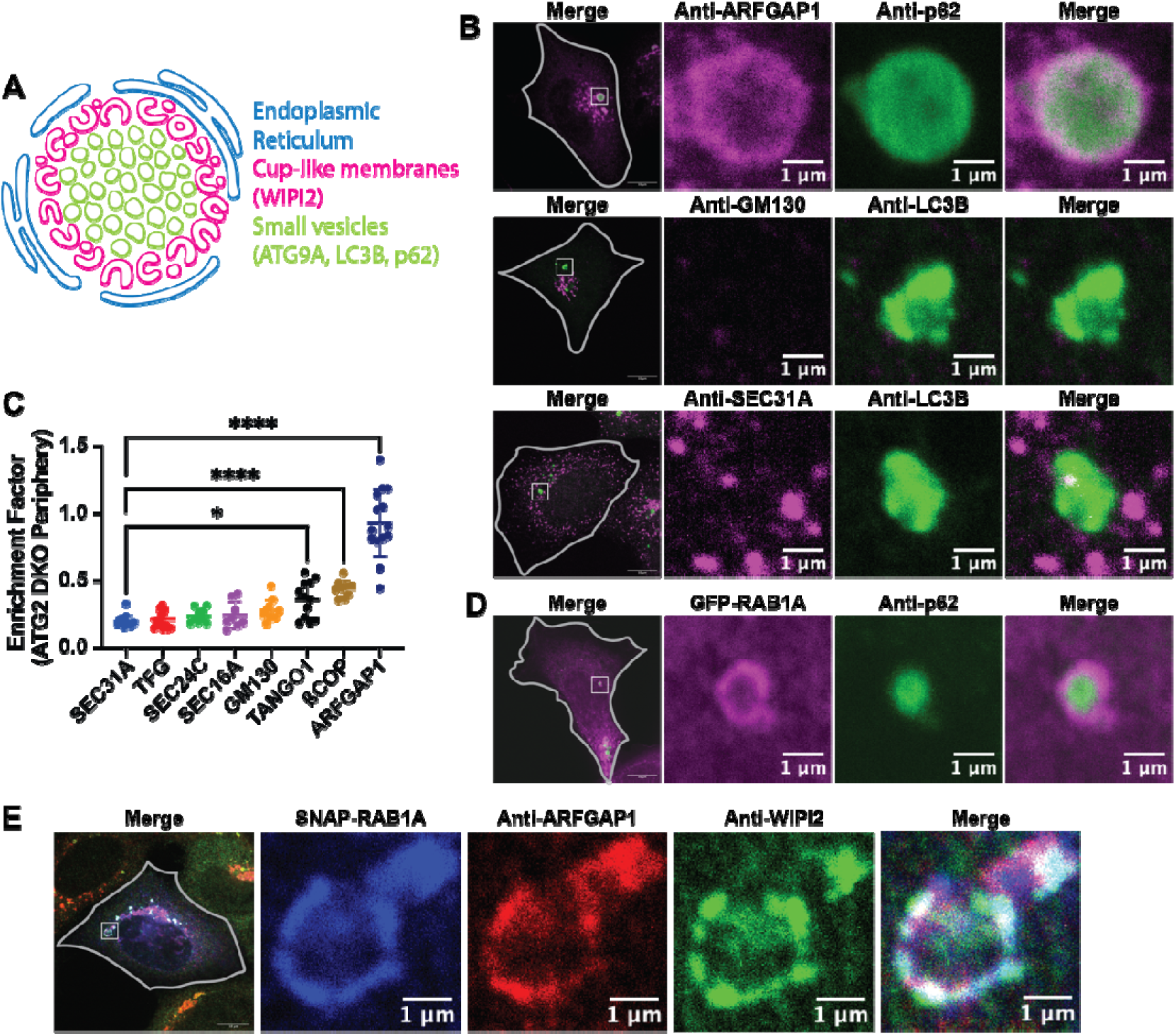
ATG2 deletion results in the accumulation of ARFGAP1 and RAB1A on autophagic membranes. **(A)** Schematic of the membranes that accumulate at the ATG2 DKO compartment based on the TEM images in Olivas et al^9^. The core of the compartment contains small vesicles that stain for ATG9A, LC3B, and p62. The periphery of the compartment is comprised of more complicated cup shaped membranes that stain for WIPI2. The entire compartment is enwrapped by ER tubules. **(B)** Immunofluorescence images demonstrating the distribution of early secretory membranes around the ATG2 DKO compartment which is marked by either p62 (mouse antibody) or LC3B (rabbit antibody). The images presented are single confocal slices of the center of the ATG2 DKO compartment. The gray lines show the cell periphery and the insets focus on the largest accumulation of p62 or LC3B in the cell. **(C)** Quantification of the images in (B) and in (Fig. S4A). The enrichment factor is equal to the mean of the given protein at the periphery of the ATG2 DKO compartment compared to its mean throughout the cell. Statistical significance was assessed by one way ANOVA. *, adjusted P value <0.05. **, adjusted P value <0.01. ***, adjusted P value <0.001. ****, adjusted P value <0.0001. Data from three biological replicates were pooled together for each of the conditions. **(D-E)** Immunofluorescence images demonstrating the enrichment of overexpressed GFP-RAB1A (D) or SNAP-RAB1A (E) at the periphery of the ATG2 DKO compartment as determined by the presence of p62 (D) or WIPI2 (E). Presented as a single confocal slice. The gray lines show the periphery of the cell and the insets focus on ATG2 DKO compartments that are distal from the Golgi.

We therefore performed IF against various markers of the early secretory pathway in ATG2 DKO cells to look for colocalization with this compartment (represented by staining for p62 (mouse antibody) or for LC3B (rabbit antibody); Figs. 5B-D). We found that ER exit site markers such as SEC16A, SEC24C, SEC31A, and TFG were not enriched around the compartment, neither was the Golgi protein GM130 (Figs. 5B-C and S5A). ßCOP was modestly enriched, but at levels that by eye, did not suggest encapsulation of the compartment the way autophagy proteins collect. In addition, overexpressed mEmerald-ERGIC53 did not localize to the ATG2 DKO compartment (Fig. S5B). In contrast, ARFGAP1 targeted all of the ATG2 DKO compartments, forming a robust spherical shell at the periphery of the p62 signal in HEK293 ATG2 DKO cells (Fig. 4B-D), or permeating the entire compartment in HeLa ATG2 DKO cells^41^ (Fig. S5C).

We were unable to find a strong RAB1A signal by immunofluorescence, however, when overexpressed, tagged versions of RAB1A also formed a sphere around the HEK293 ATG2 DKO compartment (Fig. 5D. Both ARFGAP1 and SNAP-RAB1A colocalized with WIPI2 (Fig. 5E), consistent with a model whereby these membranes arrive at the PAS prior to ATG2A recruitment and subsequent autophagosome biogenesis.

### ARFGAP1 accumulates at the ATG2 DKO compartment independently of the Golgi

The vast majority of ARFGAP1 is found at the Golgi, but ATG2A specifically localizes to extra-Golgi ARFGAP1 puncta (Fig. 1C, S1J). Consistent with a non-Golgi origin for these membranes, both natural and induced perturbations of Golgi structure have no impact on the recruitment of ARFGAP1 membranes to the p62-rich compartments in ATG2 DKO cells. First, in the course of these studies we observed that ARFGAP1 persistently colocalized with p62 in cells undergoing mitosis (Fig. 6A,B,C), even as the Golgi fragments and dissipates. Second, when cells were treated with brefeldin A to inhibit the formation of the Golgi^42^, ARFGAP1 again persistently colocalized with p62 (Fig. 6B,C). As brefeldin A specifically targets GBF1, the GEF for the ARF proteins^43^, this result also suggests that ARFGAP1 localization to the p62 compartment is independent of GTP bound ARF proteins. Third, overexpression of GFP-SidM/DrrA, an enzyme from *Legionella Pneumophila* which acts as both a GEF^44, 45^ and a GDF^46^ for RAB1, fragments the Golgi^44^ and sequesters RAB1 to the plasma membrane^45^. We found that following GFP-SidM expression in our cells, ARFGAP1 maintained colocalization with p62, even as the Golgi-associated ARFGAP1 as well as the Golgi-marker GM130 were completely dissipated (Fig. 6D, S6A). Thus, the ARFGAP1 localizing to the p62 compartment is not influenced by dispersal of the Golgi and does not colocalize with other Golgi markers, implying a non-Golgi origin for these structures.

**Figure 6.**
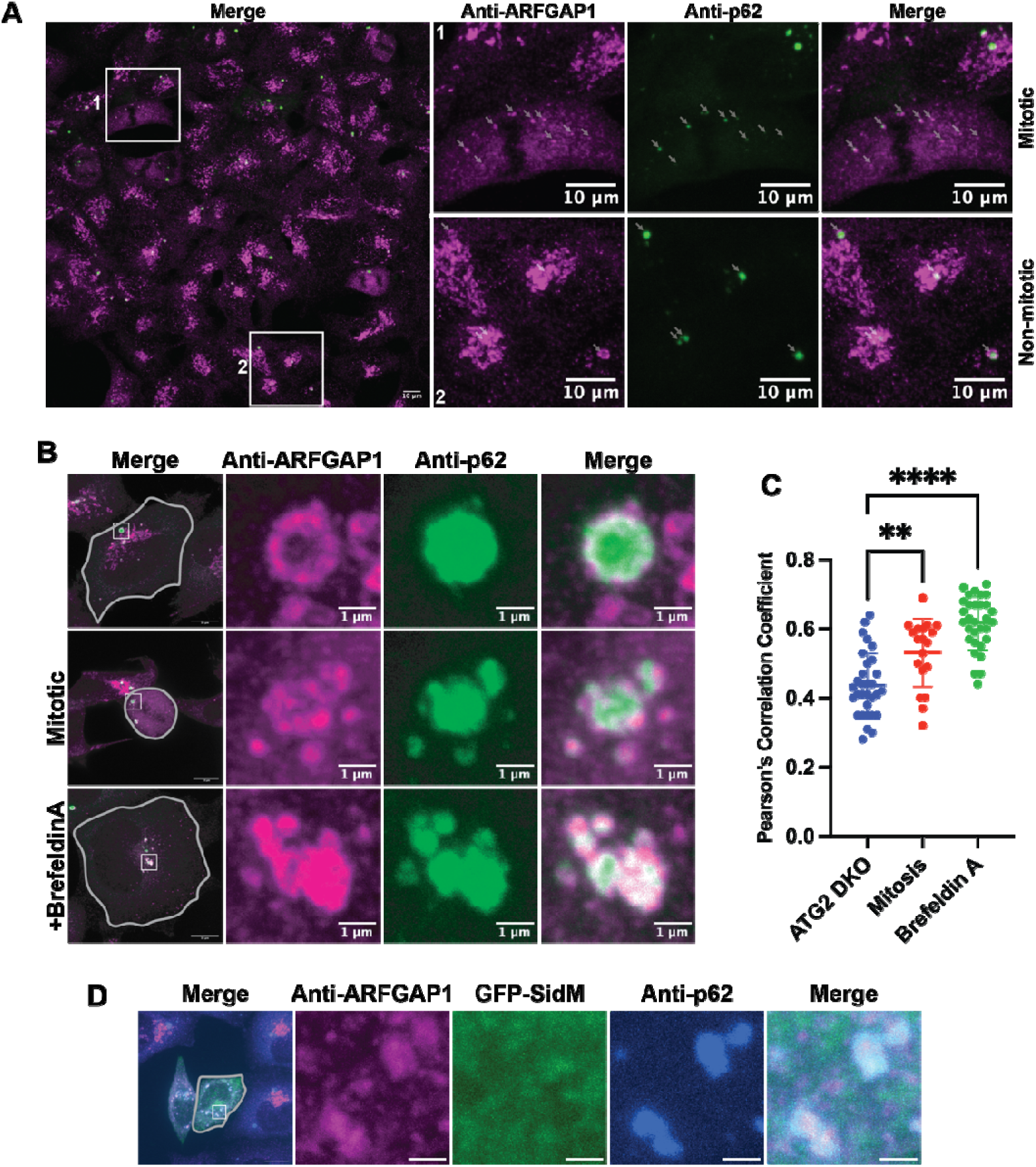
ARFGAP1 accumulation at the ATG2 DKO compartment is independent of the Golgi. **(A)** Immunofluorescence microscopy demonstrates that ARFGAP1 colocalizes with p62 in ATG2 DKO cells. Inset 1 highlights a cell undergoing mitosis, in which the colocalization of these two markers persists as denoted by the arrows. Inset 2 shows several cells, a ring-like accumulation of ARFGAP1 around p62 is clearly visible, separate from the Golgi. Maximum intensity projection of a confocal image. **(B-C)** Immunofluorescence images showing that perturbations which disrupt Golgi formation don’t affect the recruitment of ARFGAP1 to the ATG2 DKO compartment. HEK293 ATG2 DKO cells were treated or not treated with 5 μg/ml Brefeldin A for one hour, following which they were immunostained against ARFGAP1 and p62. The gray lines show the periphery of the cell and the insets focus on ATG2 DKO compartments as seen by the p62 accumulations. Colocalization between ARFGAP1 and p62 was assessed in (C) using the Pearson’s Correlation Coefficient. In the untreated sample, cells were separated into two groups: mitotic and non-mitotic, based on the ARFGAP1 fluorescence pattern. Statistical significance was assessed by one way ANOVA. *, adjusted P value <0.05. **, adjusted P value <0.01. ***, adjusted P value <0.001. ****, adjusted P value <0.0001. **(D)** Immunofluorescence image demonstrating that Golgi disruption by SidM/DrrA overexpression did not perturb the colocalization between ARFGAP1 and p62. HeLa ATG2 DKO cells were transfected with GFP-SidM and then immunostained against ARFGAP1 and p62. The gray line shows the periphery of the cell and the inset focuses on the largest accumulation of p62.

Finally, we tested whether the GAP domain of ARFGAP1 or the ALPS motif are necessary for its localization to the PAS, by comparing the colocalization of various tagged constructs with GFP-ATG2A and TagBFP-LC3B (Fig. S6A). Disruption of the GAP domain via the well characterized R50K mutation had no impact on its localization to the ATG2/LC3B-positive phagophores (Fig. S6B). Mutation of the ALPS domain (L207D, V279D) altered ARFGAP1 distributions in the cell such that specific localizations could no longer be quantified (the redistribution to the cytoplasm created a high background that was difficult to properly threshold). Nonetheless, even in this condition there remained clear accumulations of this construct at phagophores indicating that recruitment is not strongly dependent upon the ALPS motif. As these elements of ARFGAP1 are essential to its Golgi function, these results further suggest that the ARFGAP1/RAB1A membranes involved in autophagosome biogenesis derive from early secretory membranes prior to the Golgi.

## Discussion

Here we describe an interaction of ARFGAP1/RAB1-positive membranes with the autophagic biogenesis machinery including the lipid transporter ATG2. These membranes likely derive from a privileged region of the ERGIC consistent with the literature tying these intermediate secretory membranes to autophagy^17, 21^. In particular, the presence of both ARFGAP1 and RAB1 plus the general proximity to ERES proteins across a variety of assays is consistent with an early secretory origin for these membranes. Intriguingly, ARFGAP1 and RAB1 coalesce around the periphery of the ATG2 DKO compartment exactly where we previously described complex cup-shaped membranes, so called proto-phagophores^9^, accumulate. We do not detect any other ERES or ERGIC markers at these sites, implying proto-phagophores derive from only a subdomain of the ERGIC which we suspect is remodeled specifically for the purpose of supporting autophagy.

How do the ERGIC-derived RAB1 membranes contribute to autophagosome formation? Ypt1, the yeast homologue of RAB1A, was identified early on as a genetic regulator of autophagy^34^, and subsequently a number of studies have connected both Ypt1 and RAB1 to autophagosome biogenesis, but the precise mechanism of action remains contentious. In yeast, Ypt1 has been shown to interact with Atg1^35^ and Atg17^47^, highly suggestive of a role in regulating or recruiting the kinase machinery driving initiation of autophagosome biogenesis. Human RAB1A binds the VPS34 complex favoring conformations that likely support PI3P formation^48^. Thus, either or both of the early acting complexes in autophagosome biogenesis may engage RAB1 proteins. In this study, we find that RAB1 depletion elevates LC3B-II levels, suggesting that RAB1 acts either downstream or independently from the lipidation machinery. Further, we present two orthogonal and highly quantitative methods to follow LC3B degradation in the lysosome. Both methods unambiguously show a clear, albeit partial, genetic dependence of LC3B degradation upon RAB1 consistent with RAB1 promoting autophagy. As we detect RAB1A at the ATG2 DKO compartment, RAB1A recruitment appears to precede ATG2A recruitment to the PAS, consistent with an interaction with either the ULK1 or VPS34 complexes, but is not essential to support LC3B-II formation on early autophagy membranes.

We propose that ARFGAP1/RAB1 membranes are directly integrated into the phagophore prior to ATG2 recruitment. The fusion of vesicular sources has been proposed as a general mechanism for phagophore biogenesis^49, 50^. For example, the HyPas model postulates that FIP200 vesicles from the *cis*-Golgi fuse with endosomally derived ATG16L1 vesicles to create the nucleating structure for autophagosome formation^22^. Recent reports, including our own, have also identified ATG9 vesicles as a seed for autophagosome biogenesis, which expand through lipids delivered by ATG2^9, 10^ and then scrambled into both leaflets by ATG9^6–8^. An integration of these models is necessary. Geometric estimates of the surface-to-volume ratio in yeast autophagosomes, gleaned from cryo-electron microscopy tomograms, suggest that as much as 20% of the membrane might derive from fusion rather than lipid transport in order to deliver luminal volume^51^, which cannot be supplied by lipid transport. We therefore propose a model in which an early fusion step between ATG9A vesicles and ARFGAP1/RAB1 membranes, both of which accumulate in ATG2 DKO cells, primes the nucleating membrane for autophagosome biogenesis.

## Materials and Methods

### Plasmids and reagents

The following plasmids were used in this study: pLVX Puro GFP-ATG2A (Addgene 250863; Previously published^3^), pLVX Puro APEX2-GFP-ATG2A (Addgene 250864; This study; a PCR fragment containing flanking EcoRI cutsites around APEX2 was generated from pcDNA3 APEX2-NES (Addgene #49386) and was inserted into GFP-ATG2A by ligation), pCMV 3xFLAG-ATG2A (Addgene 250865; Previously published^3^), pLVX Neo TagBFP-LC3B (Addgene 250866; This study; pLVX puro TagBFP-LC3B^3^ and pLVX-EGFP-IRES-Neo (Addgene #128660) were cut with EcoRI and BamHI, following which the TagBFP-LC3B fragment was ligated into the pLVX Neo backbone), pDESt47 Sar1-GFP (Addgene #67409), SAR1B-mOrange2 (Addgene #166899), SAR1BH79G2-mOrange2 (Addgene #166900), SAR1BH79G-3xFLAG (Addgene 250867; This Study; A BamHI-3xFLAG-NotI fragment was generated by oligo annealing, following which it was ligated into the SAR1BH79G-mOrange2 backbone that was cut by the same restriction enzymes), SAR1B-BFP (Addgene 250872; This Study; A BamHI-BFP-NotI fragment was generated by PCR, following which it was ligated into the SAR1B-mOrange2 backbone that was cut by the same restriction enzymes), SAR1BH79G-BFP (Addgene 250873; This Study; A BamHI-BFP-NotI fragment was generated by PCR, following which it was ligated into the SAR1BH79G-mOrange2 backbone that was cut by the same restriction enzymes), mRuby-SEC23A (Addgene 36158), EGFP-SEC23A (Addgene #66609), pEGFP-SEC24A (Addgene 250868; This Study; a XhoI-SEC24A-PacI fragment was generated by PCR from HA-SEC24A which was a generous gift from the Kundu lab. The fragment was ligated into the backbone of pEGFP-SEC24D, which was cut by the same restriction enzymes), pEGFP-SEC24B (Addgene 250869; This Study; a XhoI-SEC24B-PacI fragment was generated by PCR from HA-SEC24B which was a generous gift from the Kundu lab. The fragment was ligated into the backbone of pEGFP-SEC24D, which was cut by the same restriction enzymes), EGFP-SEC24C (Addgene 250870; This Study; a AgeI-EGFP-XhoI fragment was generated by PCR and was ligated into the backbone of pEYFP-SEC24C (Addgene #66608) which was cut by the same restriction enzymes), pEGFP-SEC24D (Addgene #32678), GFP-SEC31A (Addgene #42124), GFP-SEC16A (Addgene #36155), ARFGAP1-TagRFP (This Study; ARFGAP1-TagRFP was a gift from the Rothman lab. A frame shift and a point mutation were corrected by site directed mutagenesis), ARFGAP1-GFP (Gift from the Antonny Lab), pMRX IB HaloTag7-mGFP (Addgene # 184903), pMRX No HaloTag7-mGFP-LC3B (Addgene # 184901), pLVX GFP (Addgene 250871; This Study; a fragment containing EGFP was generated from pLVX GFP-ATG2A and was ligated into pLVX puro (Clonetech, 632159) that was cut with EcoRI using Gibson Assembly), GFP-RAB1A (Gift from the Rothman Lab), SNAP-RAB1A (Gift from the Rothman Lab), pGEX2T GST-GFP-RAB1A (Addgene 250877; this study; a GFP-RAB1A fragment was generated from the GFP-RAB1A plasmid from the Rothman lab and inserted in to the pGEX2T backbone), and pGEX2T GST-GFP-GL1 (Addgene 2508740).

Antibodies for immunoblotting were used at 1:1000 and include the following: anti-ATG2A (Cell Signaling Technology, 15011S), anti-p62 (BD BioSciences, 610832), anti-GAPDH (Abcam, ab9484), anti-LC3B (Cell Signaling Technology, 3868S), anti-RB1CC1 (Thermo Fisher Scientific, 172501AP), anti-RAB1A (Cell Signaling Technology, D3X9S), anti-RAB1B (Proteintech, 17824-1-AP), anti-ARFGAP1 (Proteintech, 13571-1-AP), anti-FLAG (Sigma-Aldrich, F1804-200UG), anti-GFP (Cell Signaling Technology, 2956S), anti-mCherry (Invitrogen, PA5-34974), ECL anti-rabbit IgG horseradish peroxidase-linked (GE Healthcare, NA934V), and ECL anti-mouse IgG horseradish peroxidase-linked (GE Healthcare, NA931V).

Antibodies for immunofluorescence were used at 1:500 and include the following: anti-Biotin (Rockland, 200-301098S), anti-GFP nanobody (Massive Photonics), anti-ARFGAP1 (Proteintech, 13571-1-AP;), anti-SEC31A (BD Biosciences, 611280), anti-SEC24C (Cell Signaling Technology, 14676S), anti-MIA3 (Tango1; Sigma, HPA055922), anti-TFG (Abcam, ab156866), anti-COPI (CMIA10; made by Rothman Lab), anti-LC3B (MBL, PM036), anti-p62 (BD BioSciences, 610832), anti-SEC16A (Abcam, ab70722), anti-SEC24D (Cell Signaling Technology, 14687S), anti-GM130 (BD Biosciences, 610823), anti-ERGIC53 (Abcam, EPR6979), Alexa Fluor 405 goat anti-rabbit IgG (Thermo Fisher Scientific, A31556), Alexa Fluor 405 goat anti-mouse IgG (Thermo Fisher Scientific, A31553), Alexa Fluor 594 goat anti-rabbit IgG (Thermo Fisher Scientific, A11037), Alexa Fluor 647 goat anti-rabbit IgG (Thermo Fisher Scientific, A21245), and Alexa Fluor 647 goat anti-mouse IgG (Thermo Fisher Scientific, A21236).

The following reagents were purchased for FLASH-PAINT: Cy3B-modified DNA oligonucleotides were custom-ordered from IDT. Sodium chloride 5 M (cat: AM9759) were obtained from Ambion. Ultrapure water (cat: 10977-015) was purchased from Invitrogen. µ-Slide 8-well chambers (cat: 80807) were purchased from ibidi. Methanol (cat: 9070-05) was purchased from J.T. Baker. Glycerol (cat: G5516-500ml), protocatechuate 3,4-dioxygenase pseudomonas (PCD) (cat: P8279), 3,4-dihydroxybenzoic acid (PCA) (cat: 37580-25G-F) and (+−)-6-hydroxy-2,5,7,8-tetra-methylchromane-2-carboxylic acid (Trolox) (cat: 238813-5 G) were ordered from Sigma. 1× Phosphate Buffered Saline (PBS) pH 7.2 (cat: 10010-023) was purchased from Gibco. Paraformaldehyde (cat: 15710) were obtained from Electron Microscopy Sciences. Bovine serum albumin (cat: 001-000-162) was ordered from Jackson ImmunoResearch. Triton X-100 (cat: T8787-50ML) was purchased from Sigma-Aldrich. DNA labeled Nanobodies were obtained from Massive Photonics.

SiControl, FIP200, RAB1A, RAB1B, and ARFGAP1 were acquired from Horizon Discovery (D-001810-10-05, L-0082383-00-0005, L-008958-01-0005, L-013321-02-005; Horizon Discovery).

### Cell culture

HEK293 cells (ATCC, CRL_1573), H4 (CVCL_1239), and HeLa (CVCL_0030) cells were cultured at 37°C and 5% CO2 in DMEM (Thermo Fisher Scientific, 11965092) supplemented with 10% FBS (Thermo Fisher Scientific, 10438062) and 1% penicillin-streptomycin (Thermo Fisher Scientific, 15140122). Cells were starved by incubation in Earle’s Balanced Salt Solution (Thermo Fisher Scientific, 24010043) for biochemical assays or in Hank’s Balanced Salt Solution (Thermo Fisher Scientific, 14025092) for live cell imaging assays and treated with 0.1 µM bafilomycin A1 (Enzo, BML-CM110-0100) for 2 h.

### Lentivirus production and transduction

HEK293 cells were seeded into a 10-cm plate. At 70% confluence, cells were transfected with 4.3 µg psPAX2 (Addgene #12260), 0.43 µg pCMV-VSV-G (Addgene, #8454), and 4.3 µg target plasmid using 20 μl Lipofectamine 3000 (Thermo Fisher Scientific, L3000008). DNA was added to 500 μl Opti-MEM (Thermo Fisher Scientific, 31985070) and Lipofectamine 3000 to 500 μl Opti-MEM in separate 1.5 ml microcentrifuge tubes. The tubes were then mixed and left for 10 min before adding the mixture dropwise to cells in fresh DMEM. After overnight incubation at 37°C, the medium was replaced with fresh DMEM. Cell medium was then collected after 24-48 h and filtered with a 0.45-µm syringe filter (Pall, 4184). Generated virus was either used immediately or was aliquoted and stored at −80°C. HEK293 cells to be transduced were seeded into a 6-well plate. Undiluted virus was added to cells and incubated at 37°C for 24 h. The medium was replaced with fresh DMEM and left for another 24 h. The cells were then treated with 2 µg/ml puromycin (Clontech, 631306), 2 mg/ml Geneticin (Thermo Fisher Scientific, 10131035), or 10 µg/ml Blasticidin (Gibco, A11139-03) for at least 2 wk. For the case of pMRX No HaloTag7-mGFP-LC3B, which lacked a selection marker, the cells were transduced and subsequently sorted by flow assisted cell sorting to normalize GFP expression levels.

### Proximity labeling

Proximity labeling was performed as previously described^27^. ATG2 DKO cells stably expressing APEX2-EGFP-ATG2A were cultured for two weeks in media containing heavy, medium, or light isotopes of arginine and lysine (Cambridge Isotope Laboratories, CLM-2265-H-PK, DLM-2640-PK, CNLM-291-H-PK; Sigma-Aldrich, 608033). All conditions were expanded to 3×15cm plates prior to proximity labeling. Heavy labeled cells were starved in EBSS and treated with 100 nM Bafilomycin A1 (Enzo, BML-CM110-0100) for 2 hrs prior to labeling. All conditions were incubated with 500 μM Biotin Phenol (Biotinyl Tyramide; Sigma-Aldrich, SML2135) for 30 min at 37°C. The heavy and medium isotope conditions were treated with 1 mM H2O2 (Sigma-Aldrich, H1009-100ML) for 1 min at room temperature. All conditions were then aspirated and scraped in quencher solution containing 10 mM sodium azide (Sigma-Aldrich, S2002-100G), 10 mM sodium ascorbate (Sigma-Aldrich, A4034-100G) and 5 mM Trolox (Sigma-Aldrich, 238813-5G) in DPBS. The cells were then spun down at 77xg for 5 min at 4°C and were resuspended in quencher solution. This process was repeated for a total of three washes. The cells were then lysed in RIPA buffer (50 mM Tris, 150 mM NaCl, 0.1% SDS, 0.5% sodium deoxycholate (Sigma-Aldrich, D6750-10G), 1% NP40 (Roche, 11754599001) and 1x protease inhibitor (Roche, 11836170001)) for 30 min prior to centrifugation for 10 min at 18,407xg at 4°C. The supernatants were collected and protein concentration was assessed via Bradford assay. Heavy, medium and light cells were mixed at a 1:1:1 ratio for a combined mass of 3-4 mg of protein. The protein mixtures were subsequently incubated with streptavidin magnetic beads (Pierce, 88817) for 2 hrs at 4°C after which they were washed twice with RIPA lysis buffer, once with 1 M KCl, once with 0.1 M Na_2_CO_3_, once with 2 M urea in 10 mM Tris (pH 8.0) and twice with RIPA lysis buffer for each sample. All washes were performed at 4°C. The proteins were eluted off of the beads by boiling the samples in 2x LDS sample buffer (Invitrogen, NP0007). The eluted fraction was run on a gel (Invitrogen, NP0341) for a 10 minutes. The resulting gel plug was stained (Invitrogen, 46-5034) and excised and sent to the Yale Keck facility for mass spectrometry analysis.

### Mass spectrometry

MS was performed at Yale Mass Spectrometry & Proteomics Resource of the W.M. Keck Foundation Biotechnology Resource Laboratory. Samples were analyzed in biological triplicates. The data were normalized to internal controls and total spectral counts. For differential analysis, 10E-04 was added to all values prior to log2-transformation, and an unpaired Student t-test was performed. Differentially abundant proteins were considered those with an absolute fold change > 1.5 and p <0.05.

### FLASH-PAINT Imaging

Cells were fixed with 4% PFA (Electron Microscopy Services, 15710-S) for 30 min. After three washes in 1x PBS for five minutes each, cells were blocked and permeabilized with 3% BSA and 0.25% Triton X-100 at room temperature for 1 h. Next, cells were incubated with the anti-ARFGAP1 antibody, anti-SEC31A antibody, and the GFP-Nanobody in 3% BSA and 0.1% Triton X-100 at 4 °C overnight. Additionally, all other primary antibodies were pre incubated with the corresponding nanobodies (Tables S1) at 4 °C overnight. The next day, after four washes (30 s, 60 s, 2× 5 min) cells were incubated with the nanobodies corresponding to ARFGAP1 antibody and SEC31A antibody for ∼2 h at room temperature. Next, unlabeled excess secondary nanobodies (to block unlabeled epitopes) were added to pre-incubation antibody-nanobody mixes at room temperature for 5 min. Next, the cells were incubated with the pooled antibody-nanobody mix for ∼2.5 h at room temperature. After four washes (30 s, 60 s, 2× 5 min), the sample was post-fixed with 3% PFA and 0.1% GA for 10 min. Finally, samples were washed three times with 1× PBS for 5 min each before adding the imaging solution.

The imaging solution consisted of 1x PBS and 500 mM NaCl and was supplemented with 1× Trolox, (100× Trolox: 100 mg Trolox, 430 μL 100 % Methanol, 345 μL 1M NaOH in 3.2 mL H_2_O), 1× PCA (40× PCA: 154 mg PCA, 10 mL water and NaOH were mixed and pH was adjusted 9.0), and 1× PCD (100× PCD: 9.3 mg PCD, 13.3 mL of buffer (100 mM Tris-HCl pH 8, 50 mM KCl, 1 mM EDTA, 50 % Glycerol)). All three stock buffers were frozen and stored at -20 °C.

Fluorescence imaging was carried out on an inverted Nikon Eclipse Ti2 microscope (Nikon Instruments) with the Perfect Focus System, attached to a Andor Dragonfly unit. The Dragonfly was used in the BTIRF mode, applying an objective-type TIRF configuration with an oil-immersion objective (Nikon Instruments, Apo SR TIRF 60×, NA 1.49, Oil). As excitation laser, a 561 nm (1W nominal) was used. The beam was coupled into a multimode fiber going through the Andor Borealis unit reshaping the beam from a gaussian profile to a homogenous flat top. As dichroic mirror a CR-DFLY-DMQD-01 was used. Fluorescence light was spectrally filtered with an emission filter (TR-DFLY-F600-050) and imaged on a scientific complementary metal oxide semiconductor (sCMOS) camera (Sona 4BV6X, Andor Technologies) without further magnification, resulting in an effective pixel size of 108 nm. The power at the objective lens was ∼10 % of the power set at the laser.

Imaging was carried out using the corresponding Imager and Adapter (Table S2, S3, S5) in imaging buffer. 25,000 frames were acquired at 25 ms exposure time. The readout bandwidth was set to 540 MHz. Laser power (@561 nm) was set to 80 mW (measured before the back focal plane (BFP) of the objective), corresponding to ∼1.8 kW/cm^2^ at the sample plane. After imaging the sample was subsequently washed three times with 200 µL each with 1× PBS (on the microscope) followed by an incubation with the corresponding eraser (Table S4; at 20 nM) for ∼ 5 min. This process was repeated iteratively for ten sequential rounds (Table S5).

DNA-PAINT data was reconstructed, postprocessed (drift correction and alignment of imaging rounds) and rendered with the Picasso package^52^.

### Focused ion beam scanning electron microscopy and correlative light electron microscopy

For focused ion beam scanning electron microscopy (FIBSEM) CLEM, ATG2 DKO HEK293 cells overexpressing GFP-ATG2A were plated on 35-mm MatTek dish (MatTek, P35G-1.5-14-CGRD). The cells were starved for two hours prior to pre-fixation in 4% PFA + 0.25% glutaraldehyde and three washes in 1x PBS. Regions of interest by fluorescence light microscopy were selected and their coordinates on the dish were identified using phase contrast. Cells were further fixed with 2.5% glutaraldehyde in 0.1 M sodium cacodylate buffer, postfixed in 2% OsO4 and 1.5% K4Fe(CN)6 (Sigma-Aldrich) in 0.1 M sodium cacodylate buffer, en bloc stained with 2% aqueous uranyl acetate, dehydrated, embedded in Embed 812, and polymerized at 60°C for 48 hrs. Cells of interest were relocated based on the pre-recorded coordinates. Epon block was glued onto the SEM sample mounting aluminum stub and platinum en bloc coating on the sample surface was carried out with the sputter coater (Ted Pella, Inc.). The cell of interest was relocated under SEM imaging based on pre-recorded coordinates and FIB-SEM imaged in a Crossbeam 550 FIB-SEM workstation operating under SmartSEM (Carl Zeiss Microscopy GmbH) and Atlas 5 engine (Fibics, Inc.). The imaging resolution was set at 7 nm/pixel in the X, Y axis with milling being performed at 7 nm/step along the Z axis to achieve an isotropic resolution of 7 nm/voxel. Images were aligned and exported with Atlas 5 (Fibics, Inc.), further processed and segmented with DragonFly Pro software (Object Research Systems [ORS], Inc.).

### Expi293 culture and transfection

3xFLAG-ATG2A was expressed and purified from Expi293 suspension cells. Cells were cultured at 37°C, 8% CO2 in vented PETG flasks (Thermo Scientific, Nalgene; 4115-0500) with 125 rpm rotation. Prior to transfection, cells were grown to 3-5 x 106 cells/mL then diluted to 2.5×106 cells/mL and allowed to grow for 18-24 hours. Upon reaching 4.5-5.5 x 106 cells/mL, the culture was diluted to 3 x 106 cells/mL with fresh media. Cells were transfected with 100 μg plasmid per 100 mL of cultured cells. For every 100 mL of cells, 100 μg DNA was diluted in 5 mL OptiMEM. 300 µL of 1 mg/mL polyethylenimine linear 25k (Polyscience, 23966) was diluted into 5 mL OptiMEM in a separate tube. Both tubes were incubated for 5 minutes at room temperature then combined and further incubated for 30 minutes. The DNA/PEI mixture was then added dropwise into Expi293 cultures. 12-18 hours after transfection, protein expression was enhanced using 700 μL of 500 mM valproic acid. Cells were then harvested after 48-72 hours.

### 3xFLAG-ATG2A purification

Cell pellets were resuspended in 10 mL purification buffer A (500 mM NaCl, 20 mM HEPES, pH: 8.0, 10% glycerol, 1 mM TCEP) for every 100 mL of original culture volume. Purification buffer A was supplemented with 1x EDTA-free protease inhibitor cocktail (Roche; 11873580001). Cells were then lysed by subjecting pellet resuspensions to 6 freeze/thaw cycles in liquid nitrogen The lysates were then clarified by centrifugation at 18,500 x G, and supernatants were transferred to 150 μL of anti-Flag M2 agarose resin per 100 mL of initial culture volume (prewashed with 10 mL purification buffer A). Lysates were incubated with beads under mild agitation for 2 hours at 4 °C. Lysates and beads were then passed through a gravity filtration column and washed with 4 CVs of purification buffer. Beads were then incubated with 2.5 mM ATP and 5 mM Mg2+ in purification buffer overnight at 4 °C. Columns were then washed with 2 CVs of purification buffer, followed by elution with 50 μL of 1 mg/mL 3x-Flag (DYKDDDDK) peptide (ApexBio Technology; A6001) per 100 mL of initial culture volume. Protein concentration was determined against a BSA standard curve by Coomassie staining prior to subsequent assays. Proteins were flash frozen in liquid nitrogen for storage at -80 °C.

### Recombinant protein expression and purification from *E. Coli*

For all in vitro experiments, GFP-GL1 refers to protein expressed and purified from constructs ending at the reactive glycine (G116) of GL1, as this construct was previously used for lipidation experiments in the absence of ATG4 proteins. In summary, human GFP-GL1 and GFP-RAB1A were cloned in the pGEX-2T GST vector. The proteins were expressed in BL21-Gold (DE3) competent cells (Agilent Technologies; 230132). The cells were cultured in 4 liters of Luria Bertani broth (LB) medium with 50 mg/ml carbenicillin (C1389; Sigma-Aldrich) and induced with 0.5 mM isopropyl β-D-thiogalactopyranoside (IPTG). Bacterial pellets were treated with EDTA-free protease inhibitor tablets in thrombin buffer (20 mM Tris pH 7.5, 100 mM NaCl, 5 mM MgCl_2_, 2 mM CaCl_2_, 1 mM DTT). The cells were broken in a cell disrupter, and the lysate was incubated with pre-washed glutathione agarose beads (G4501; Sigma-Aldrich) for 3 h at 4°C. The beads were washed three times and then incubated with thrombin (10 μl of thrombin [T6884; Sigma-Aldrich], 500 μl of thrombin buffer, 0.5 μl of DTT, 500 μl of beads) to cut the proteins from GST tags overnight at 4°C. Purified proteins were stored in 20% glycerol at −80°C.

### Lysis, gel electrophoresis, and immunoblotting

Cells were collected by scraping in PBS. The cells were centrifuged at 500 g for 3 min at 4°C, and lysed in lysis buffer (10 mM Tris pH 7.5, 150 mM NaCl, 0.5 mM EDTA, 0.5% NP-40) with EDTA-free protease inhibitor cocktail (Sigma-Aldrich, 11873580001) on ice for five minutes. Lysates were collected after centrifugation at 16,000 g for 10 min at 4°C. Total protein concentration was determined by a Bradford assay (BioRad, 5000006). 50-100 μM DTT (American Bio, AB00490-00010) and 1x LDS (Invitrogen, NP0007) loading buffer were added to each sample, which were then electrophoresed on precast Bis-Tris gels (Thermo Fisher Scientific, NP0341BOX or NP0322BOX). Gels were subsequently transferred to Immobilon-FL PVDF membranes (Sigma-Aldrich, IPFL00010) at 30-35 V for 90 min.

The membranes were blocked with 5% BSA (Sigma-Aldrich, A9647) in PBS-T (PBS containing 0.1% Tween 20; Thermo Scientific, CAS 9005-64-5) for 1 h at room temperature and incubated with primary antibody (diluted 1:1,000 in PBS-T containing 5% BSA [Sigma-Aldrich, A9647] and 0.03% sodium azide (Sigma-Aldrich, S2002)) overnight at 4°C. Membranes were washed three times in PBS-T before incubation with the secondary antibody which was diluted 1:5,000 in PBS-T containing 0.5% Omniblock (American Bio, AB10109-00100) for 1 h. Membranes were then washed three times in PBST and treated with SuperSignal West Femto substrate (Thermo Fisher Scientific, 34096) for 5 min before imaging with the Bio-Rad VersaDoc imaging system.

### Immunoprecipitation

For immunoprecipitation experiments, 15 cm plates were seeded and transfected upon reaching 80-90% confluency. The next day, the cells were collected by scraping, washed 1x in PBS, and lysed as previously described. Following quantification of protein levels by a Bradford assay, equal amounts of protein were added to pre-washed GFP-Trap beads (Proteintech, gtma-20). The volume was increased to 500 uL and the bead/lysate slurry was incubated at 4°C for two hours while rotating. The beads were pelleted using a magnetic rack and washed five times in lysis buffer (10 mM Tris pH 7.5, 150 mM NaCl, 0.5 mM EDTA, 0.5% NP-40) with EDTA-free protease inhibitor. The target proteins were eluted by boiling the beads for 10 min in 2x LDS loading buffer.

### Densitometry and quantification

Densitometry quantifications of immunoblots and in-gel fluorescence images were performed using ImageJ software. For LC3B and p62 clearance analysis (Fig. S1 A-C), the band intensity of LC3B or p62 for each condition was normalized against the GAPDH intensity of that lane. To normalize each replicate, each value was divided by the mean value of the entire replicate. This normalization technique was also employed in the co-IP experiments. Statistical significance was determined by two-way ANOVA followed by a Tukey multiple comparison test to compare cell types. To quantify the co-IP reactions, the intensity of the 3xFLAG-ATG2A IP band was ratioed against the intensity of its corresponding input band. Following normalization, statistical significance was determined by a one (Fig. S3 A-D) or two-way (Fig. 2E-F) ANOVA followed by a Dunnett’s multiple-comparisons test comparing each sample with the control lane. For the HaloTag flux assay (Fig. 2A, 2B, 3A, 3B), the Halotag flux rate was calculated as the ratio of the free Halotag band divided by the unprocessed Halotag band. Statistical significance was determined by a one (Fig. 2 A and B) or two-way (Fig 3 A and B) ANOVA followed by a Dunnett multiple comparison test to comparing each sample to the control lane. For one way ANOVA tests, parametric tests were used for all comparisons. For two way ANOVA tests, the Geisser-Greenhouse correction was used as equal variability was not assumed. All data are *n* = 3 or greater and represent biological replicates. All data were plotted with mean ± SD, and all significance values represent adjusted P value based on grouped analyses in Prism 10 (GraphPad). Asterisks indicate significance: *, P < 0.05; **, P < 0.01; ***, P < 0.001; ****, P < 0.0001. All blots were adjusted in contrast for better visualization of the bands in the figures.

### LC3B flux analysis

WT, ATG2 DKO and ATG2 DKO + APEX2-GFP-ATG2A HEK293 cells were cultured in 6-well plates to 70-90% confluence. At the beginning of the assay, the media was replaced with fresh DMEM, DMEM containing 100 nM bafilomycin A1, or EBSS containing 100 nM bafilomycin A1. After a 2 hour incubation, cells were harvested by scraping, washed with 1x PBS, and lysed. 20 μg of protein from each sample were analyzed by SDS-PAGE and immunoblotting for ATG2A, p62, GAPDH, and LC3B.

### HaloTag flux assay

HEK293 WT stably expressing either HaloTag7-mGFP or HaloTag7-mGFP-LC3B were cultured in 6-well plates to 60-70% confluence prior to transfection with either SAR1B/SAR1BH79G-mOrange2 for one day (Fig. 3A) or siRNA for two days (Fig. 2A). SAR1B transfection was performed using Lipofectamine 3000 (Thermofisher Scientific, L3000015) according to the manufacturer’s protocol. SiRNA transfection was performed using Dharmafect (Horizon Discovery, T-2001-01) according to the manufacturer’s protocol. The cells were incubated with 100 nm TMR-conjugated HaloTag ligand (Promega, G8251) for 30 min. Following two washes with 1x PBS, the cells were starved in EBSS for four hours. The cells were then collected and lysed. Following SDS-PAGE separation, in gel fluorescence was immediately measured with a ChemiDoc MP Imaging System (BioRad). Images were taken at 30 s and 100 s exposures. The gel was then transferred and immunoblotted against mCherry and GAPDH (Fig 3A) or FIP200, RAB1A, RAB1B, ARFGAP1, GAPDH, and LC3B (Fig 2A).

### GFP-mCherry-LC3B Flux analysis by FACS

H4 cells and H4 cells with a GFP-mCherry-LC3B heterozygous knock-in tag were seeded in a six well plate. Following a three day siRNA knock down, the cells were subject to a four hour starvation period in HBSS. They were subsequently trypsinized, pelleted, resuspended in HBSS, and passed through a filter prior to FACS data acquisition. The cell population was gated to ensure a single, live cell population. The ratio between the GFP and mCherry intensity was plotted as a histogram using FlowJo 10 and the ratio between the average values of each replicate was plotted using Prism 10. Three biological replicates were combined and statistically assessed using a one-way ANOVA followed by a Dunnett multiple comparison test. All data were plotted with mean ± SD, and all significance values represented adjusted P value based on grouped analyses in Prism 10 (GraphPad). Asterisks indicate significance: *, P < 0.05; **, P < 0.01; ***, P < 0.001; ****, P < 0.0001.

### Immunofluorescence

Cells seeded on coverslips were fixed in 4% PFA (Electron Microscopy Sciences, 15710), washed three times with PBS, and permeabilized in PBS containing 0.1% Saponin (Sigma-Aldrich, 47036) and 3% BSA. Incubation with primary antibodies was done at 4C for 18-24 hrs and secondary antibodies at room temperature for 1 hr at indicated concentrations (see Reagents). After each antibody incubation, cells were washed three times in PBS containing 0.1% Saponin and 3% BSA. The coverslips were mounted on precleaned microscope slides with Fluoromont-G mounting reagent (Southern Biotech, 0100-01). Images were acquired on an inverted Zeiss LSM 880 laser scanning confocal microscope with AiryScan, using Zen acquisition software (Fig S1D) or a Nikon CSU-W1 SoRa 60x oil objective, using the Nikon Elements software (Fig. 1C, 3C-F, S3A, S3C, 4A, 4C, 4E, 4F). All images were processed using ImageJ software. Images are displayed either as maximum intensity projections or single confocal slices as stated in the figure legends.

To assess the enrichment factor of early secretory proteins around the ATG2 DKO compartment (Fig. 4D), the phagophore channel which was denoted by either LC3B or p62 was masked and duplicated. Following outlier removal, one masked image was dilated and the other eroded. Following an XOR function, the resulting signal formed a ring around the ATG2 DKO compartment and the early secretory protein channel(s) were measured using this ROI. Separately, the early secretory protein channel was masked using a manual threshold and measured. The enrichment factor was calculated as (Mean_compartment_-Min_cell_)/(Mean_cell_-Min_cell_). Three biological replicates were combined and statistically assessed using a one-way ANOVA followed by a Dunnett multiple comparison test. All data were plotted with mean ± SD, and all significance represent considering adjusted P value based on grouped analyses in Prism 10 (GraphPad). Asterisks indicate significance: *, P < 0.05; **, P < 0.01; ***, P < 0.001; ****, P < 0.0001.

### Live cell imaging

Cells were plated on poly-d-lysine coated MatTek dishes (MatTek, P35GC-1.5-14-C). The following day the cells were transfected using Lipofectamine 3000 (Thermo Fisher Scientific, L3000015) according to the manufacturer’s protocol for 18-24 hrs. The cells were starved in HBSS (Thermo Fisher Scientific, 14025092) for 2-4 h and imaged in the same solution at 37°C. All images were analyzed using ImageJ and are displayed as maximum intensity projections.

For phagophore quantification (Fig. 1 C and D), each channel was masked using a manual threshold and outliers were removed. The ATG2A and LC3B channels were merged using the AND function and quantified as the total number of phagophores. The phagophore channel was then merged using the AND function with the transfected gene and quantified as the number of positive phagophores. Three biological replicates were combined and the ratio of positive over total was plotted and statistically assessed using a one-way ANOVA followed by a Dunnett multiple comparison test. All data were plotted with mean ± SD, and all significance values represent adjusted P value based on grouped analyses in Prism 10 (GraphPad). Asterisks indicate significance: *, P < 0.05; **, P < 0.01; ***, P < 0.001; ****, P < 0.0001.

## Data Availability

The data, code, protocols, and key lab materials used and generated in this study are listed in a Key Resource Table alongside their persistent identifiers at 10.5281/zenodo.20213838.

## Acknowledgments

This work was supported by grants from the National Institutes of Health (R01 GM100930 and R35 GM153482 to TJM; R01 GM151829 to JB; DA018343 to PDC), F31 AG079606 to DMF and F31 DK136246 to JLK. This research was also funded in part through Aligning Science Across Parkinson’s (ASAP-025173 to TJM and PDC) through the Michael J. Fox Foundation for Parkinson’s Research (MJFF) and the Howard Hughes Medical Institute (HHMI; PDC). FS acknowledges support from the Human Frontier Science Program (LT000056/2020-C). JB acknowledges support by the Wellcome Leap Foundation. We would like to thank Bruna Mafra de Faria, Michael Hanna, and Jeff Coleman for their support and reagent sharing. We would further like to thank Juan Bonifacino for the gift of the HeLa ATG2 DKO cells as well as the H4 GFP-mCherry-LC3B KI cells. Imaging was supported by the Yale Center for Cellular and Molecular Imaging (both the fluorescence and electron microscopy facilities). We also thank the MS & Proteomics Resource at Yale University for providing the necessary mass spectrometers and the accompanying biotechnology tools funded in part by the Yale School of Medicine and by the Office of The Director, National Institutes of Health (S10OD02365101A1, S10OD019967, and S10OD018034). The funders had no role in study design, data collection and analysis, decision to publish, or preparation of the manuscript.

## Figures and Tables

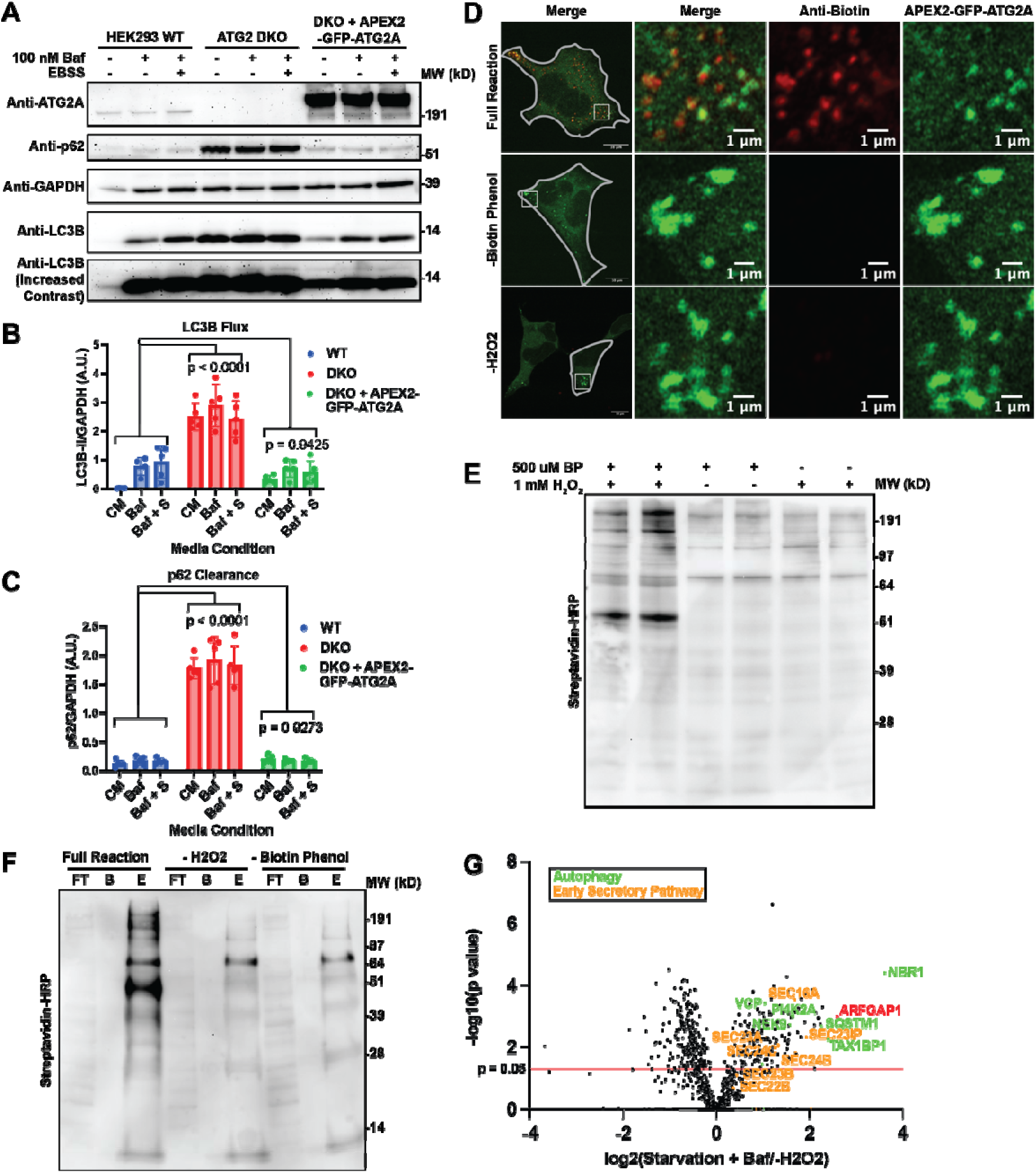

**Supplemental Figure 1.**
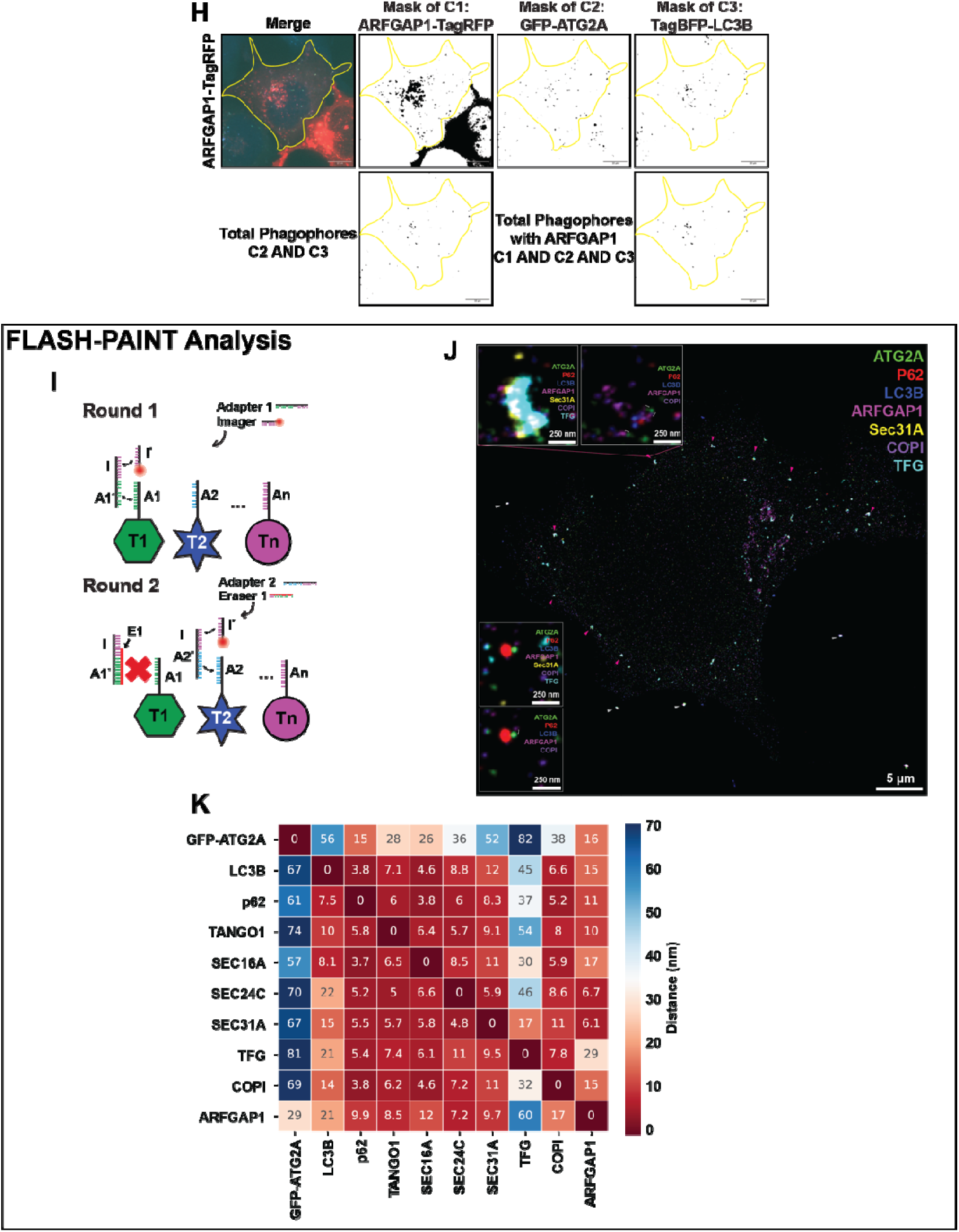
ATG2A and ARFGAP1 colocalize during starvation. **(A-C)** Immunoblot showing the restoration of LC3B flux (B) and p62 clearance (C) in ATG2 DKO cells stably expressing APEX2-GFP-ATG2A. The restoration of a wild type phenotype indicates that the fusion protein is both localizing and functioning properly. The band intensity of LC3B (B) and p62 (C) were normalized against the intensity of GAPDH for each lane. To normalize each replicate, each individual value was divided by the average of the replicate. Statistical significance was assessed by two way ANOVA. Data from five biological replicates were collected for each condition, save for the WT + Baf and WT + Baf + EBSS, which each had four replicates. **(D-F)** Single slice confocal immunofluorescence images (D) and immunoblots (E-F) demonstrating that the addition of biotin phenol and hydrogen peroxide to ATG2 DKO cells stably expressing APEX2-GFP-ATG2A results in the biotinylation of proteins proximal to ATG2A. The omittance of either biotin phenol or hydrogen peroxide resulted in the loss of the anti-biotin signal, save for endogenously biotinylated proteins. The immunoblot in (F) demonstrates that the biotinylated proteins can be enriched on streptavidin beads. **(G)** Proximity labeling of HEK293 ATG2 DKO cells stably expressing APEX2-GFP-ATG2A was performed as described in (Fig. 1A). **(H)** An example of the workflow to calculate phagophore fraction as seen in (Fig. 1D). **(I-K)** FLASH-PAINT analysis confirming the proximity of ATG2A to ARFGAP1. ATG2 DKO cells stably expressing GFP-ATG2A were labeled with all of the antibodies listed in (J), which in turn were preincubated with secondary antibodies with unique DNA sequences as depicted in (H). Each target (T1, T2… Tn) was visualized sequentially by adding an adaptor DNA strand that matched the target sequence (A1, A2… An) and an additional sequence to recruit the imager DNA strand (H). To remove the imager strand from each sequential target, a unique eraser strand was added that recognized the entirety of the adaptor sequence (e.g. A1) and part of the imager sequence, resulting in the total sequestration of the adaptor DNA strand. The subsequent adaptor sequence was then added to visualize the next target protein. Seven of the ten visualized proteins are depicted in (I), which highlights the proximity of ATG2A and ARFGAP1 at autophagic and early secretory membranes. The heatmap presented in (J) shows the median distance within 200 nm between all visualized proteins. (I); pink arrow heads = ERES. white arrow heads = exogenous beads used as a fiduciary marks in imaging. (I; insets), brackets indicate immediate proximity of ATG2 (green) and ARFGAP (maroon) as frequently observed in ERES.

**Supplemental Figure 2.**
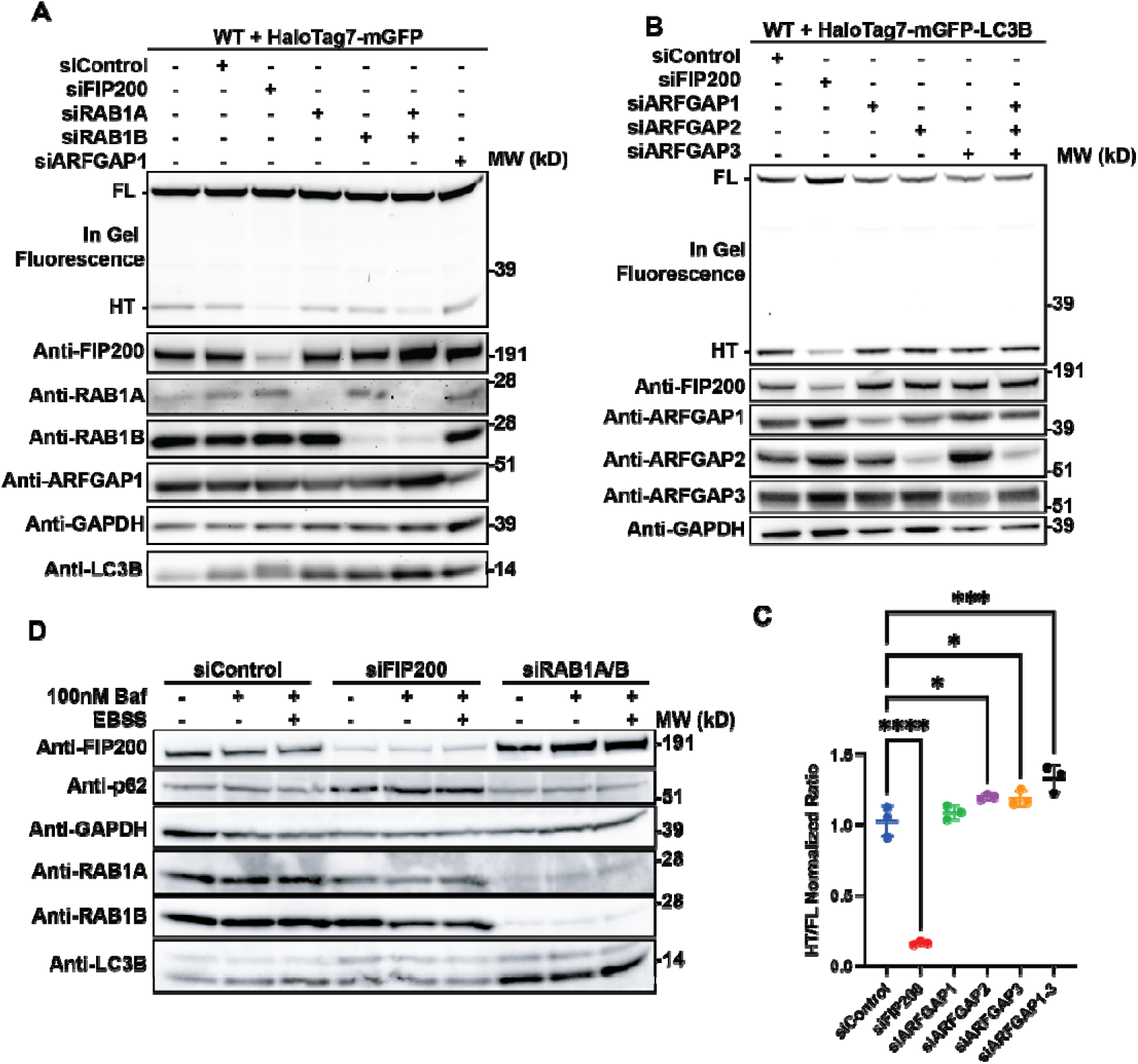
Autophagic flux depends upon RAB1 but not ARFGAP1. **(A)** In gel fluorescence image (top panel) and immunoblots demonstrating that siRNA KD of RAB1 inhibits autophagic turnover of a cytosolic reporter of bulk autophagy. In gel fluorescence reveals HaloTag7s moiety bound to the TMR ligand. Indicated bands are full length (FL) HT-mGFP protein and HaloTag7 after cleavage (HT) in the lysosome. **(B-C)** In gel fluorescence image (top panel) and immunoblots demonstrating that siRNA KD of ARFGAP1-3 slightly promotes autophagic turnover of a HaloTag7-mGFP-LC3B reporter. In gel fluorescence reveals HaloTag7s moiety bound to the TMR ligand. Indicated bands are full length (FL) HT-mGFP protein and HaloTag7 after cleavage (HT) in the lysosome. The ratio between FL and HT demonstrates autophagic flux, quantified in (C). Statistical significance was assessed by one way ANOVA. *, adjusted P value <0.05. **, adjusted P value <0.01. ***, adjusted P value <0.001. ****, adjusted P value <0.0001. Data from three biological replicates were collected for each condition**. (D)** Immunoblot showing the impact of siRAB1 knockdown on LC3B-II levels. HEK293 WT cells were not treated or treated with 100 nM bafilomycin A1 or EBSS and 100 nM bafilomycin A1.

**Supplemental Figure 3.**
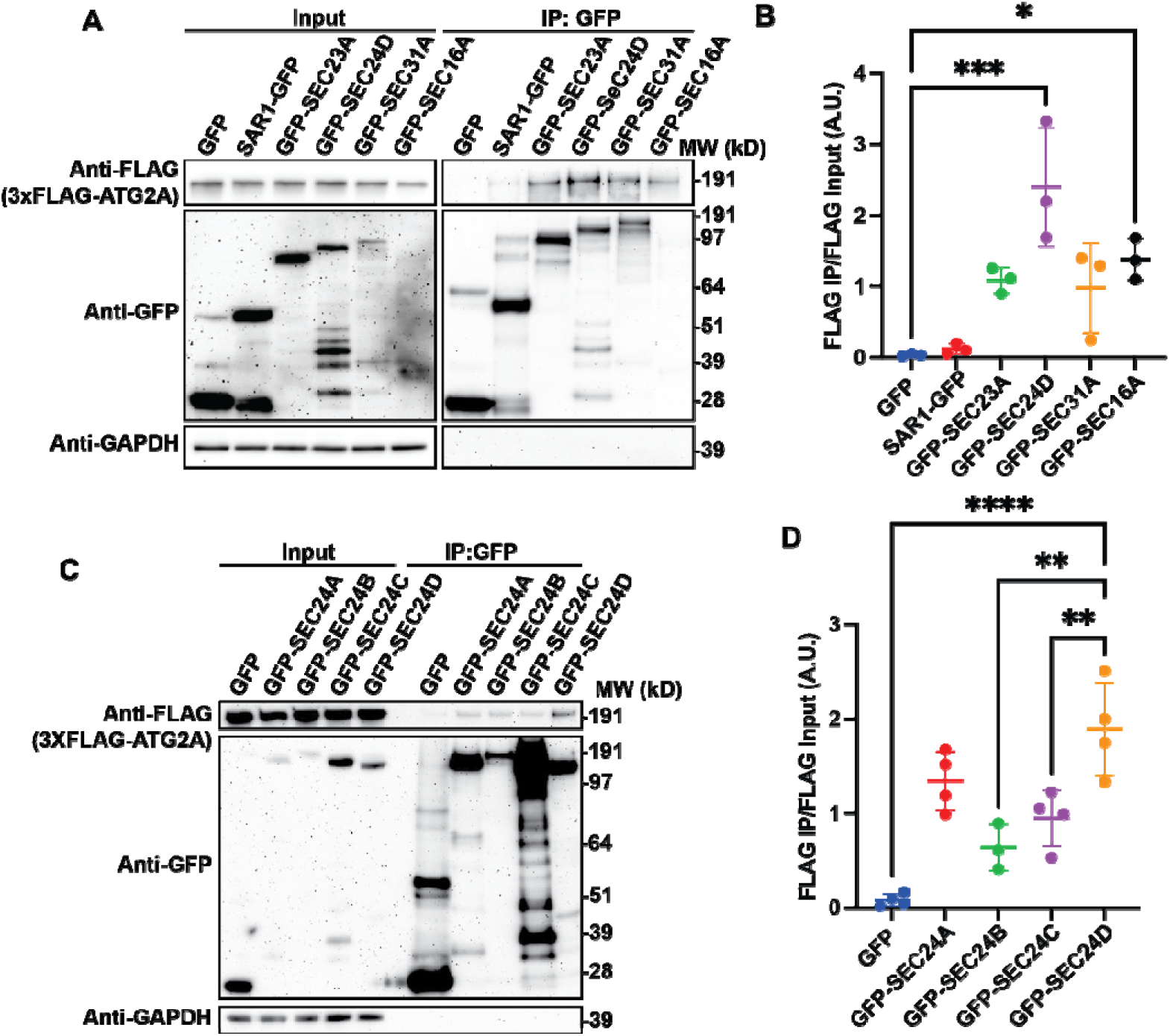
ATG2A CoIPs weakly with ER Exit Site proteins. **(A-D)** Immunoblot showing the weak CoIP between 3xFLAG-ATG2A and various ER Exit Site proteins. For the CoIP, ∼3 mg of cell lysate was used per condition (slight deviations in total amount between replicates, but not between lanes). Immunoblots quantified as in (Fig. 2B). Statistical significance was assessed by one way ANOVA. *, adjusted P value <0.05. **, adjusted P value <0.01. ***, adjusted P value <0.001. ****, adjusted P value <0.0001.

**Supplemental Figure 4.**
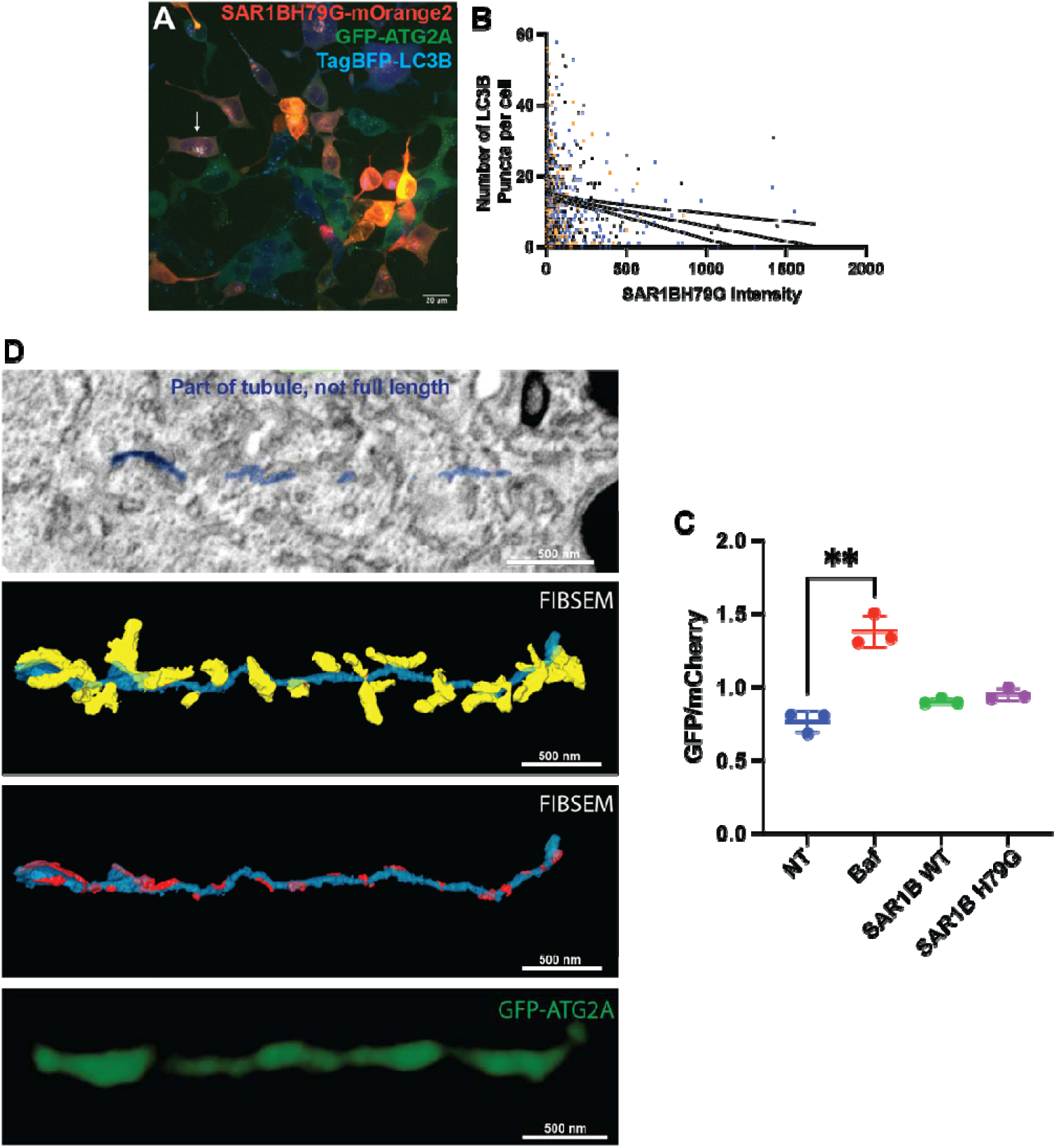
SAR1BH79G overexpression hinders autophagy and leads to the formation of ATG2A positive membrane tubules. **(A-B)** Live cell imaging shows that SAR1BH79G overexpression negatively correlates with LC3B puncta number. ATG2 DKO cells stably expressing GFP-ATG2A and BFP-LC3B were transfected with SAR1BH79G-mOrange2 and starved for 2 hours. The number of transfected cells with GFP-ATG2A tubules (white arrow in A) was counted against the total number of transfected cells, resulting in a formation rate of 1.5±0.8%. ROIs were drawn over each transfected cell, and the resulting SAR1BH79G intensity was plotted against the number LC3B puncta in (B). A linear trendline was added in black with a 95% confidence interval. **(C)** FACS data demonstrating that SAR1BH79G overexpression only results in a partial block in autophagic flux. H4 cells with a heterozygous GFP-mCherry-LC3B endogenous tag were transfected with SAR1B-mOrange2 or SAR1BH79G-mOrange2 overnight, following which they were starved in HBSS for four hours and analyzed by FACS. The ratio between GFP and mCherry is an indicator of autophagic flux, as the GFP signal is quenched in the lysosome. This ratio is displayed as averages. Statistical significance was assessed by one way ANOVA. *, adjusted P value <0.05. **, adjusted P value <0.01. ***, adjusted P value <0.001. ****, adjusted P value <0.0001. **(D)** CLEM-FIBSEM of cells prepared as in (Fig. 4C). GFP-ATG2A fluorescence corresponds to a thin tubular membrane. A single slice SEM slice is depicted in (i). Note that the segmented tubule comes in and out of the plane of this slice, so the tubule that is highlighted in turquoise is not complete. The yellow arrows are pointing at adjacent ER structures which are segmented and displayed in (ii) in yellow. At many points, these ER structures come into very close apposition with the tubule which is denoted in red. The ER structures are removed in (iii) for better visualization of the tubule. The GFP-ATG2A fluorescence signal is depicted in (iv).

**Supplemental Figure 5.**
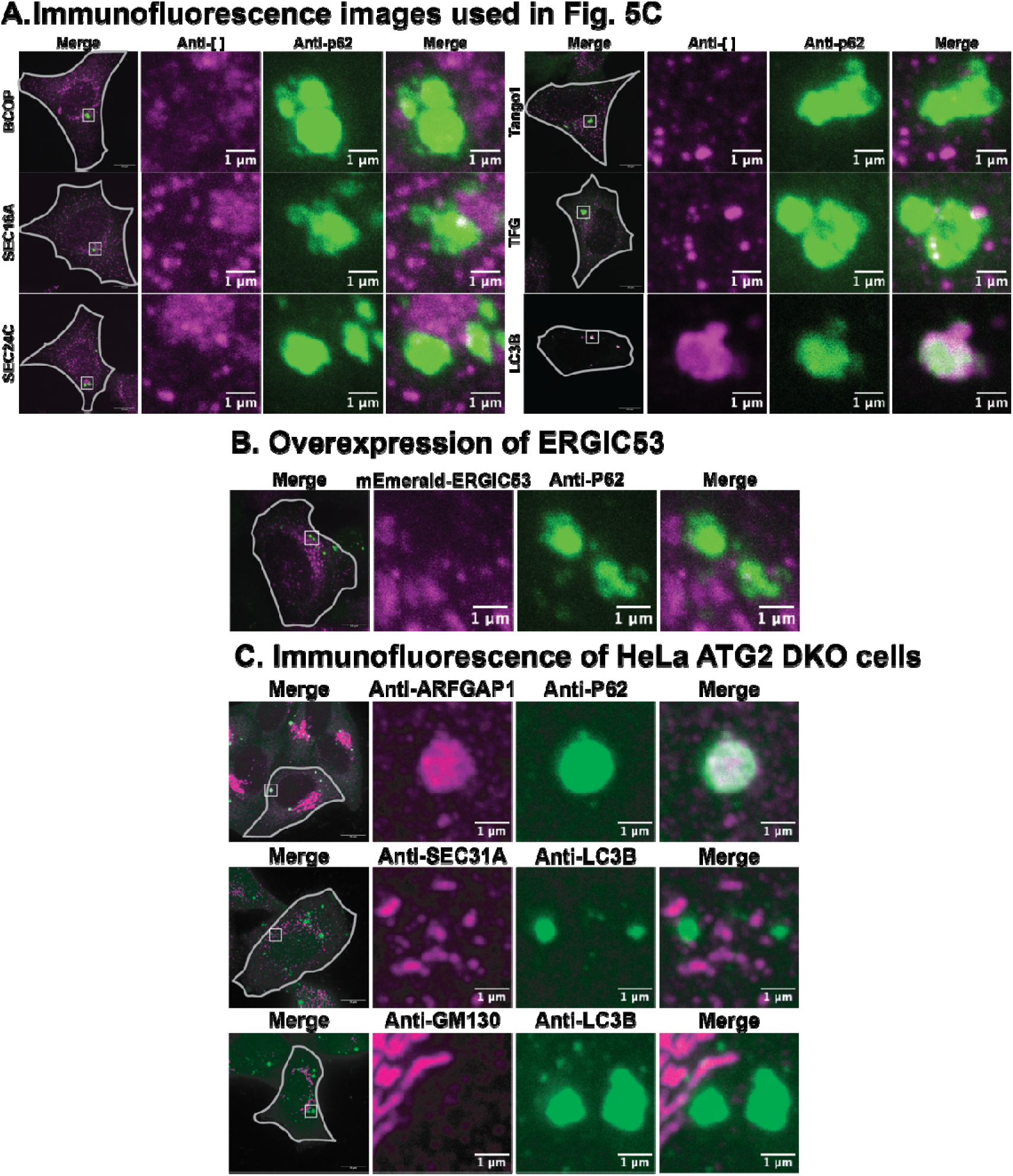
Most early secretory markers are not enriched at the periphery of the ATG2 DKO compartment. **(A-B)** Immunofluorescence images demonstrating the distribution of endogenous (A) or overexpressed (B) early secretory membranes around the ATG2 DKO compartment which is marked by p62. The images presented are single confocal slices of the center of the ATG2 DKO compartment. The gray lines show the cell periphery, and the insets focus on the largest accumulation of p62 in the cell. The images in (A) were used in the analysis of (Fig. 5C). **(C)** Immunofluorescence images in HeLa ATG2 DKO cells demonstrating a distribution of early secretory proteins as in HEK293 ATG2 DKO cells. Of note, whereas ARFGAP1 localized to the ATG2 DKO compartment in both HEK293 and HeLa cells, the distribution of ARFGAP1 was markedly different.

**Supplemental Figure 6.**
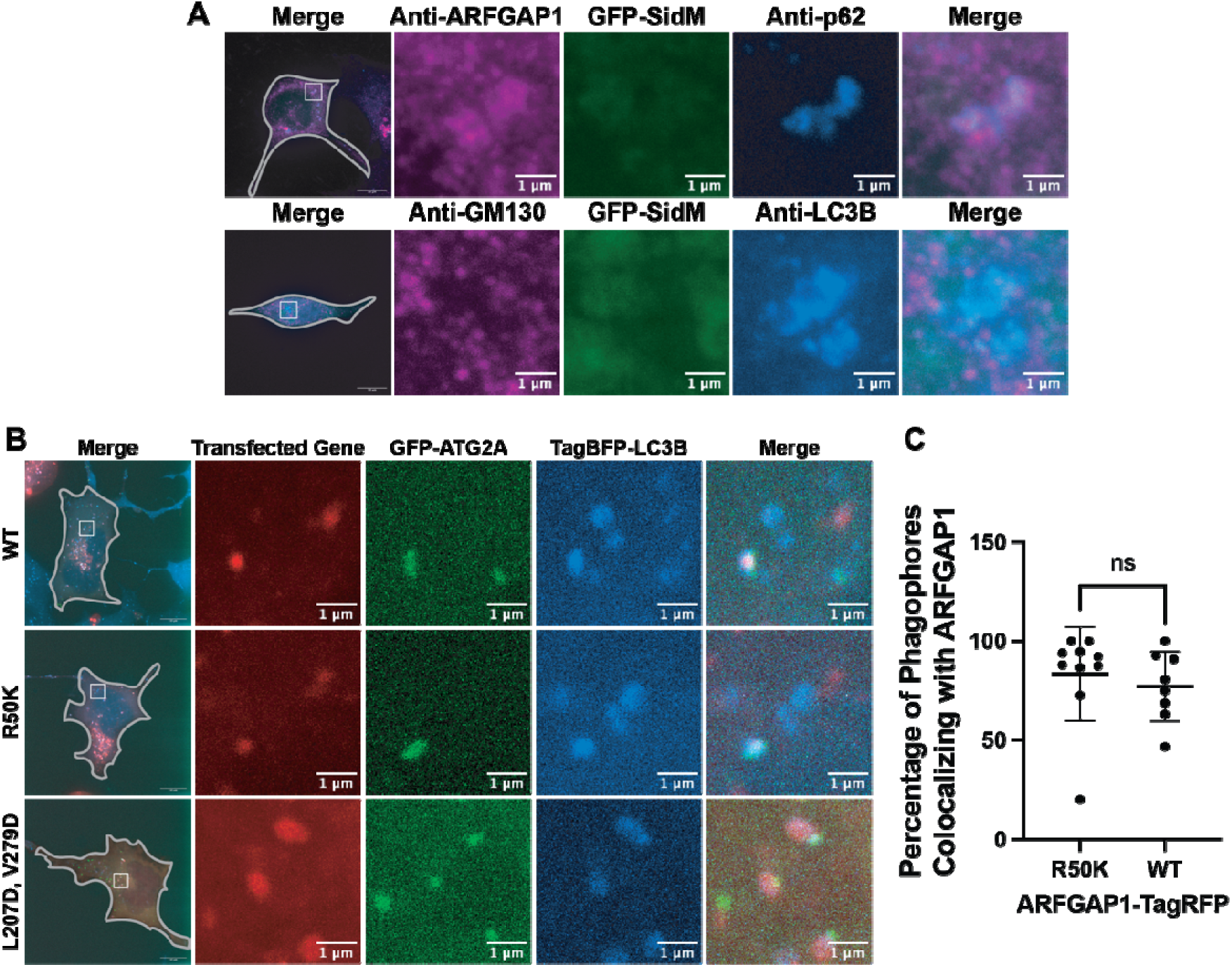
ARFGAP1 localizes to PAS independently of the Golgi, GAP activity, and ALPS motif. **(A)** Immunofluorescence images demonstrating that whereas GFP-SidM overexpression results in the complete dissipation of GM130 signal, ARFGAP1 still colocalized with the p62 signal. **(B-C)** Live cell imaging demonstrating the colocalization between ARFGAP1 mutants and phagophores, as denoted by the overlap between GFP-ATG2A and TagBFP-LC3B. Quantification for the WT and R50K construct was performed as in (Fig. 1C). Statistical significance was assessed using a two-tailed t-test.

**Table S1.** Antibody Targets and Sequences for FLASH PAINT.

| **Target** | **Nanobody** | **Sequence (3' → 5')** |
| --- | --- | --- |
| ATG2A-GFP | Anti-GFP-Nanobody-A3 | NB - TT<br>TCTTCATTAGCG |
| TFG | Anti-Rabbit-Nanobody-A15 | NB - TT<br>ATAGTGATTGGA |
| TANGO1 | Anti-Rabbit-Nanobody- A39 | NB - TT<br>TTATGTTCTGCT |
| SEC24C | Anti-Rabbit-Nanobody-A8 | NB - TT<br>ATGTTAATGGGT |
| SEC16A | Anti-Rabbit-Nanobody-A38 | NB - TT<br>ATTTAGTGTAGC |
| LC3B | Anti-Rabbit-Nanobody-A20 | NB - TT<br>ATATGATCTCCG |
| SEC31A | Anti-Mouse-Nanobody-A27 | NB - TT<br>AAAAAGTTCGAG |
| ARFGAP1 | Anti-Rabbit-Nanobody-A10 | NB – TT<br>ATAATGGATGGG |
| COPI | Anti-Mouse-Nanobody-A25 | NB – TT<br>ATAATCATGCTC |
| p62 | Anti-Mouse-Nanobody-A36 | NB – TT<br>ATAACAGAATCG |

**Table S2.** Imager Sequence.

| **Imager name** | **Sequence** | **3'-mod** |
| --- | --- | --- |
| R2-6nt | TGGTGG | Cy3B |

**Table S3.** Adaptor Sequences.

| **Adaptor Name** | **Sequence (3' → 5')** |
| --- | --- |
| A3-5xR2 | ACCACCACCACCACCACCA AA<br>CGCTAATGAA |
| A15-5xR2 | ACCACCACCACCACCACCA AA<br>TCCAATCACT |
| A39-5xR2 | ACCACCACCACCACCACCA AA<br>AGCAGAACAT |
| A8-5xR2 | ACCACCACCACCACCACCA AA<br>ACCCATTAAC |
| A38-5xR2 | ACCACCACCACCACCACCA AA<br>GCTACACTAA |
| A20-5xR2 | ACCACCACCACCACCACCA AA<br>CGGAGATCAT |
| A27-5xR2 | ACCACCACCACCACCACCA AA<br>CTCGAACTTT |
| A10-5xR2 | ACCACCACCACCACCACCA AA<br>CCCATCCATT |
| A25-5xR2 | ACCACCACCACCACCACCA AA<br>GAGCATGATT |
| A36-5xR2 | ACCACCACCACCACCACCA AA<br>CGATTCTGTT |

**Table S4.** Eraser sequences.

| **Eraser Name** | **Sequence (3' → 5')** |
| --- | --- |
| E3-5xR2 | TTCATTAGCG TT TGG |
| E15-5xR2 | AGTGATTGGA TT TGG |
| E39-5xR2 | ATGTTCTGCT TT TGG |
| E8-5xR2 | GTTAATGGGT TT TGG |
| E38-5xR2 | TTAGTGTAGC TT TGG |
| E20-5xR2 | ATGATCTCCG TT TGG |
| E27-5xR2 | AAAGTTCGAG TT TGG |
| E10-5xR2 | AATGGATGGG TT TGG |
| E25-5xR2 | AATCATGCTC TT TGG |
| E36-5xR2 | AACAGAATCG TT TGG |

**Table S5.**
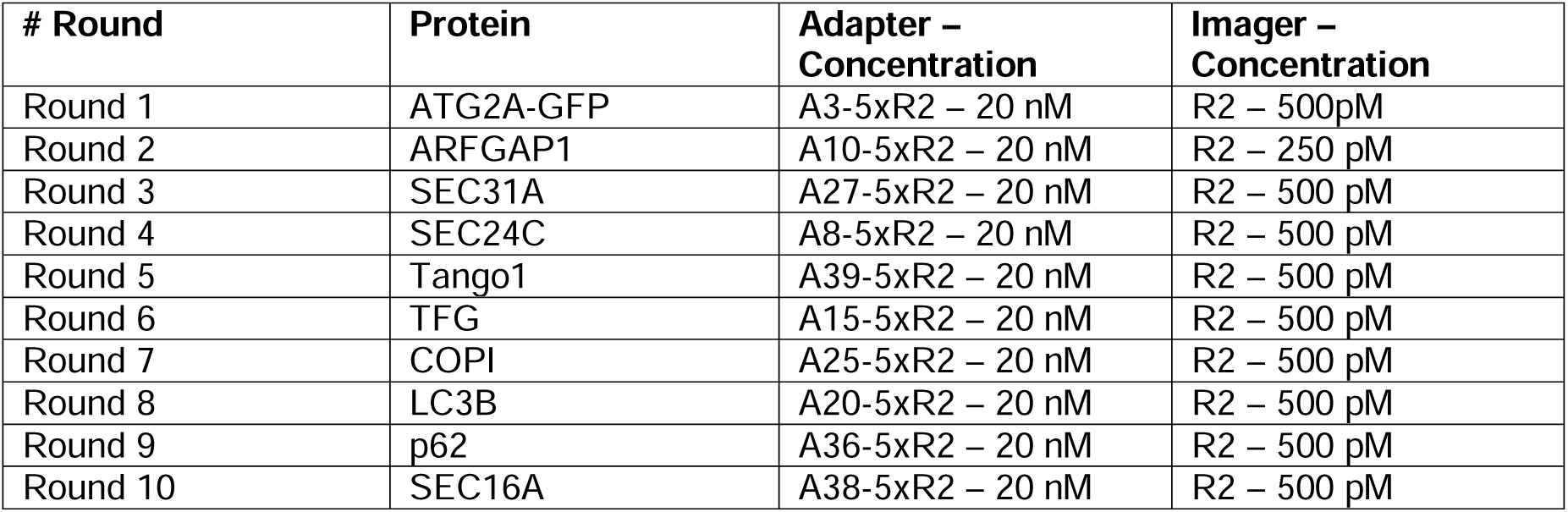
Order of sequential antibody labeling for FLASH-PAINT.

**Table S6.**
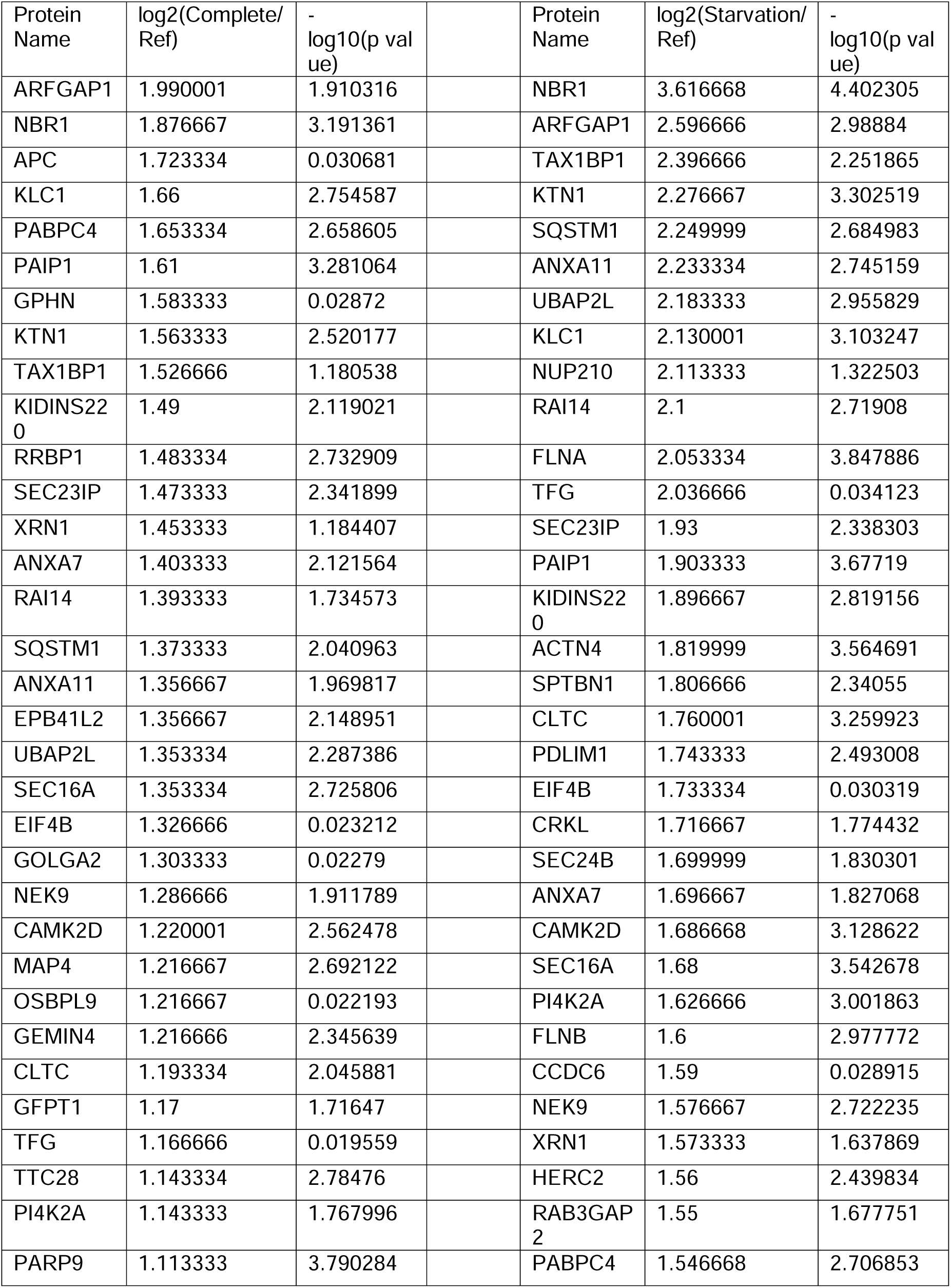
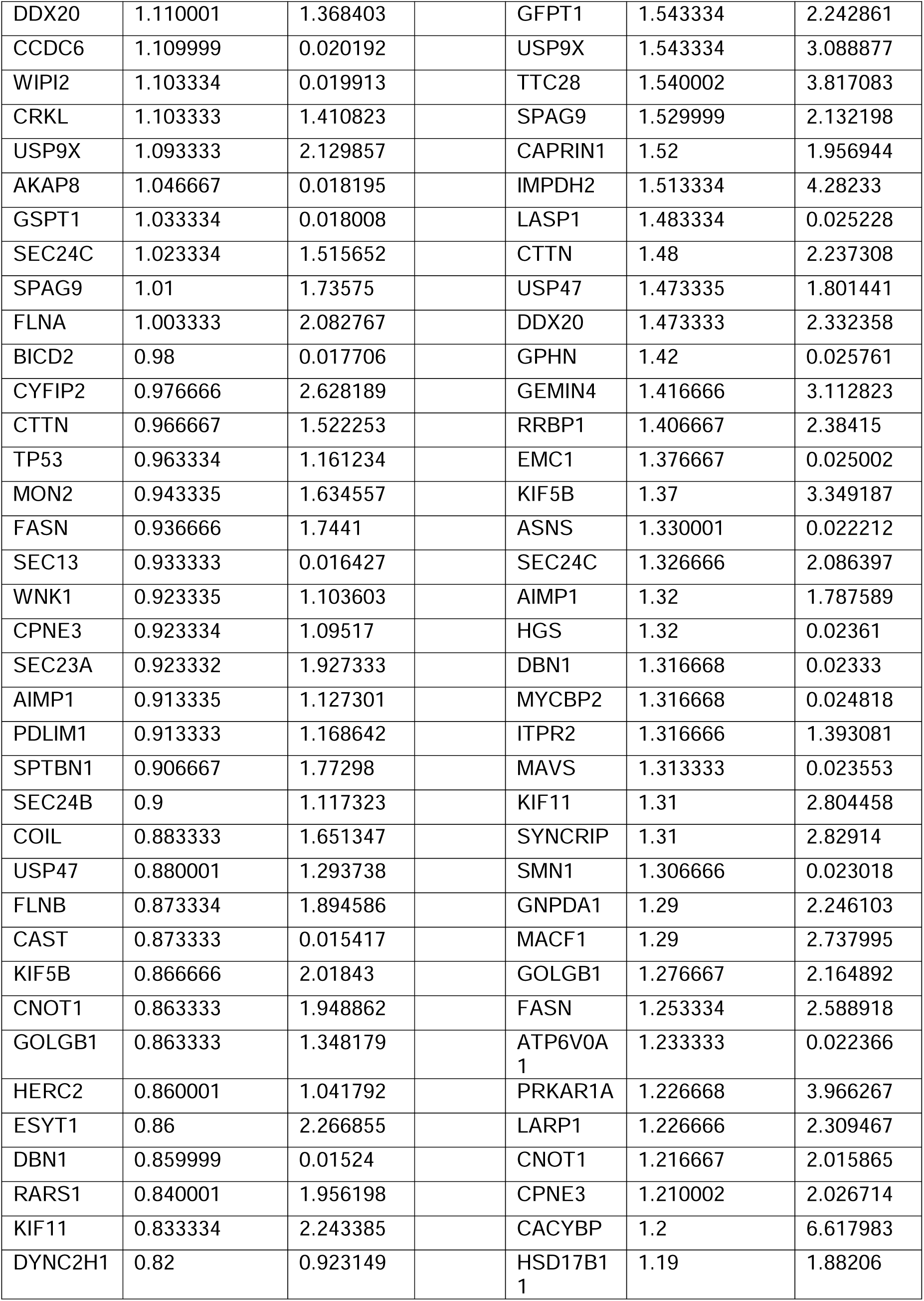
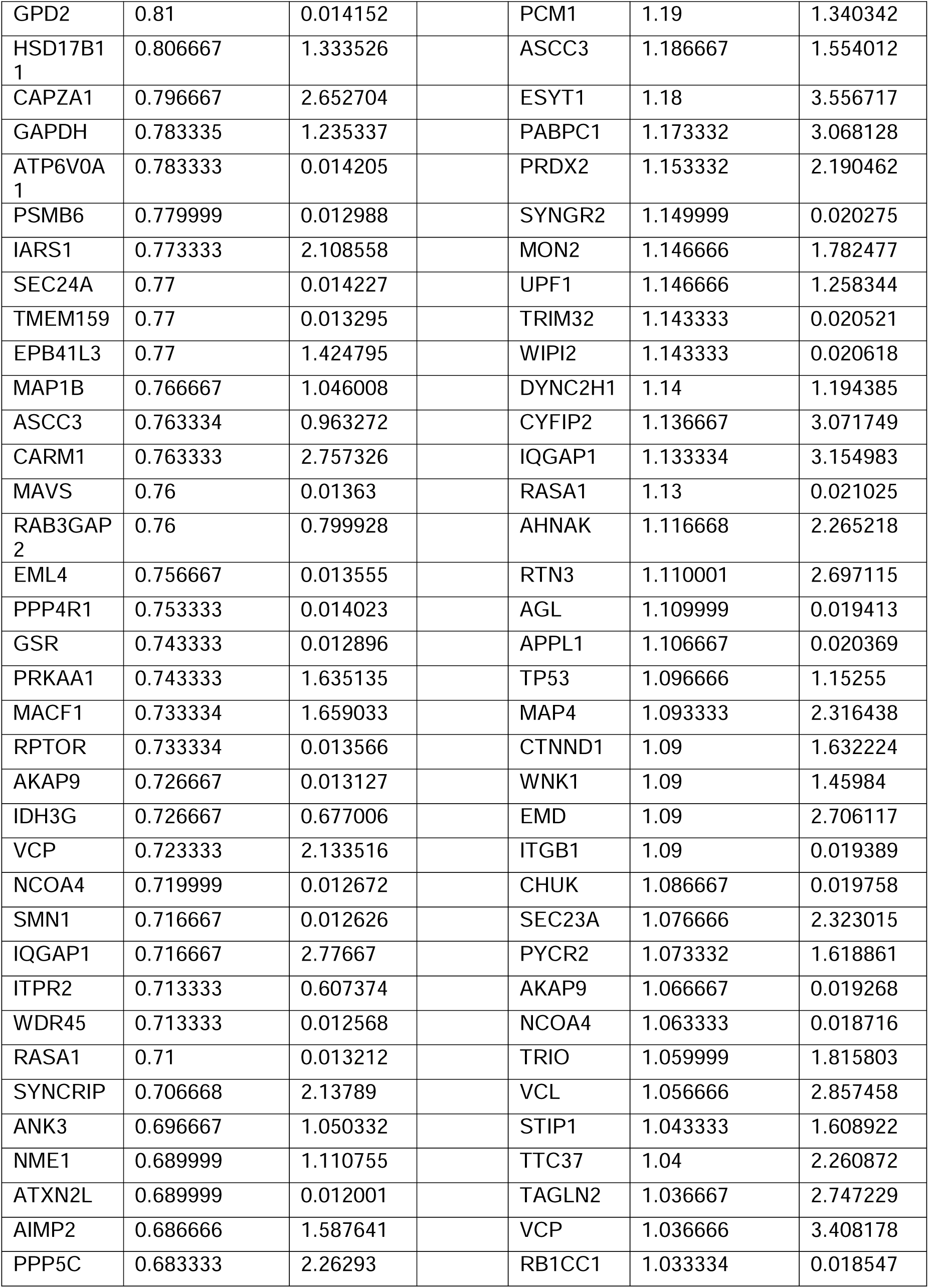
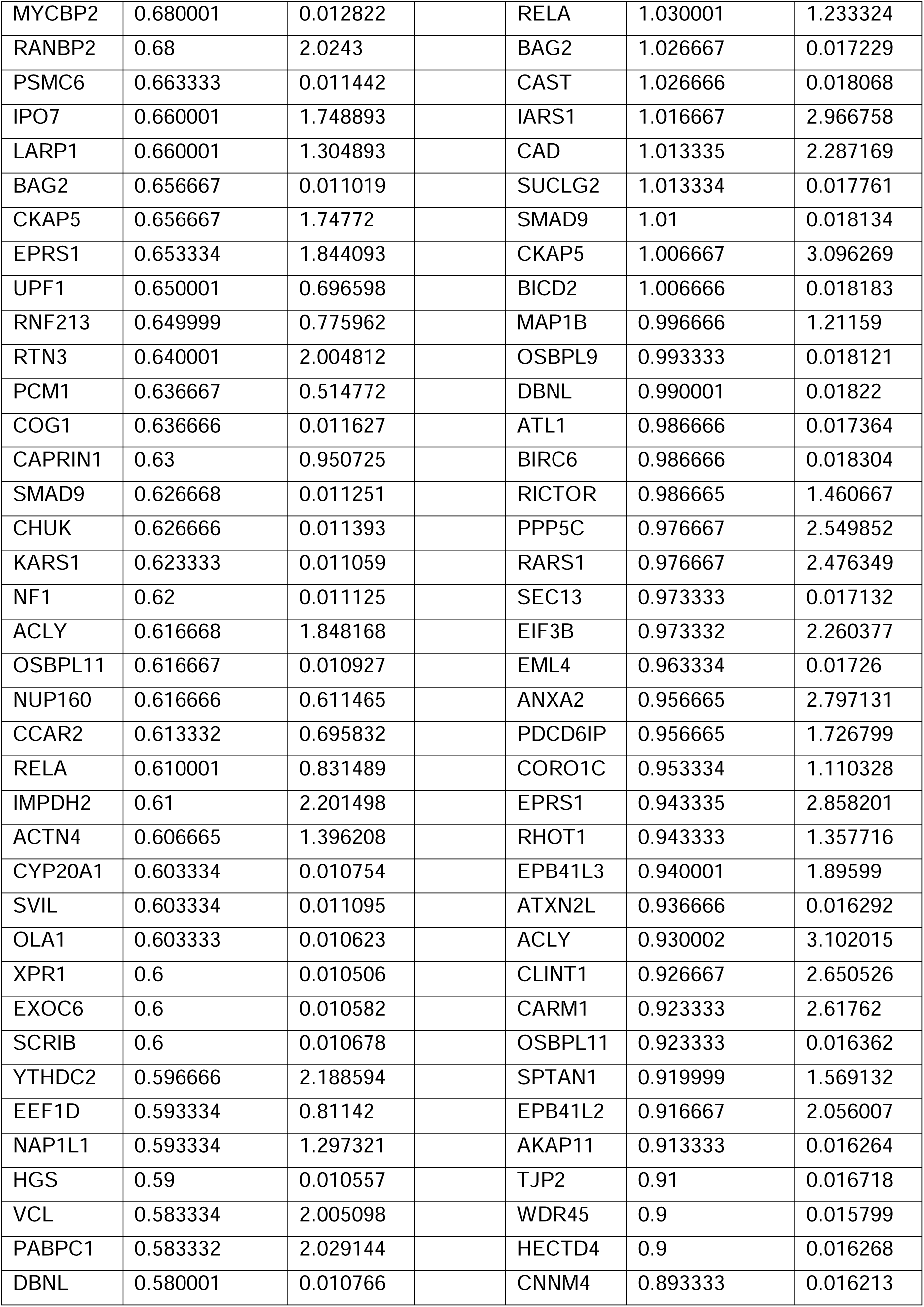
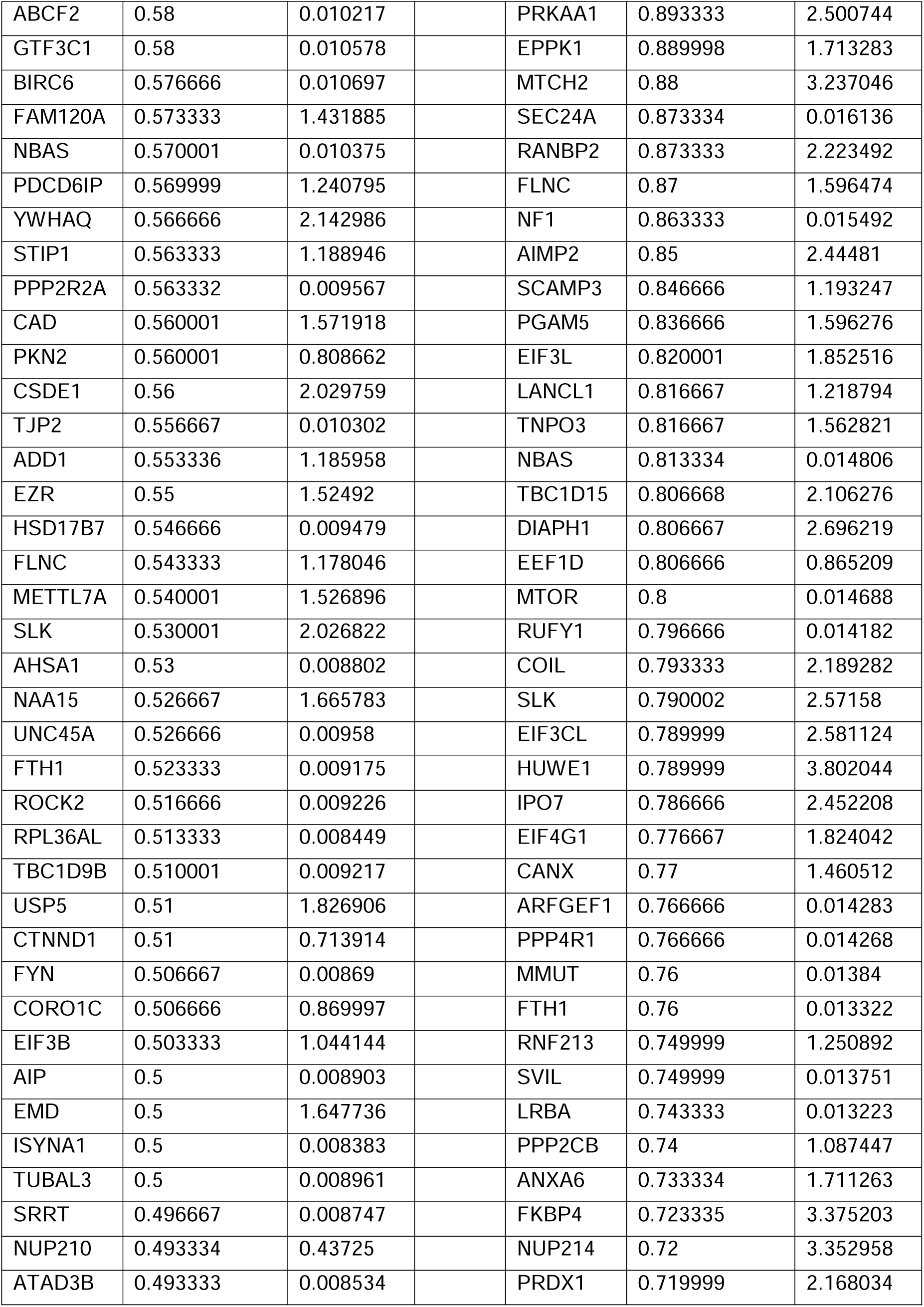
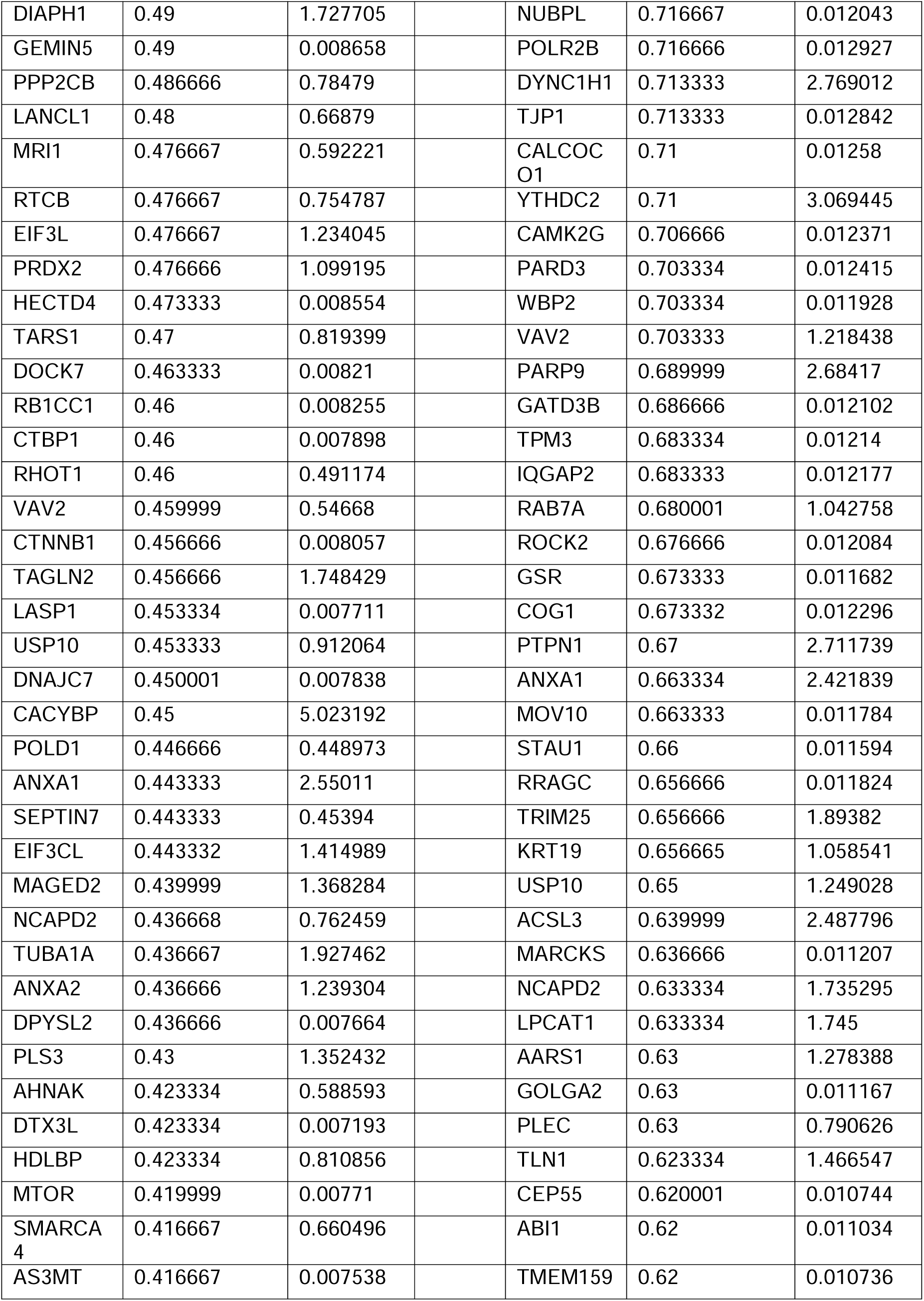
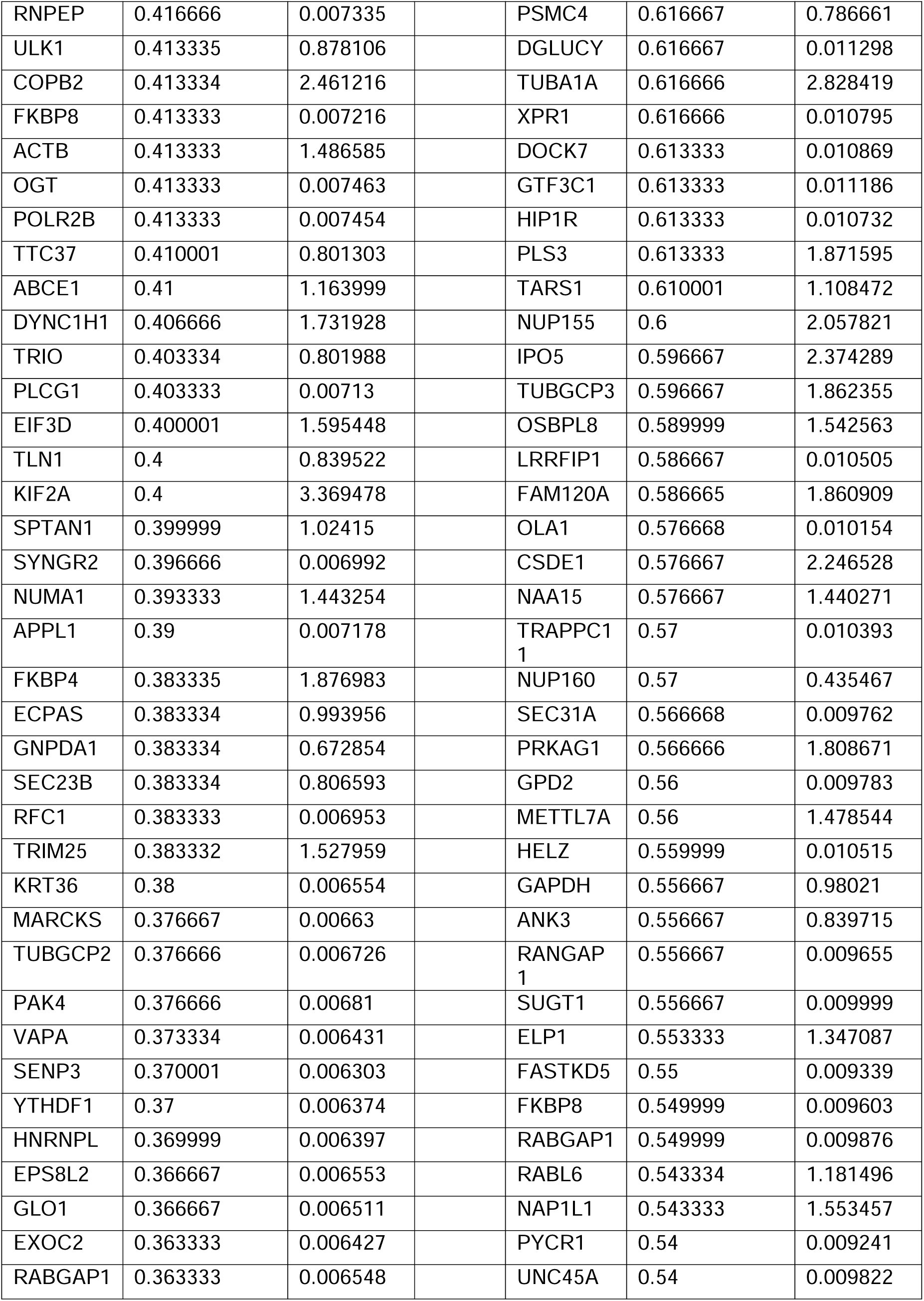
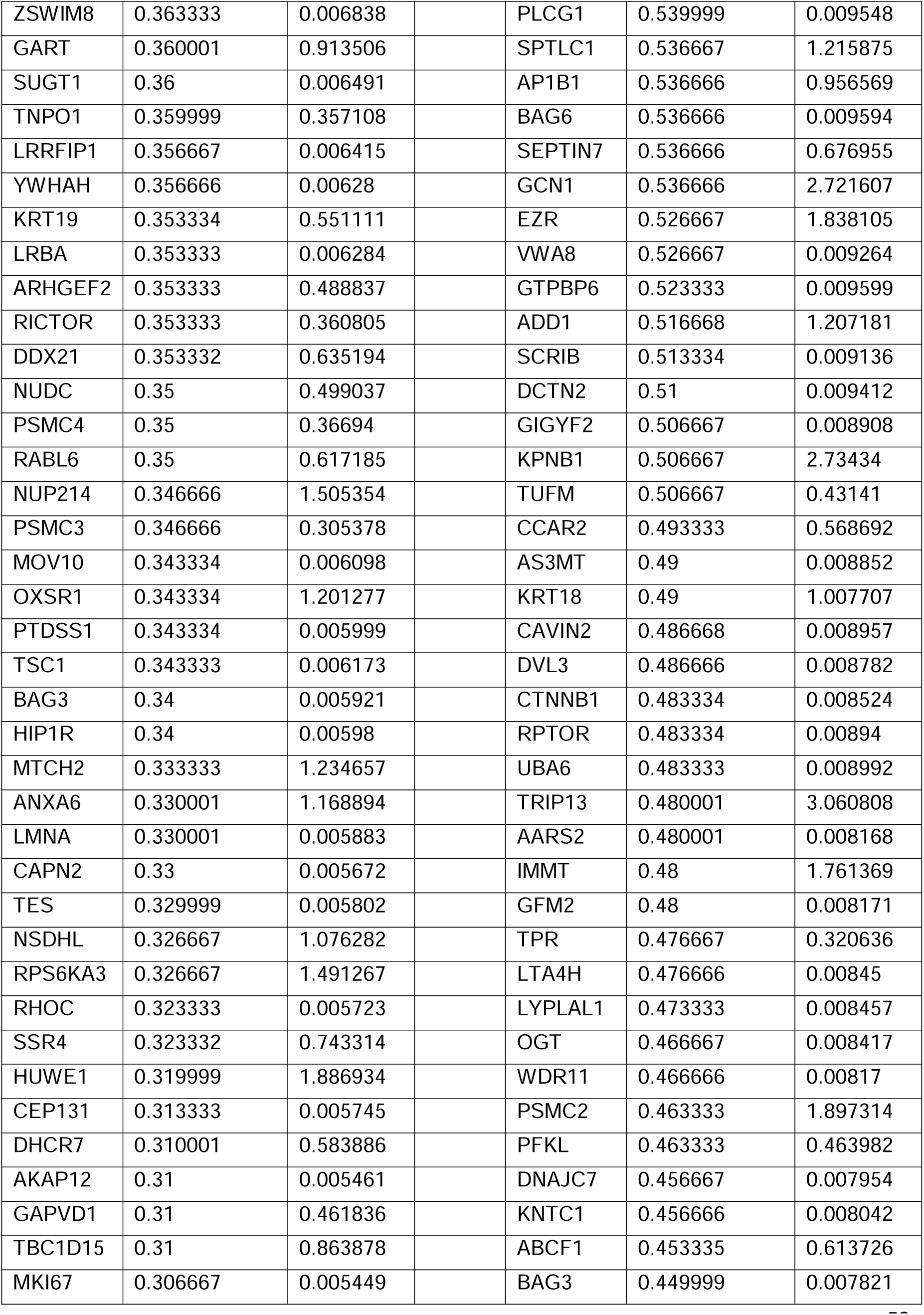
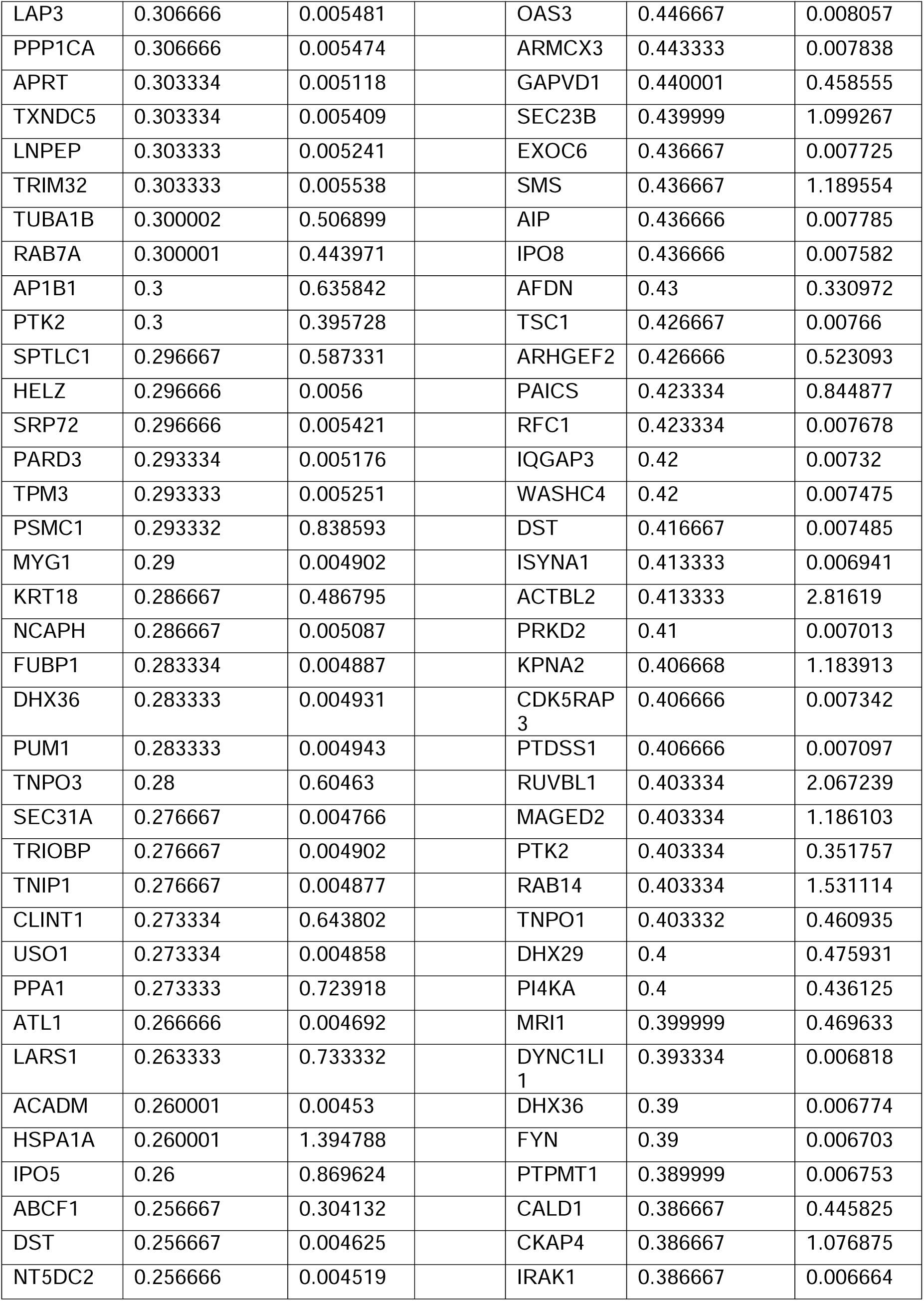
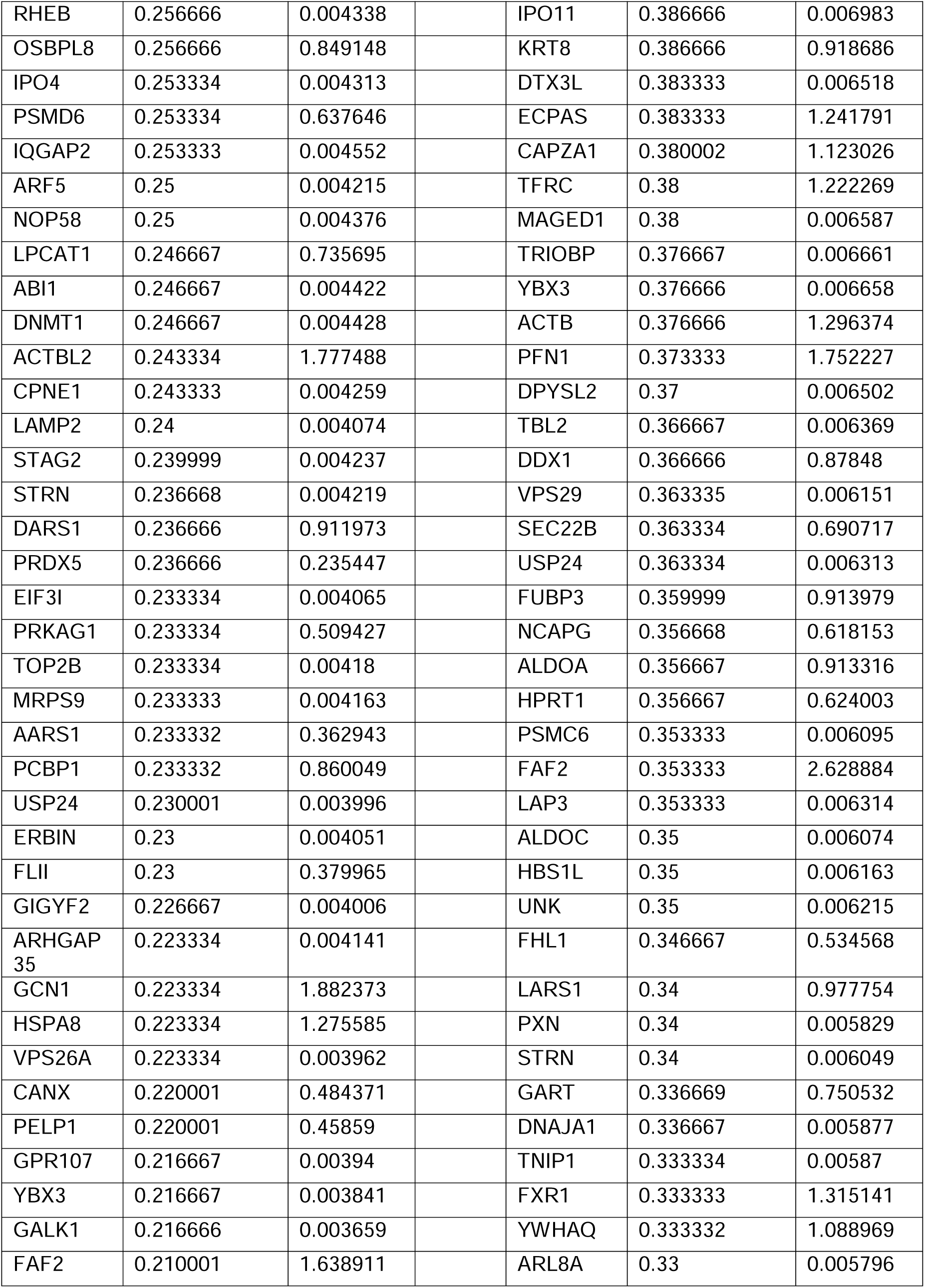
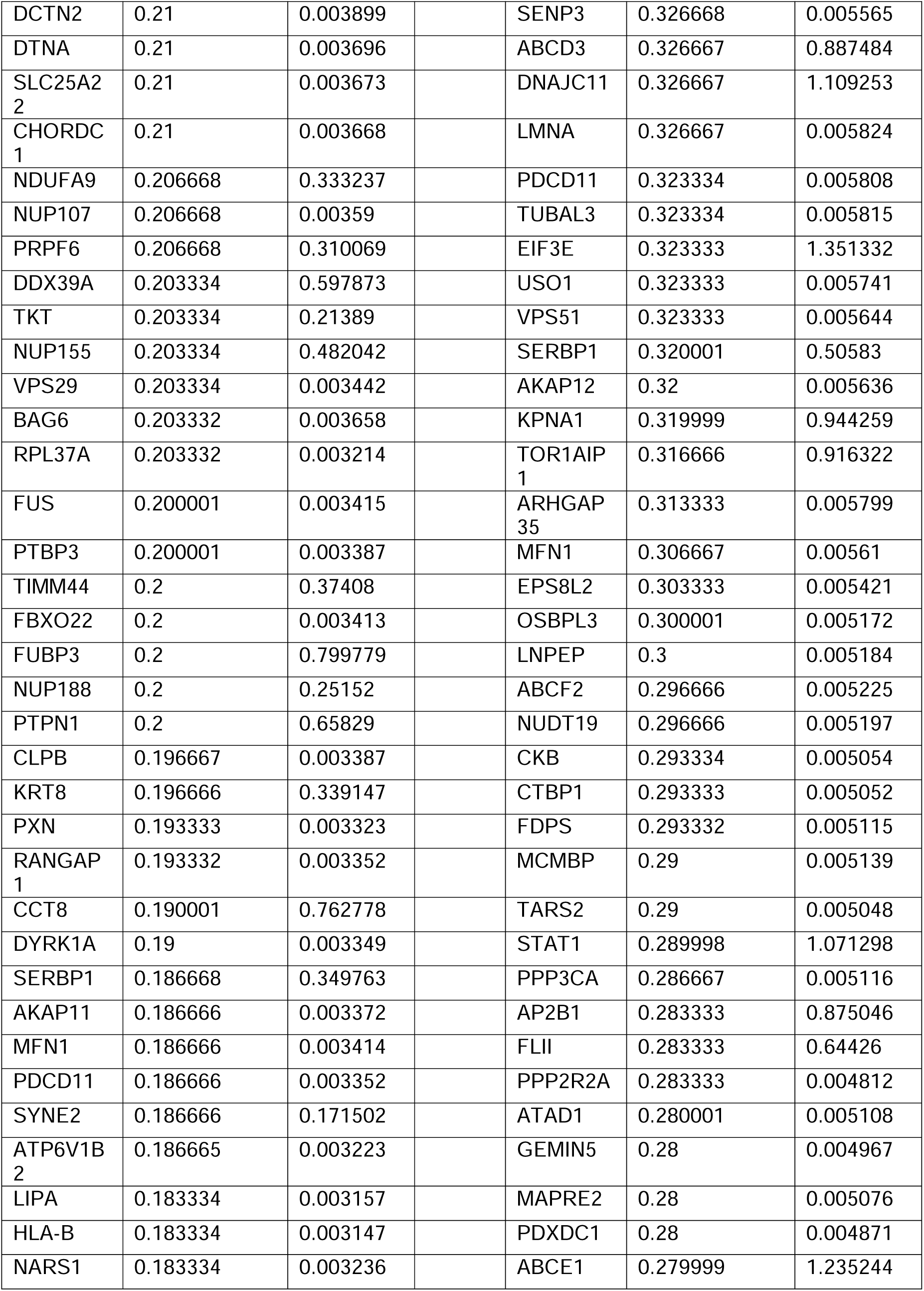
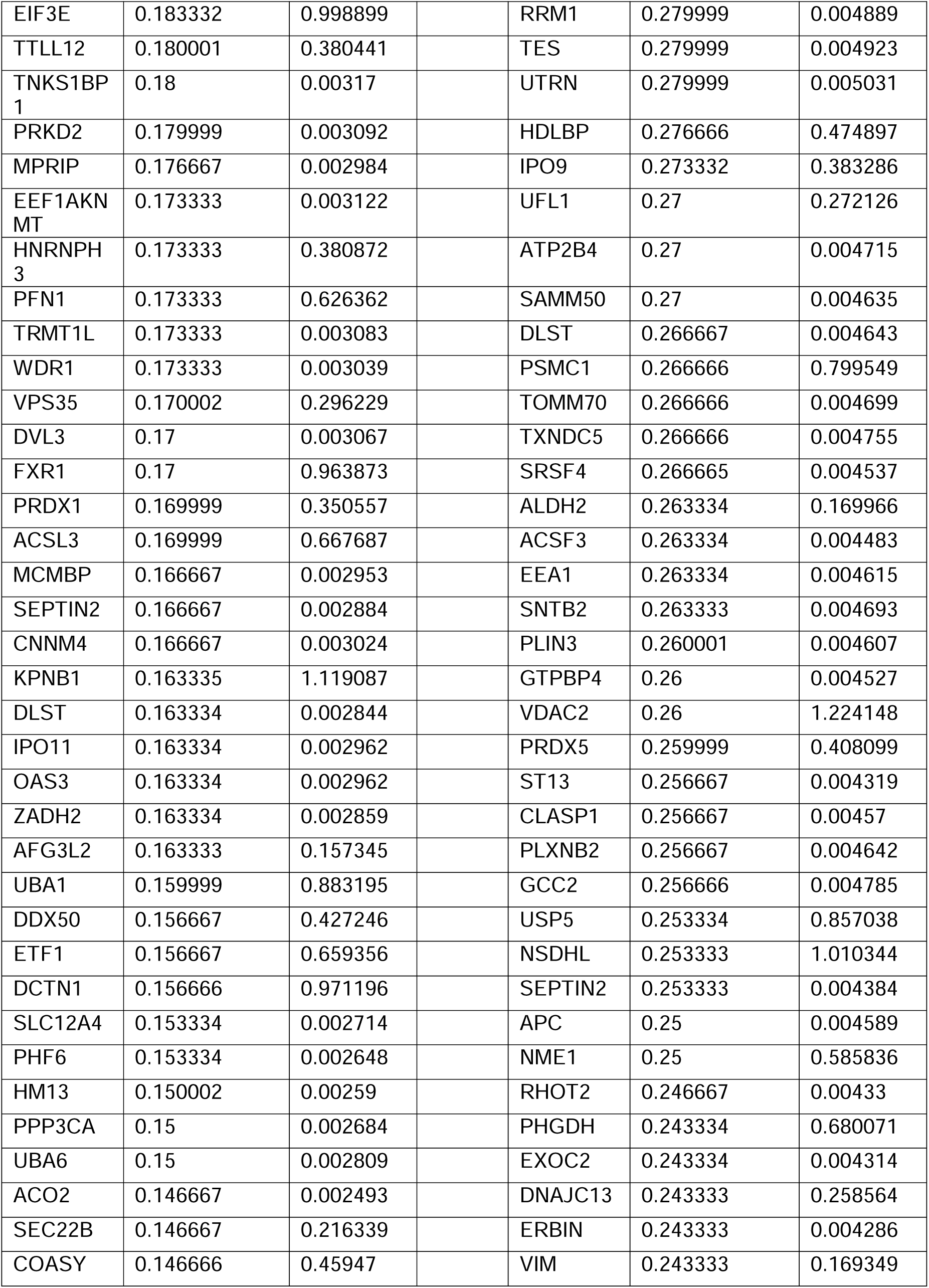
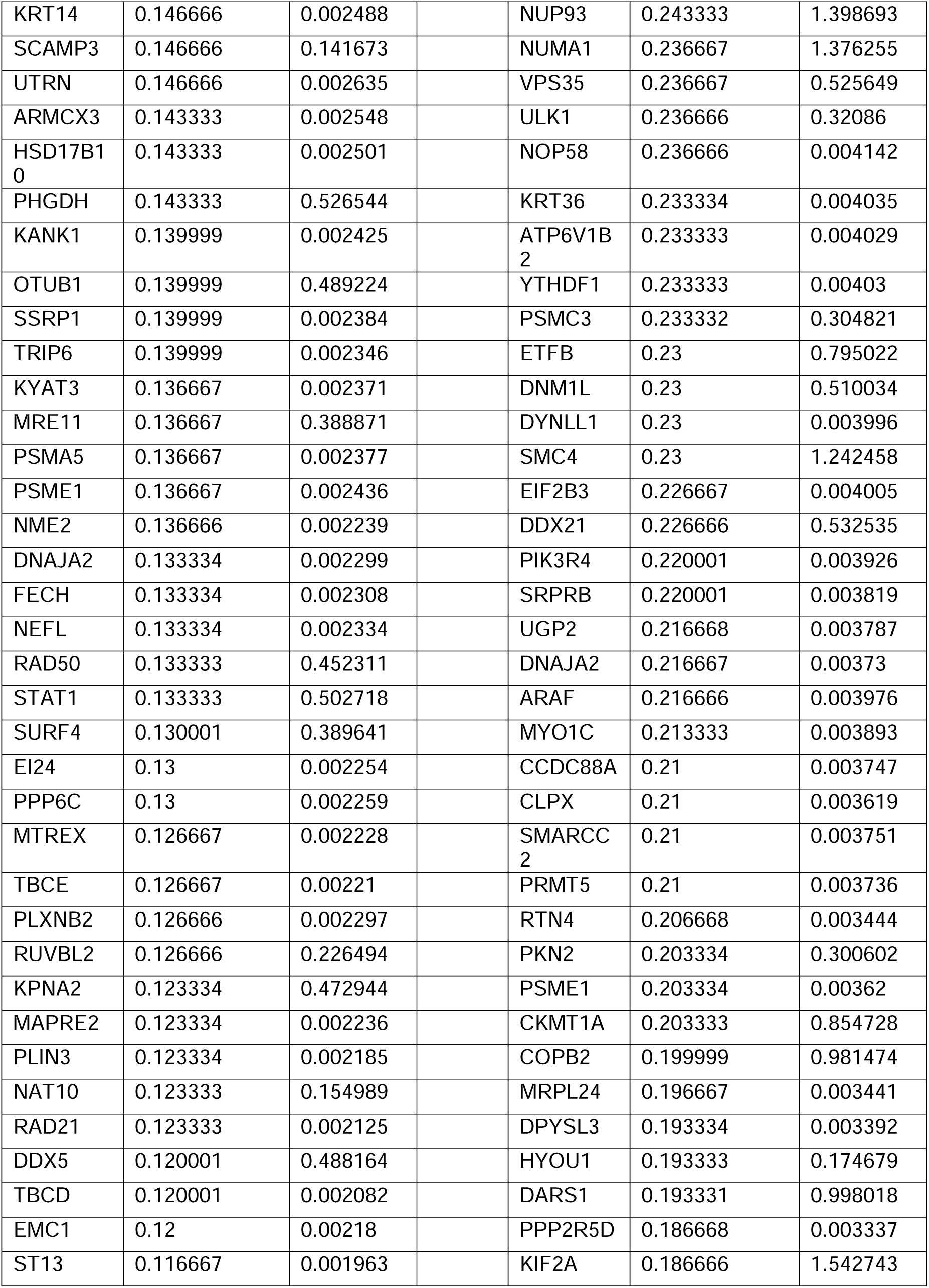
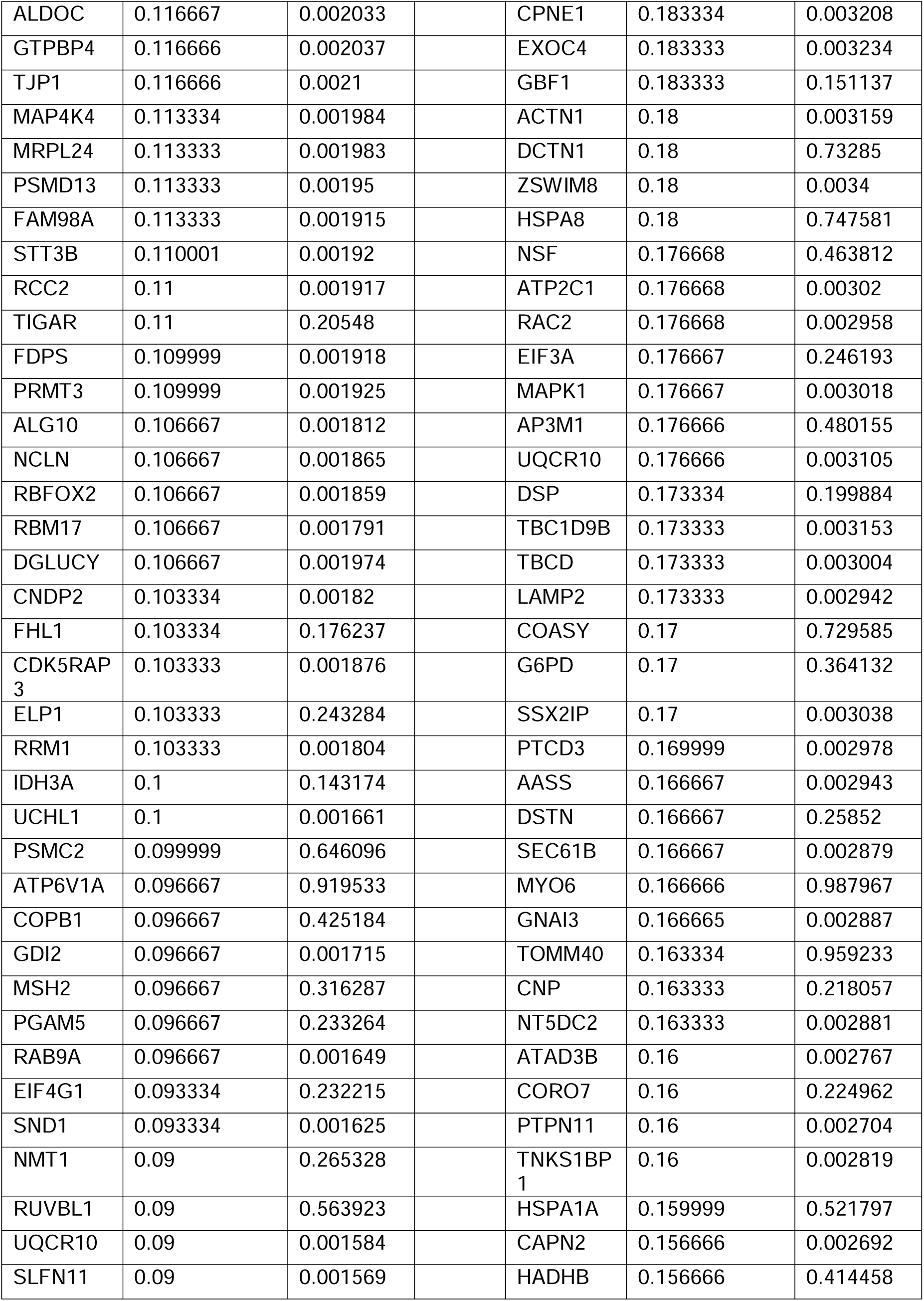
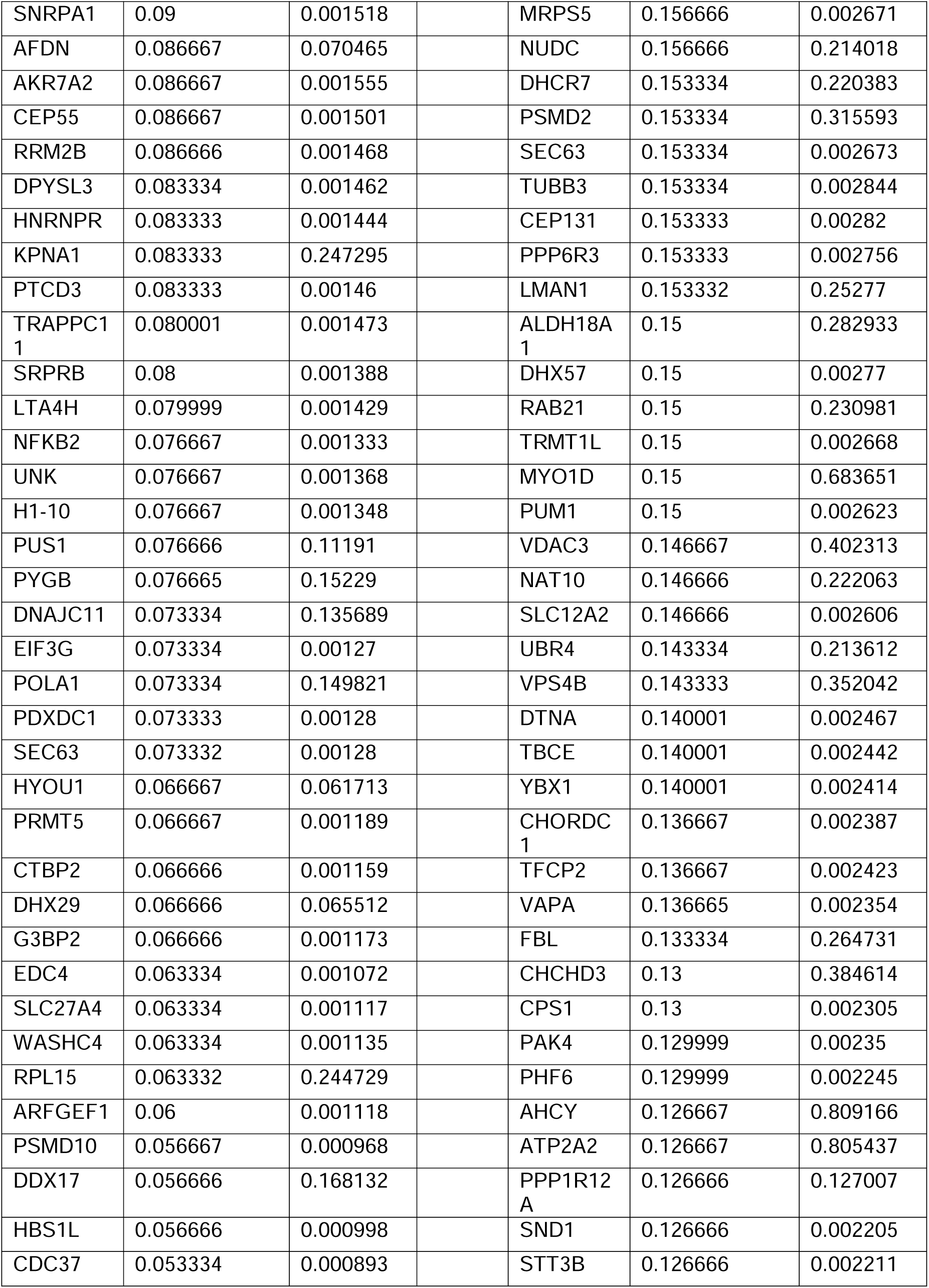
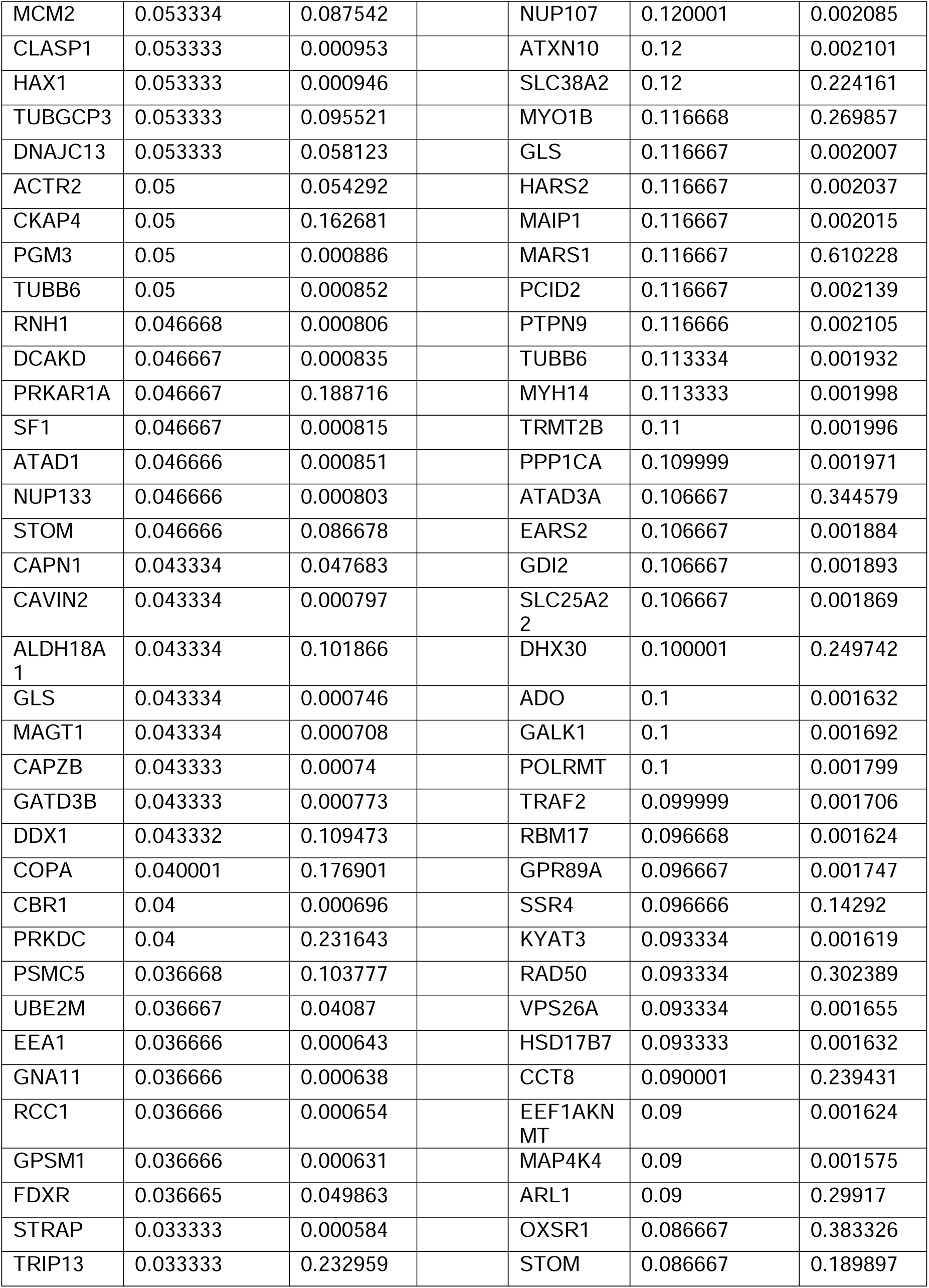
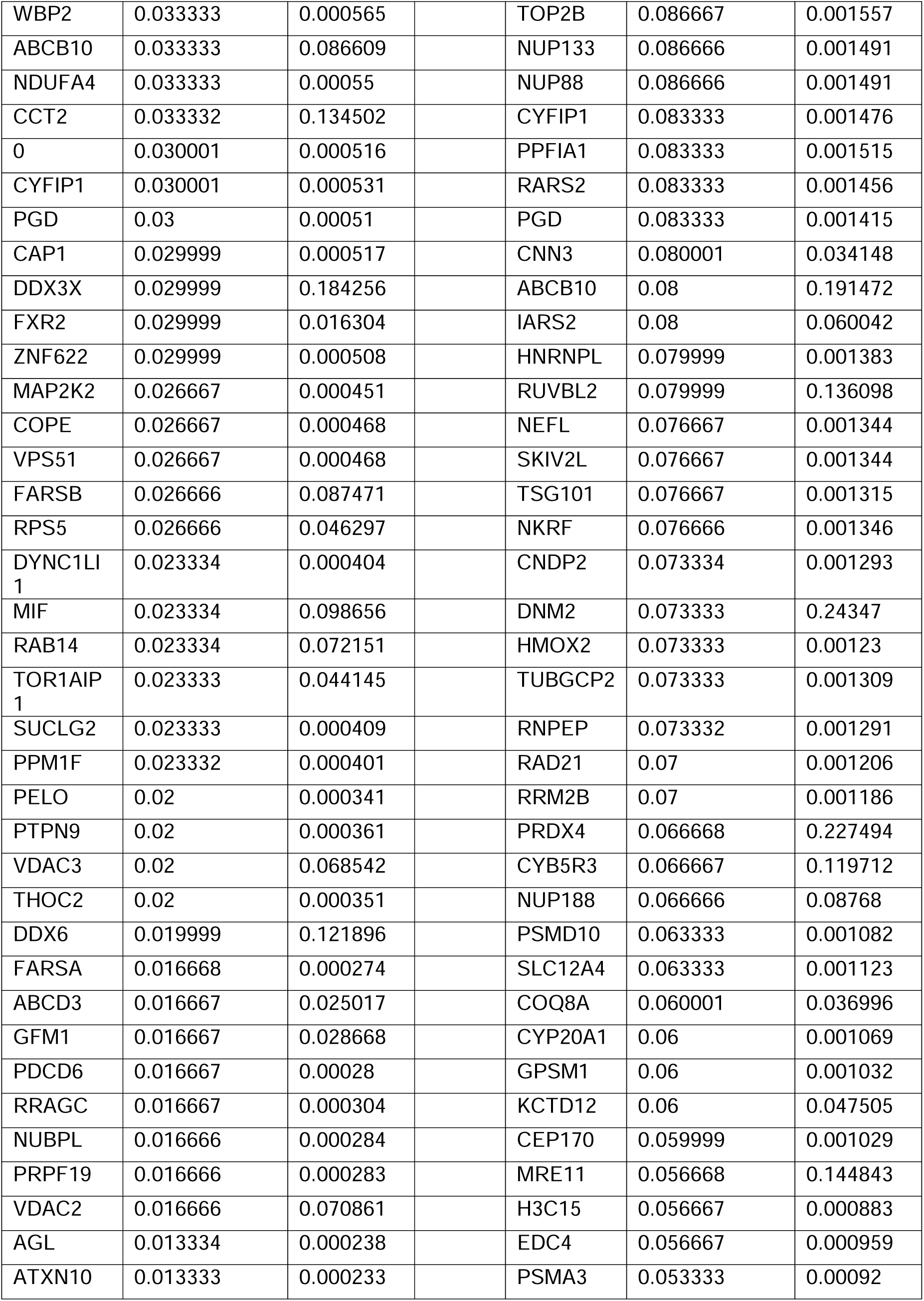
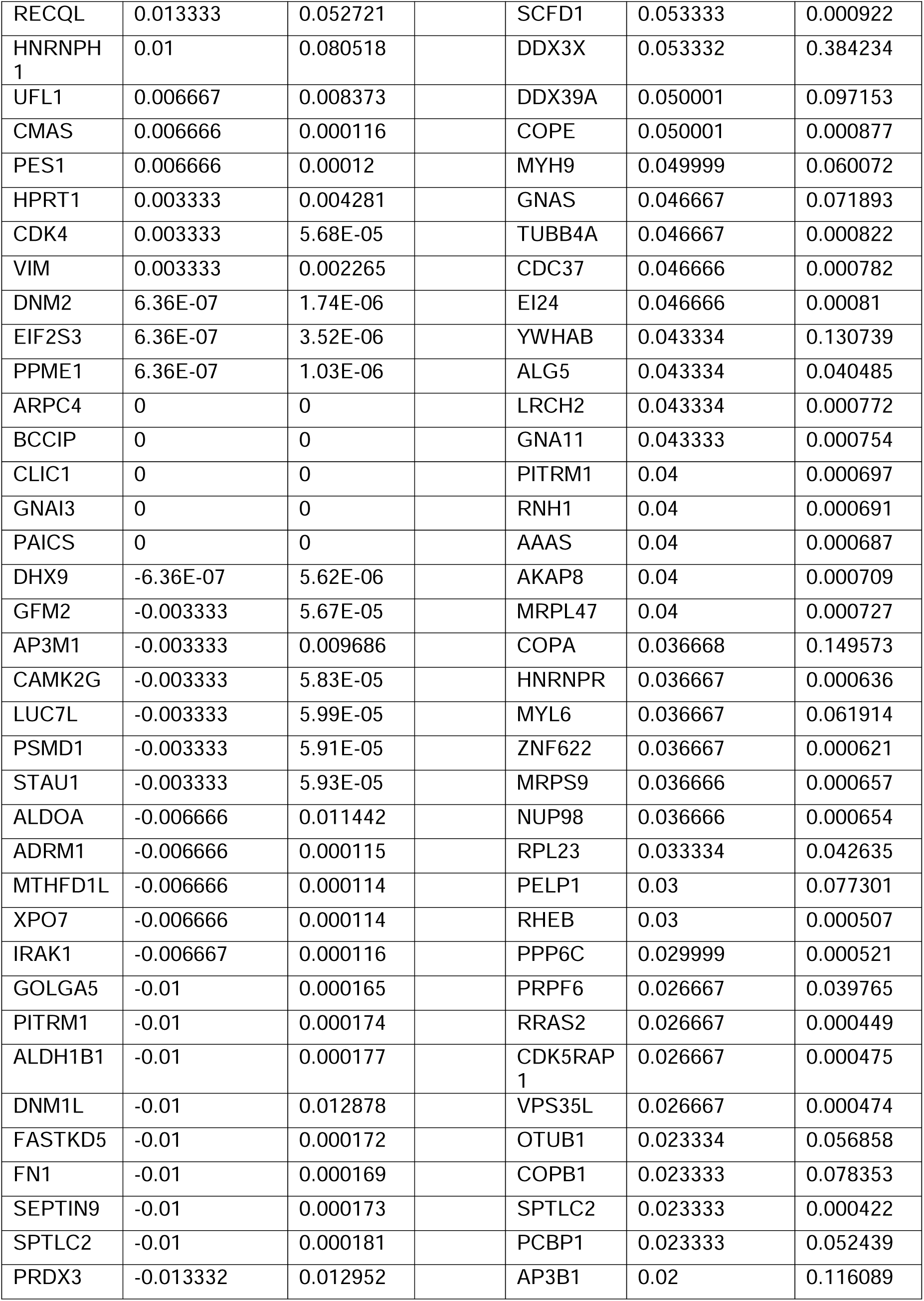
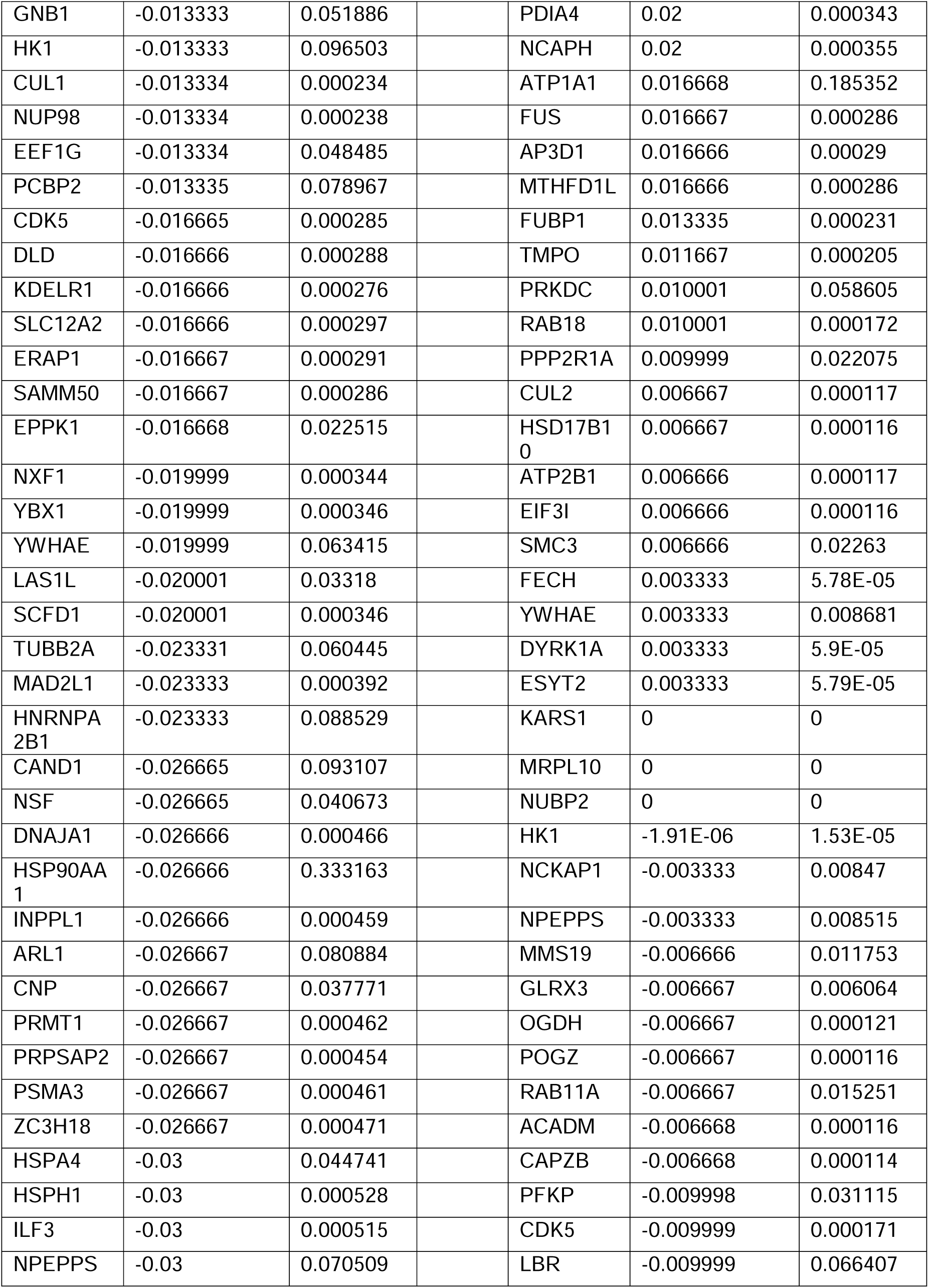
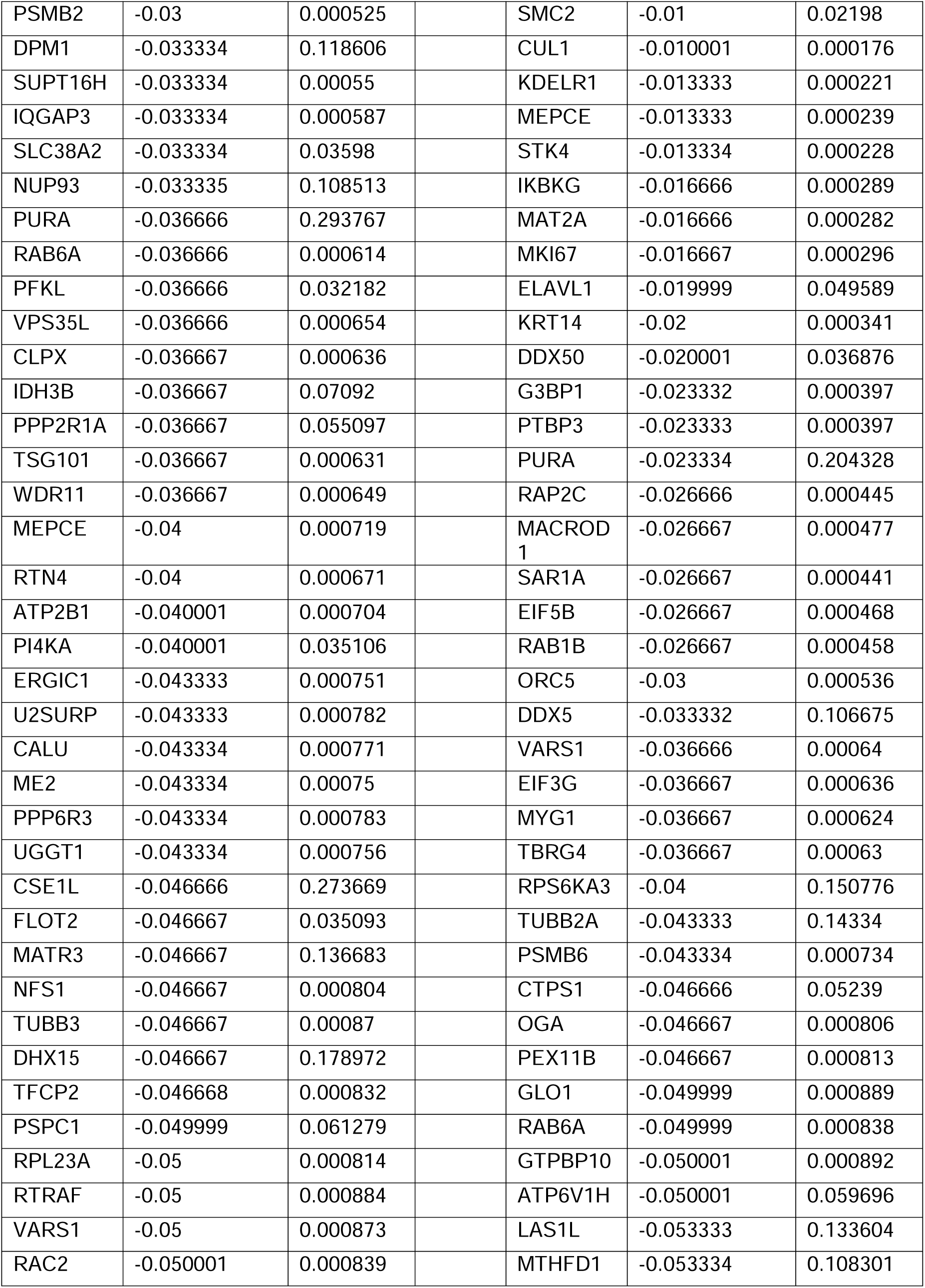
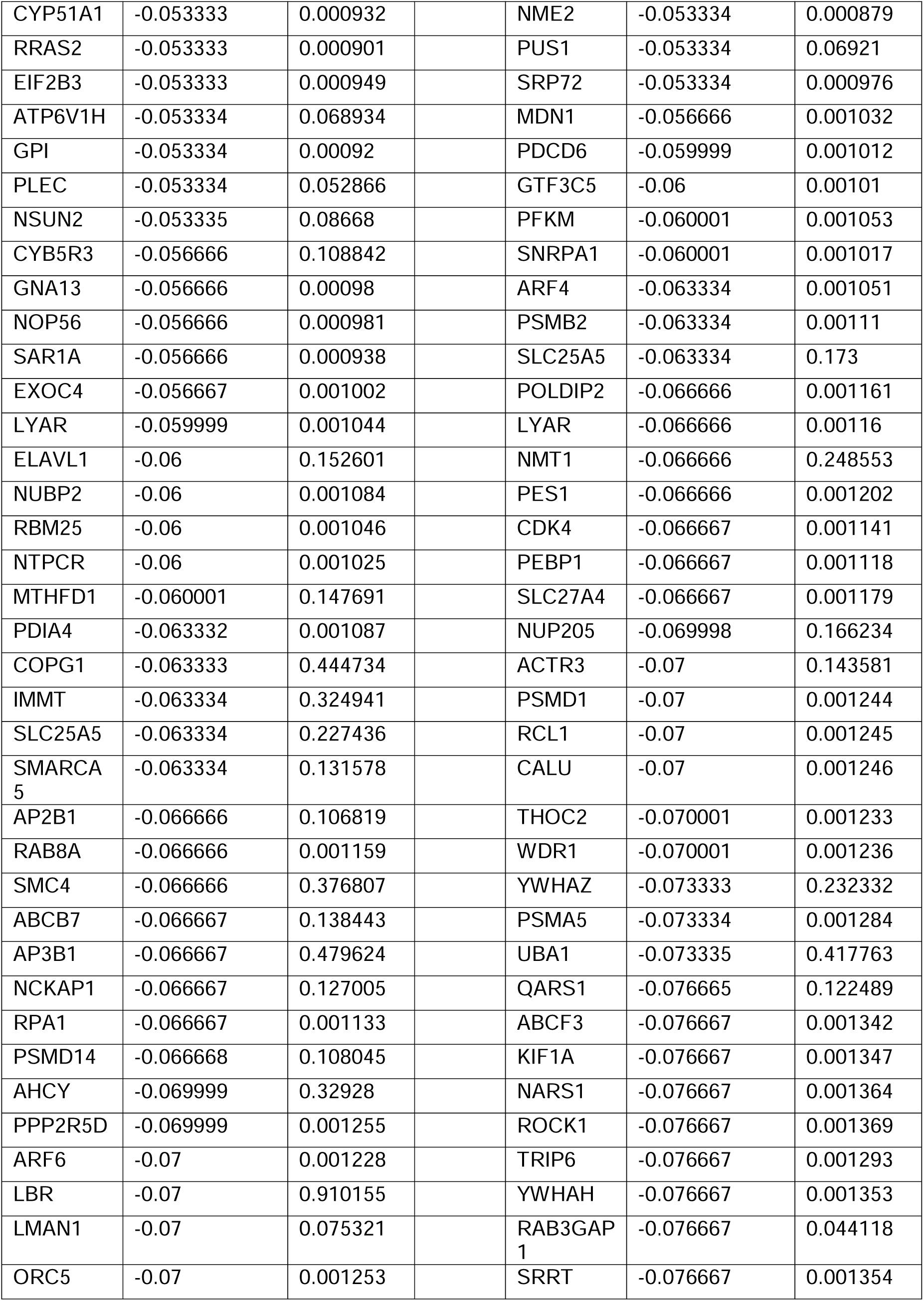
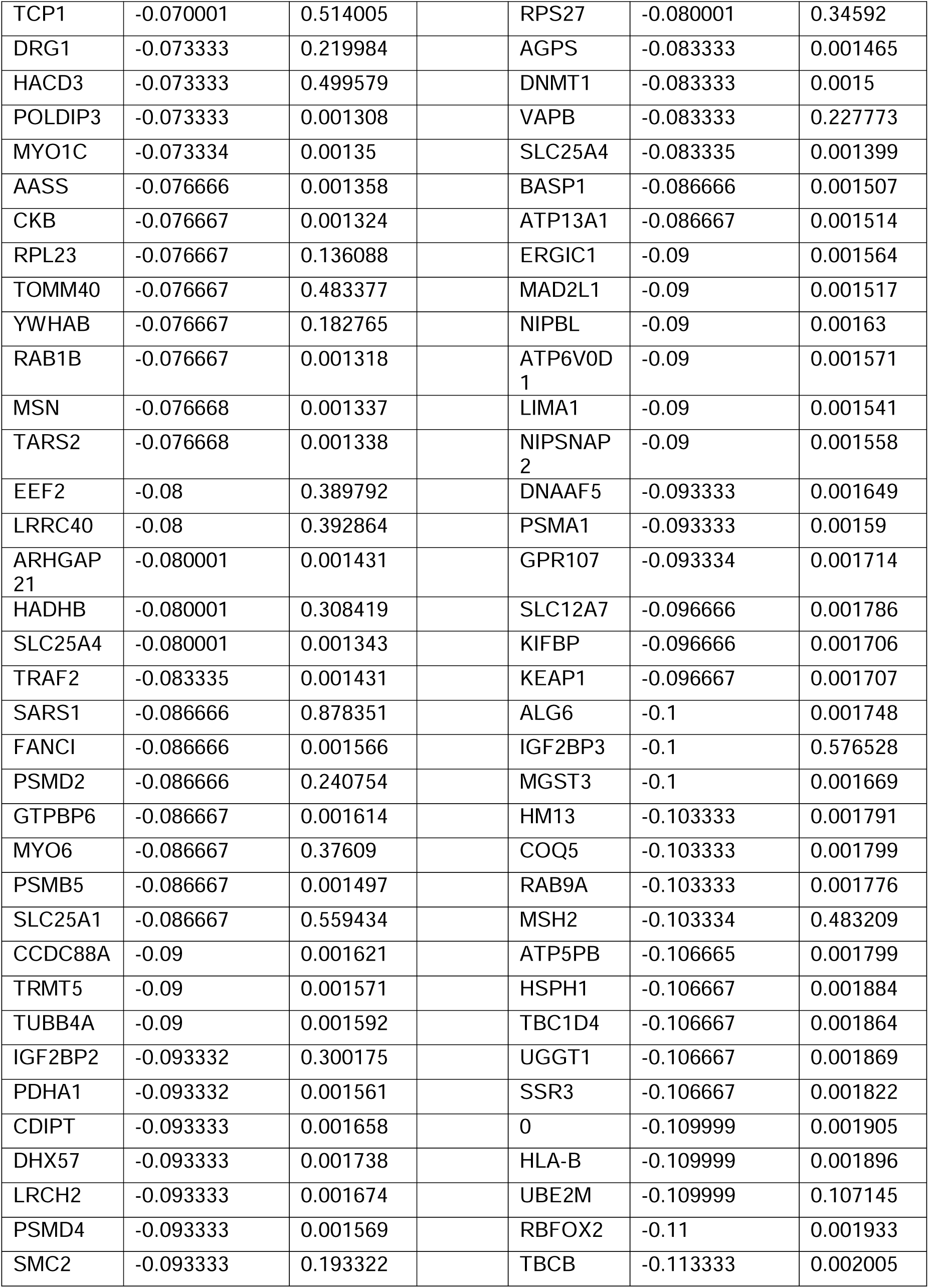
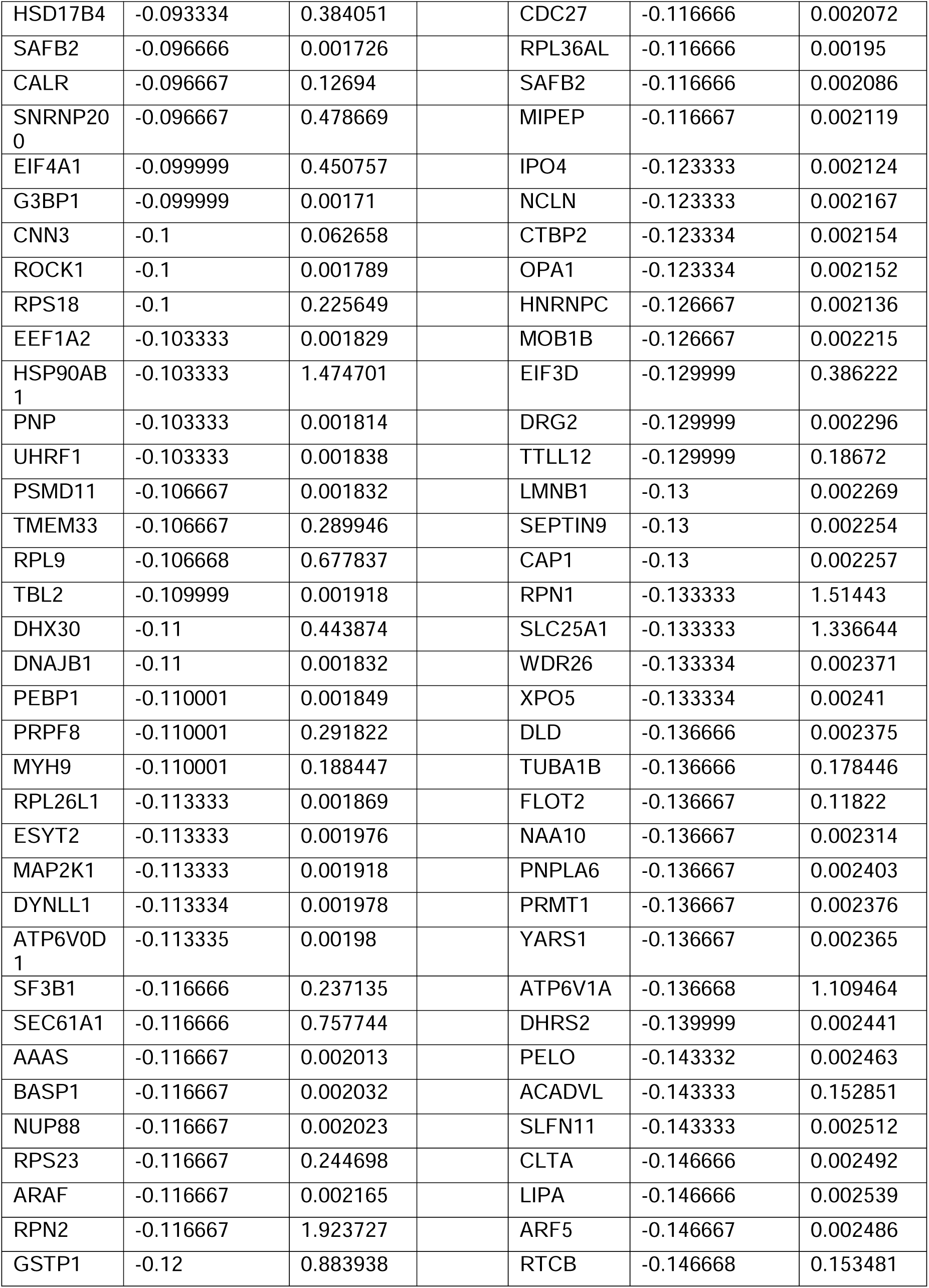
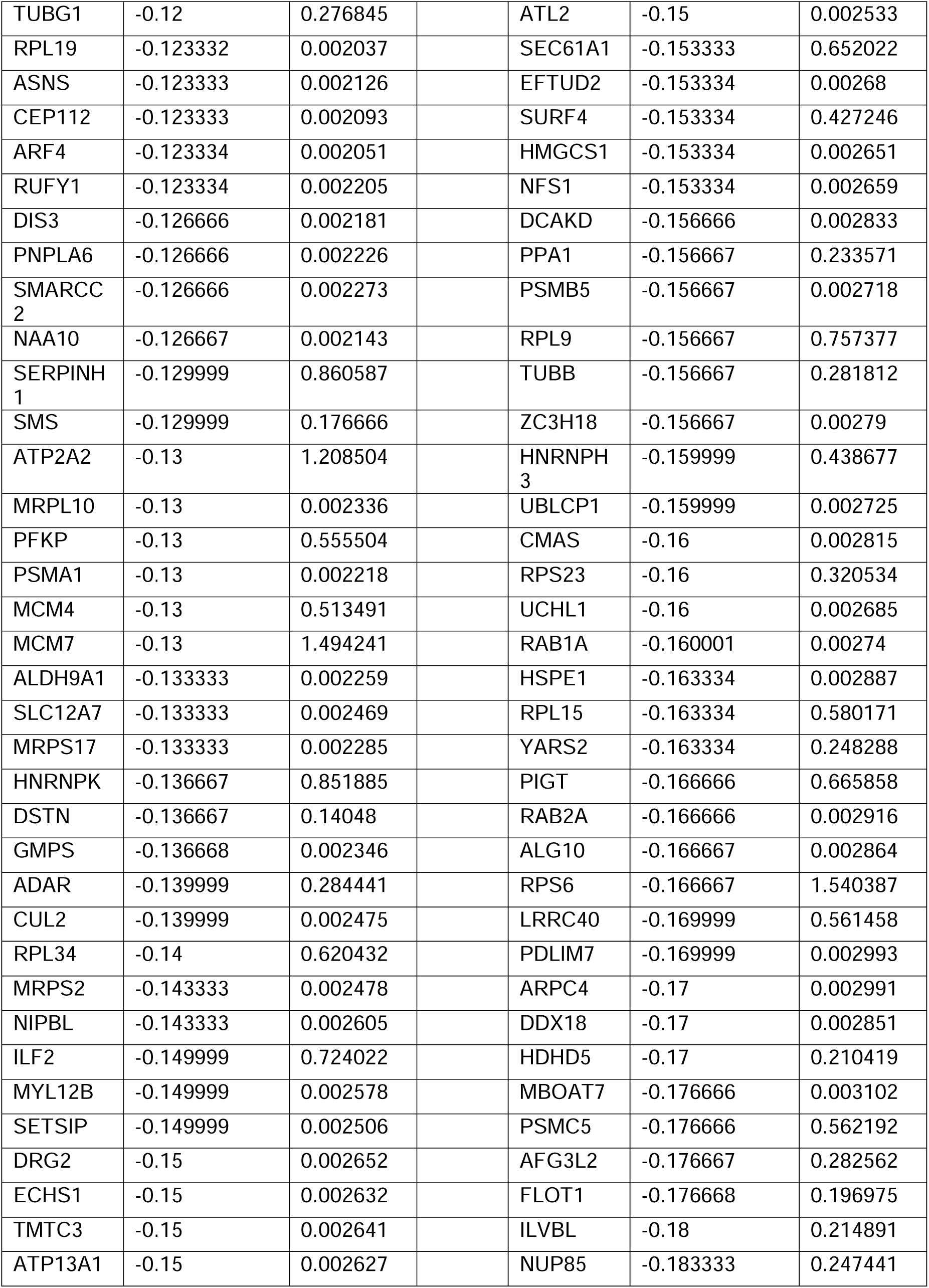
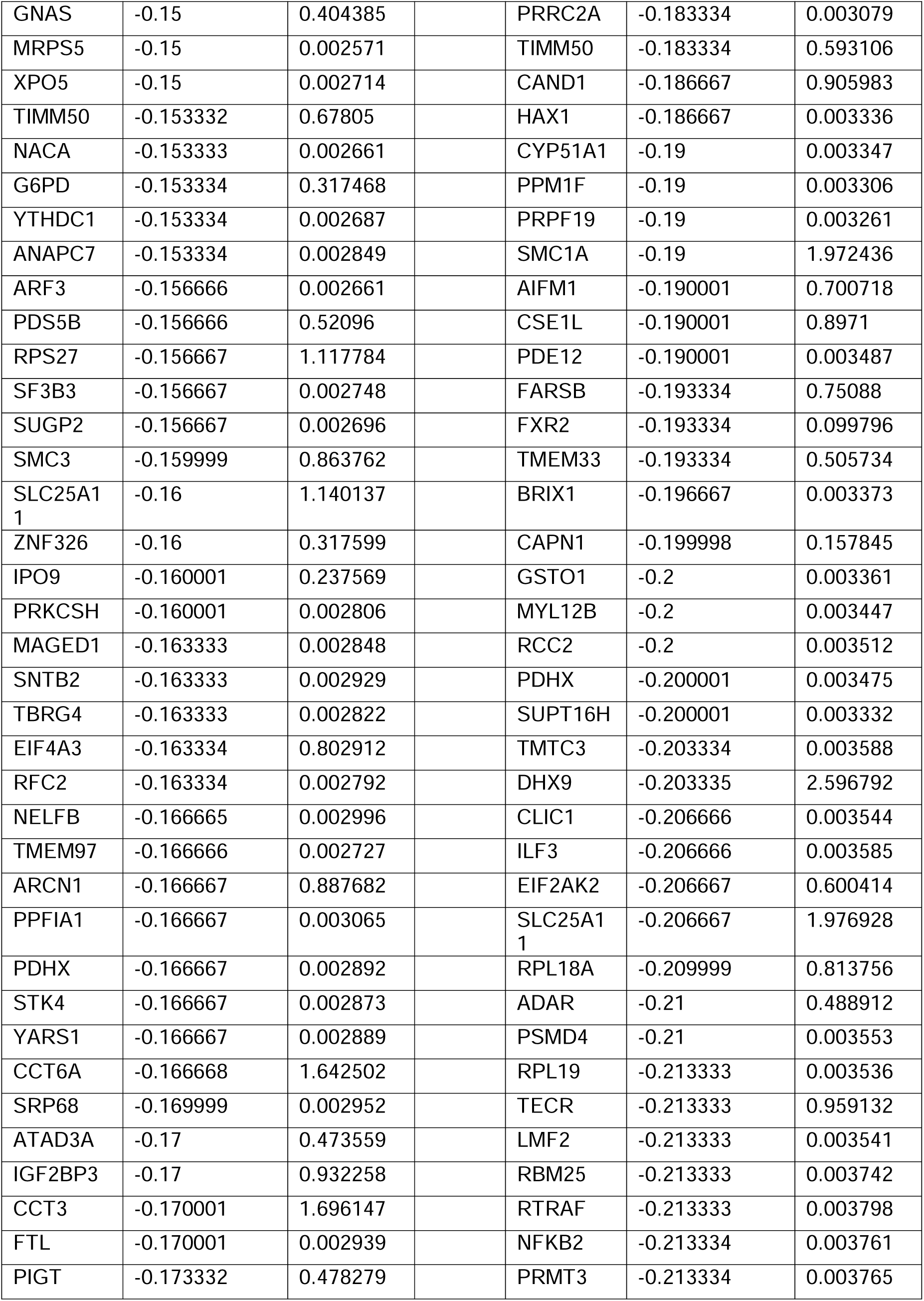
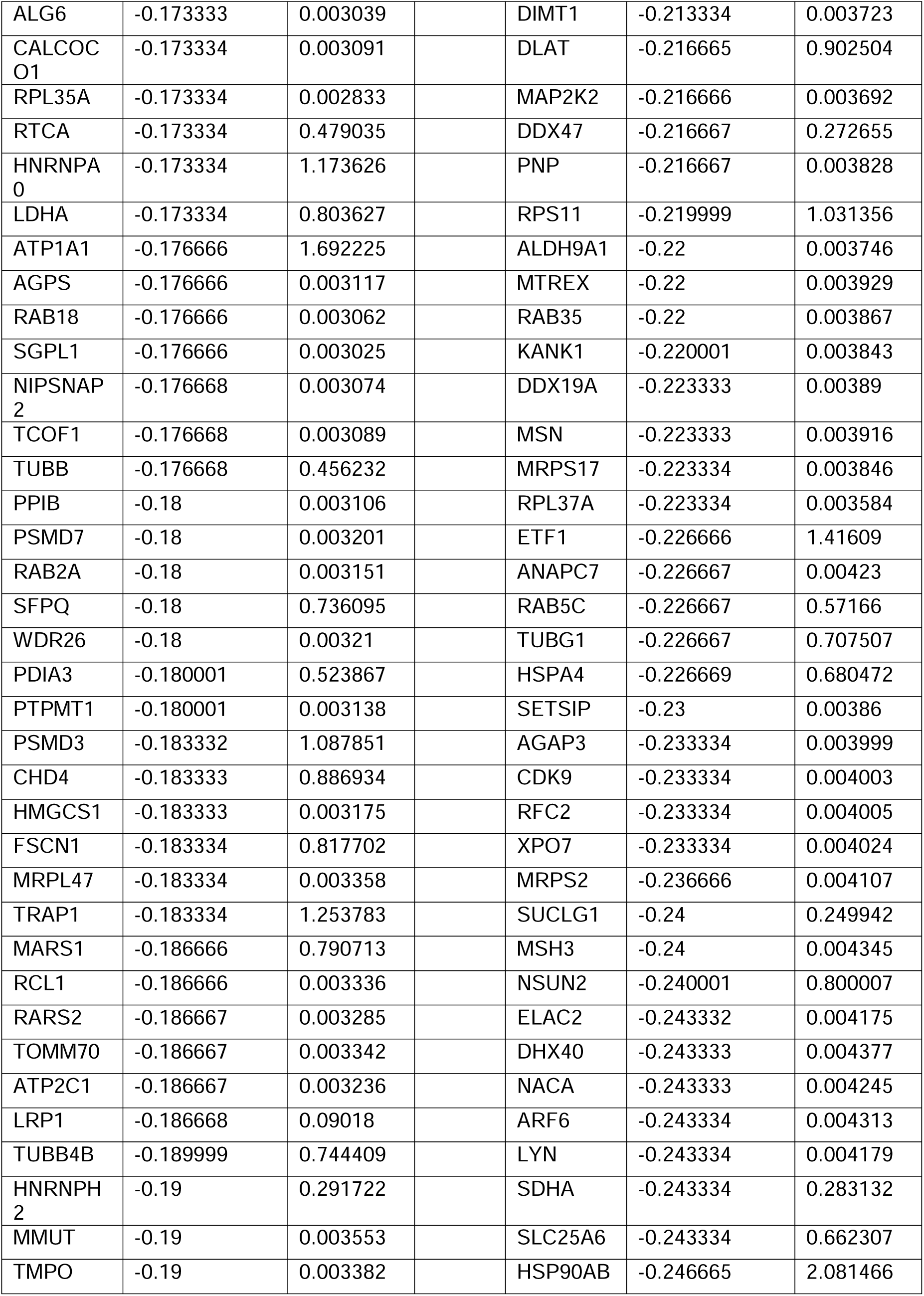
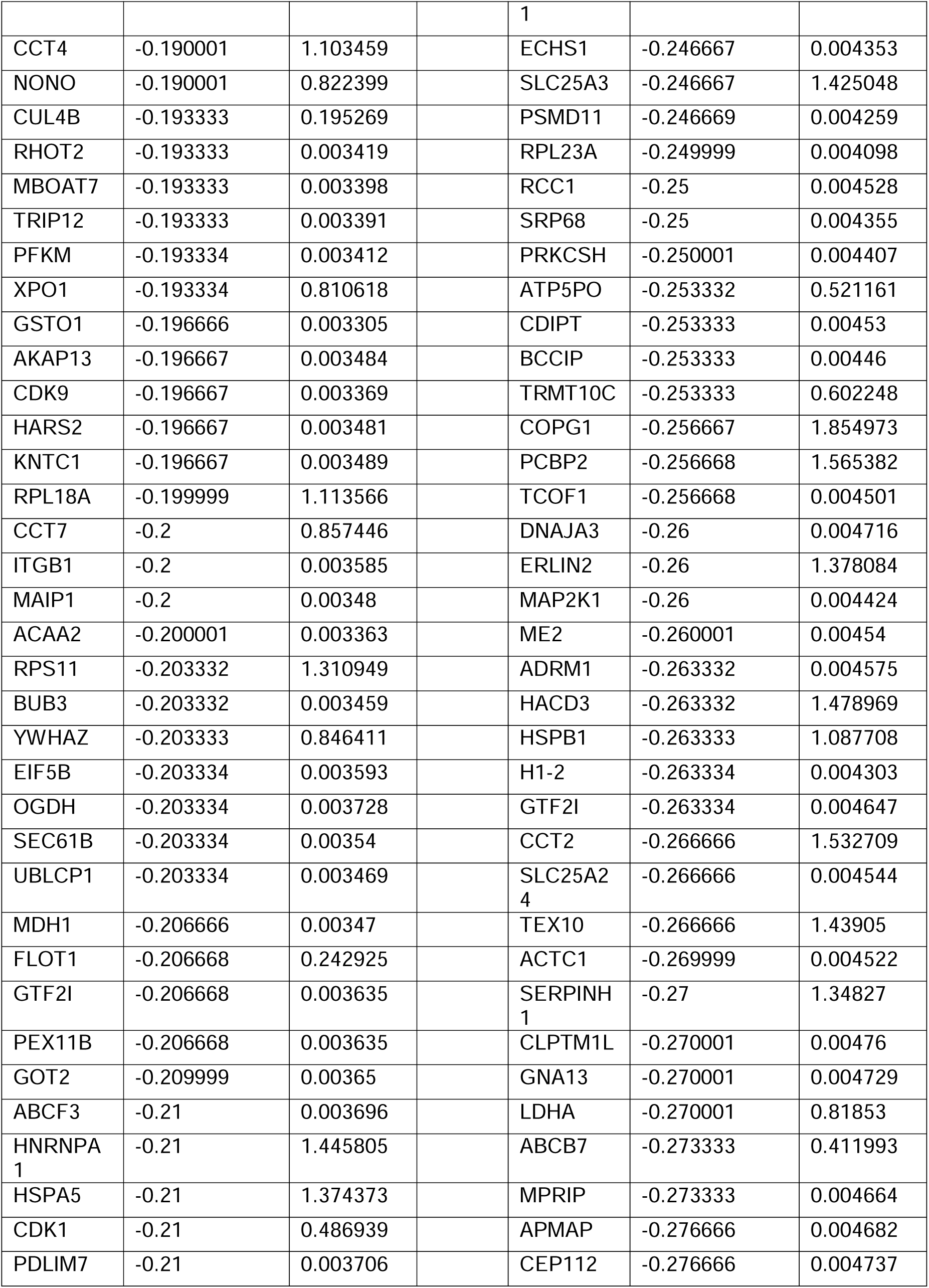
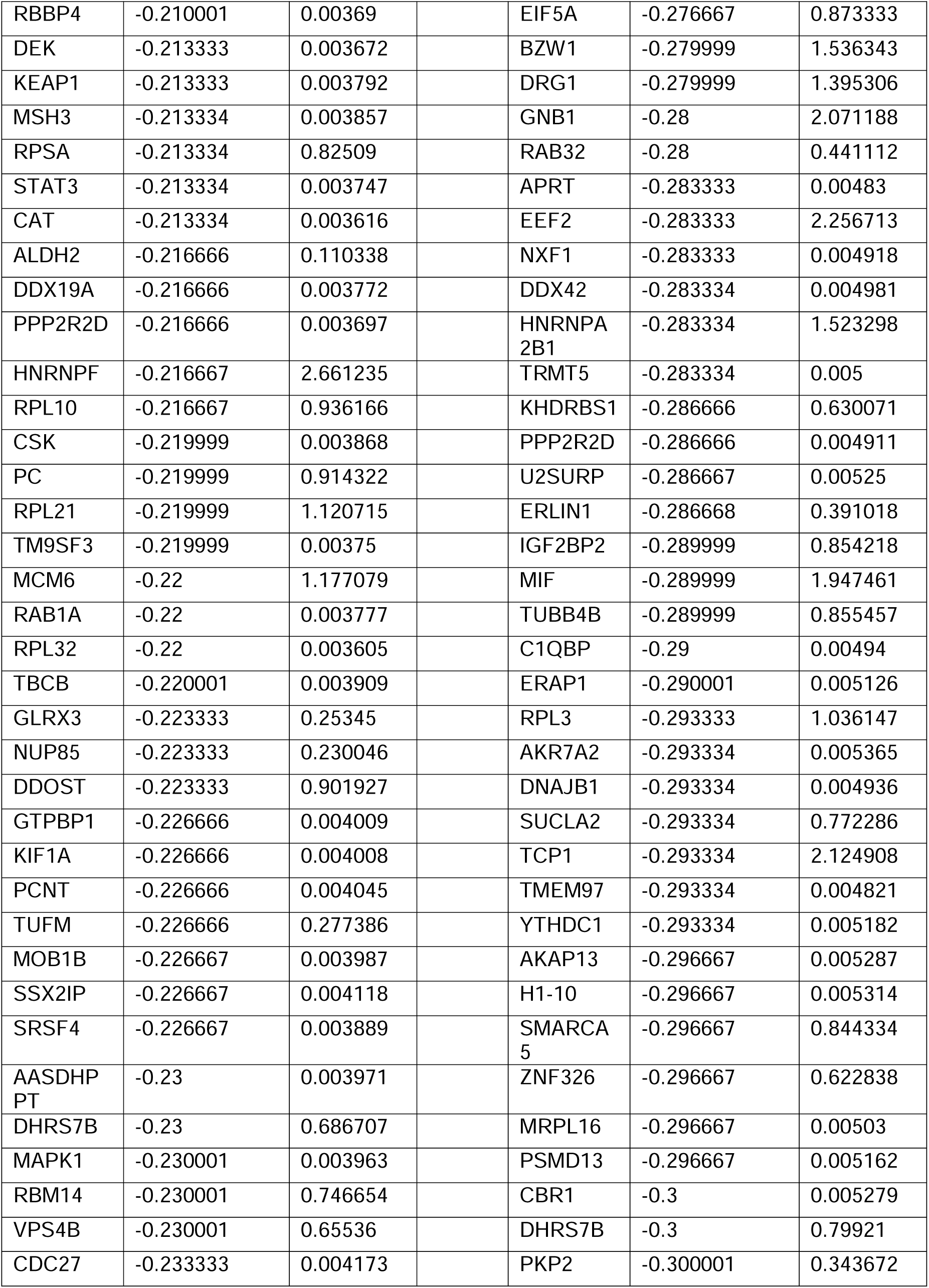
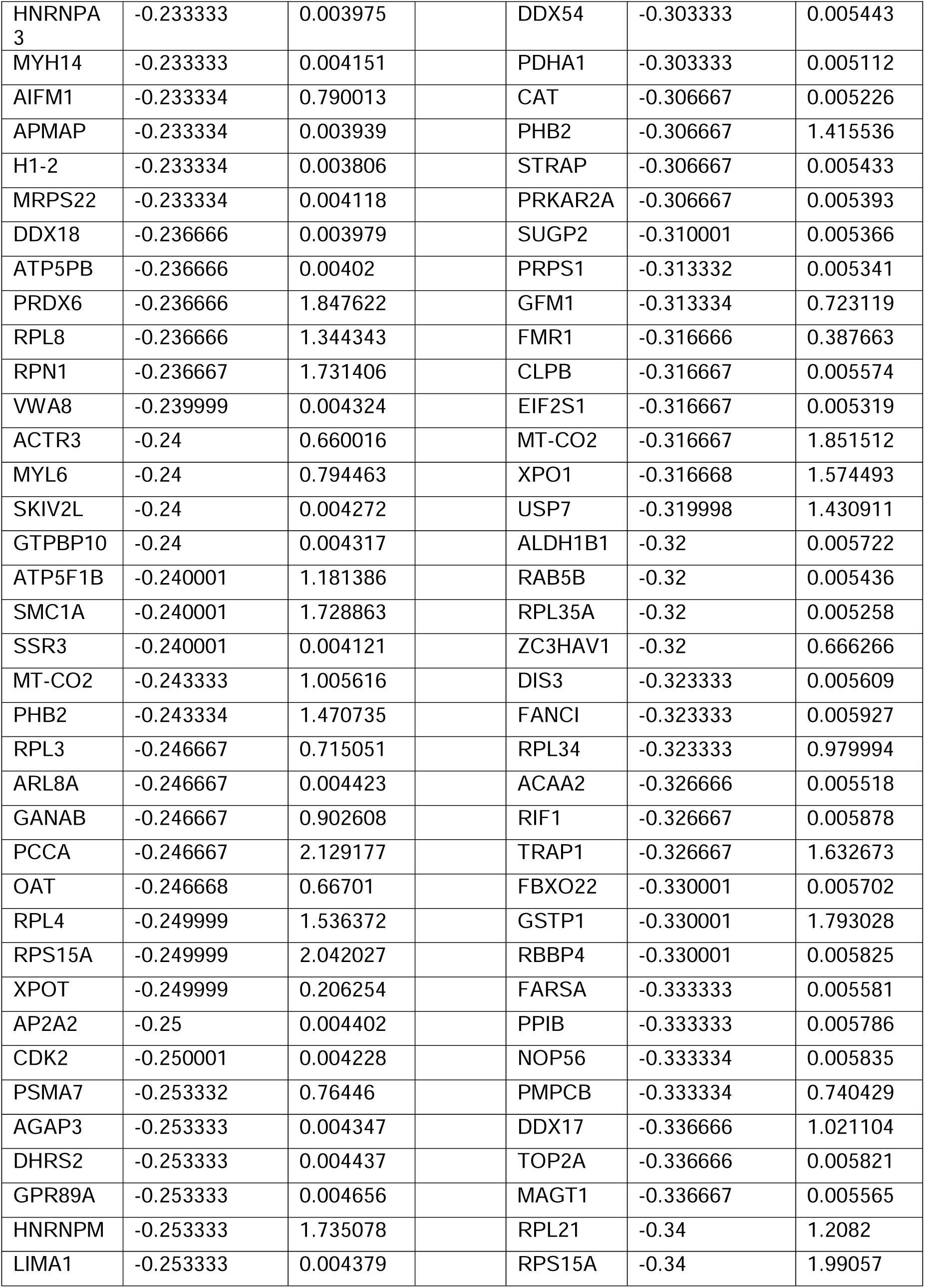
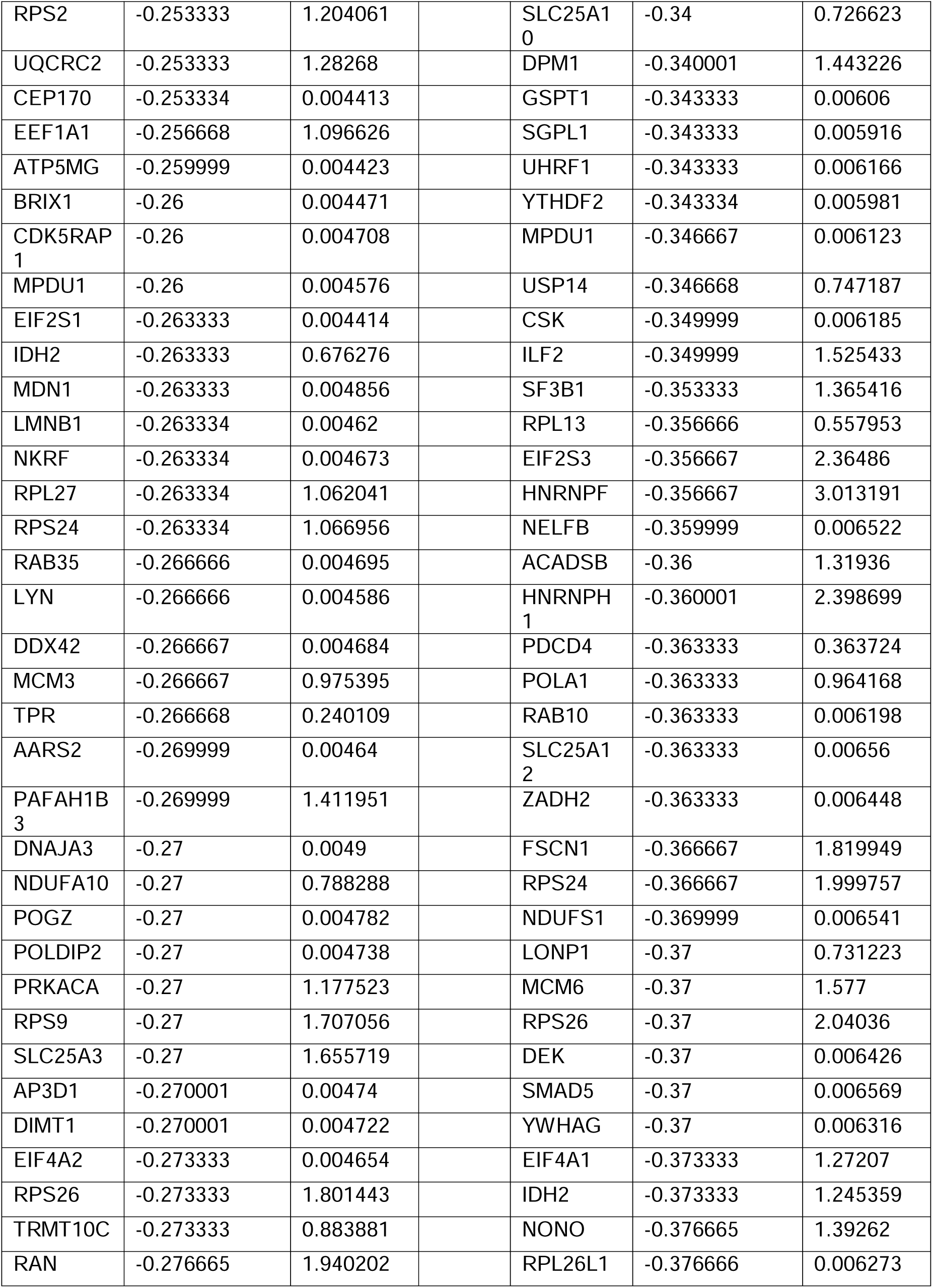
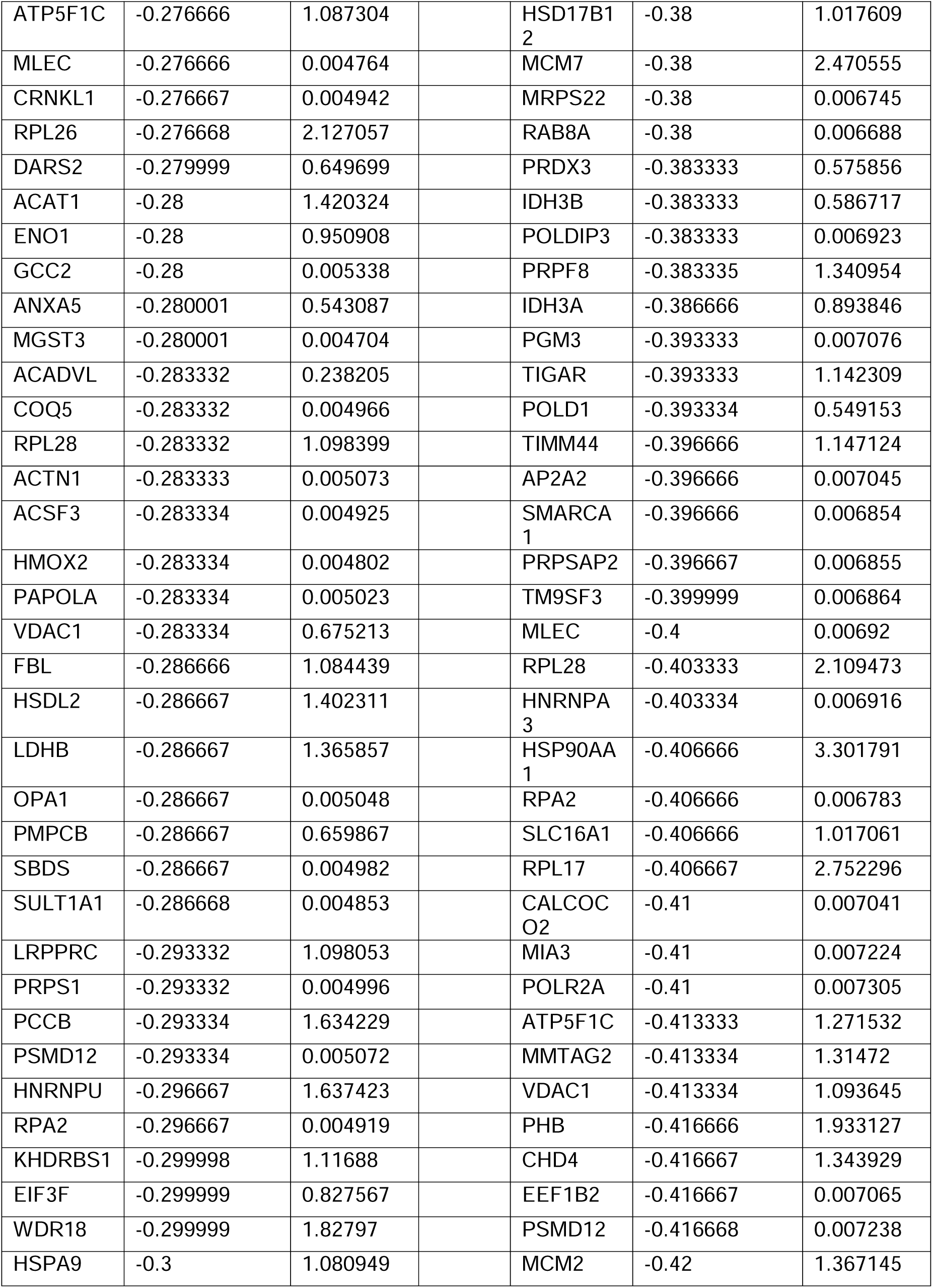
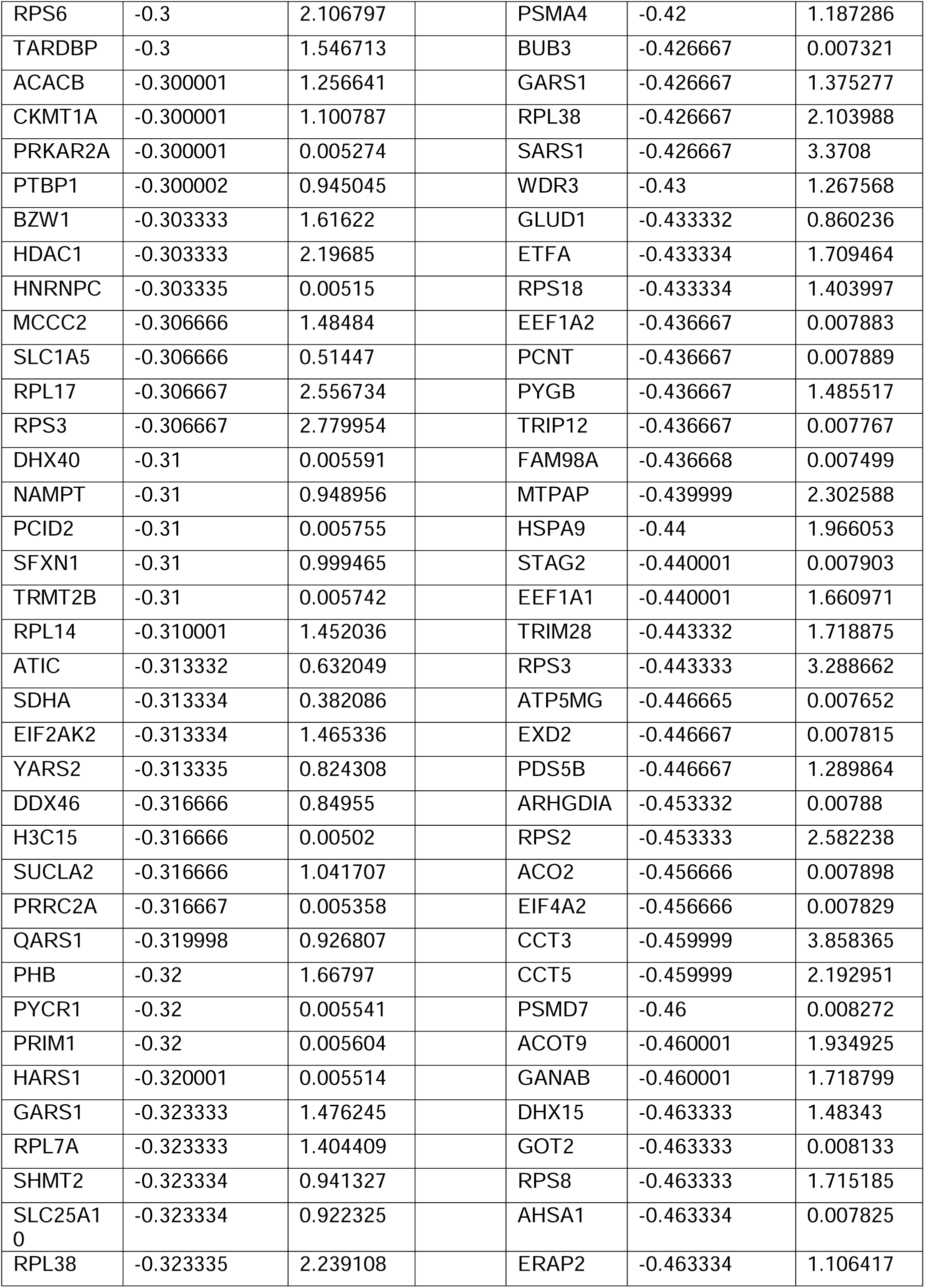
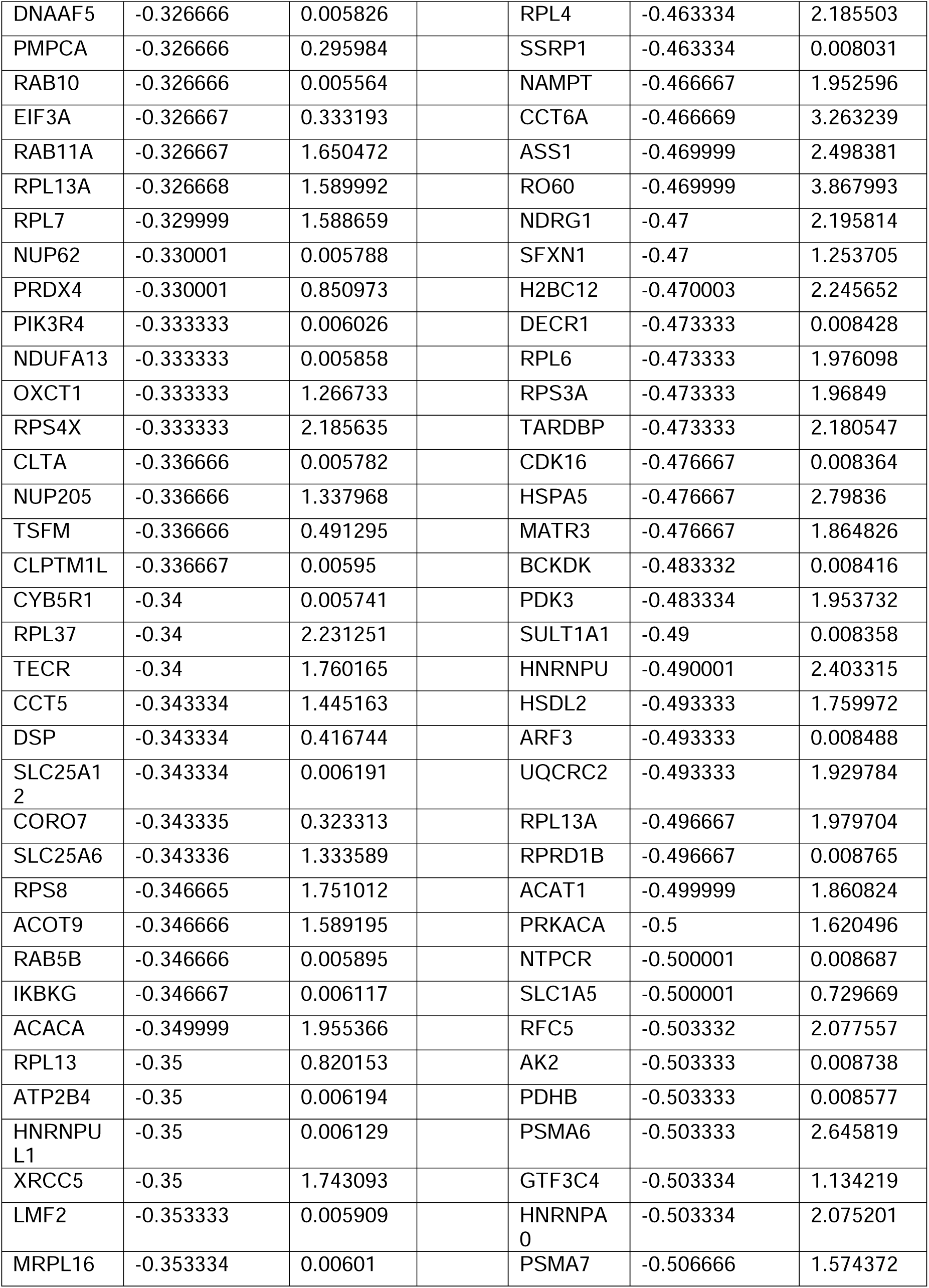
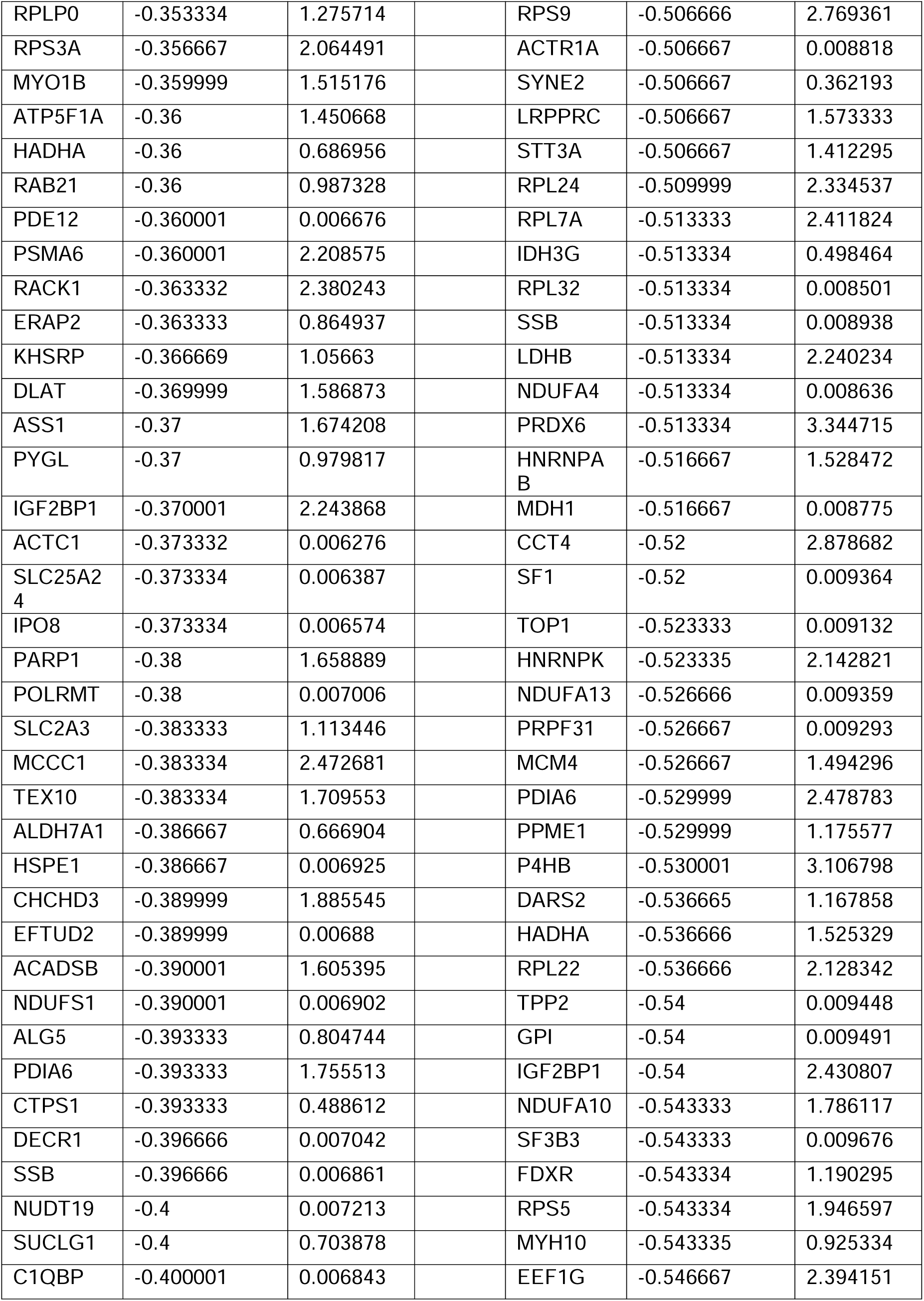
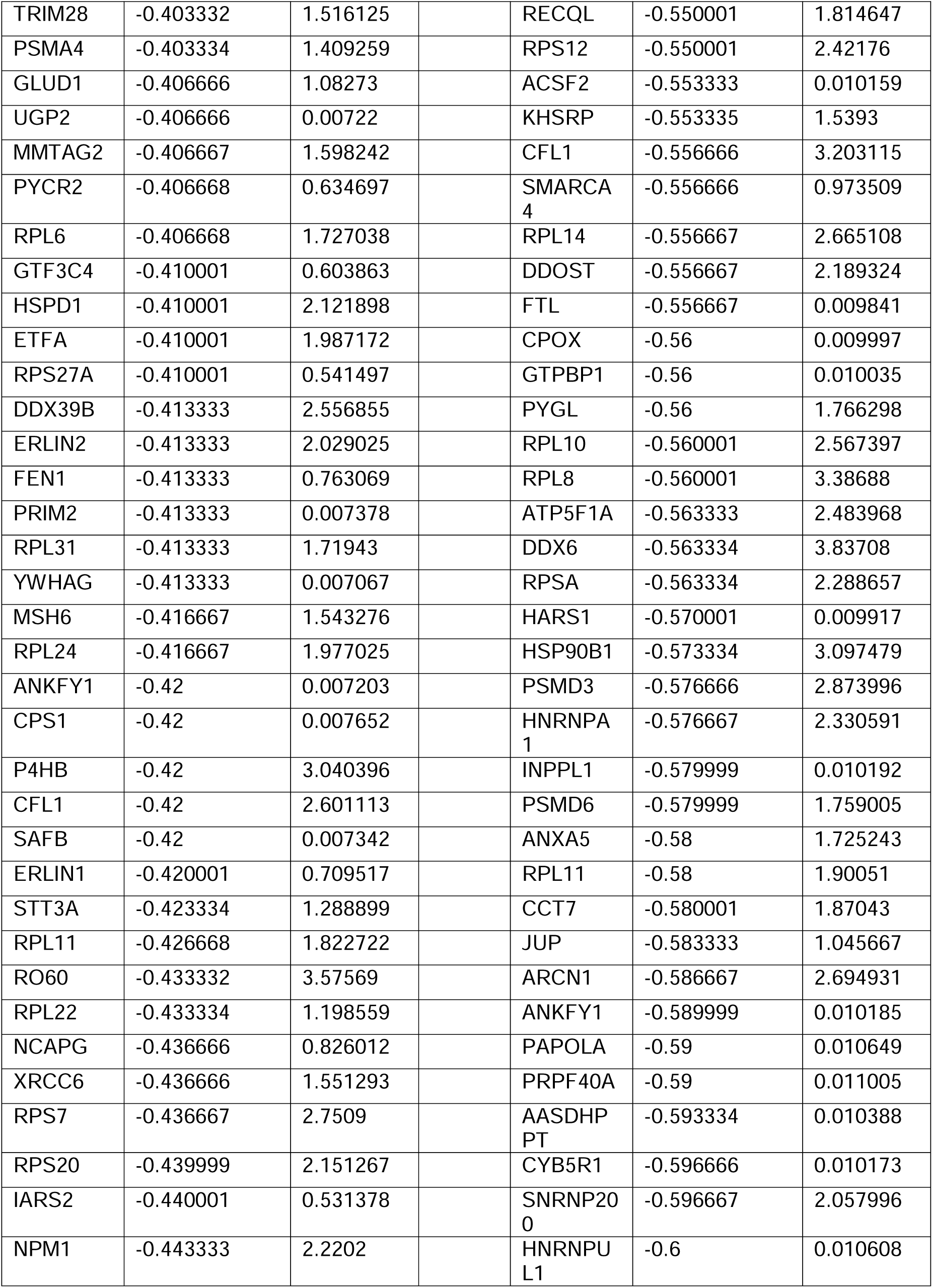
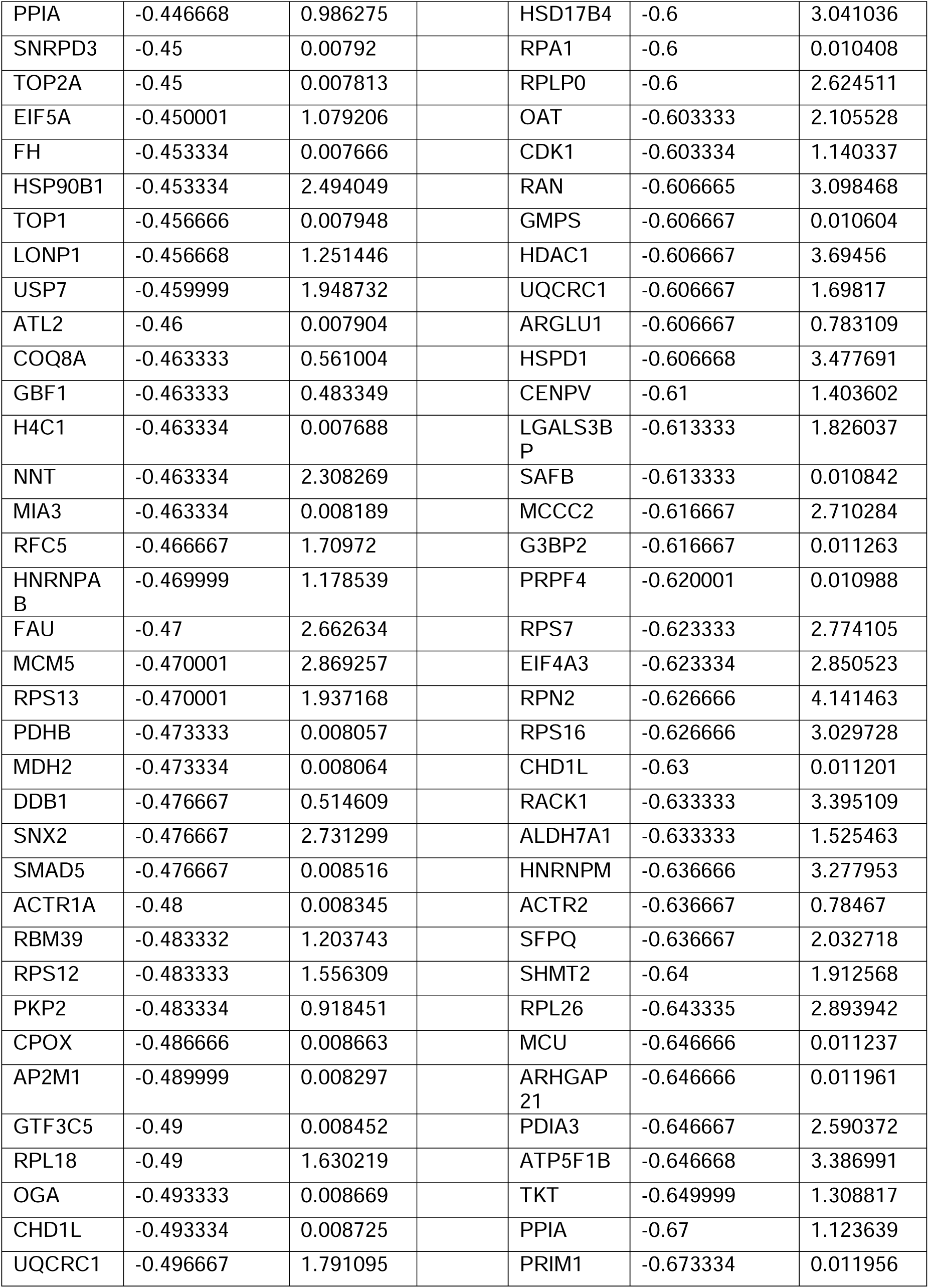
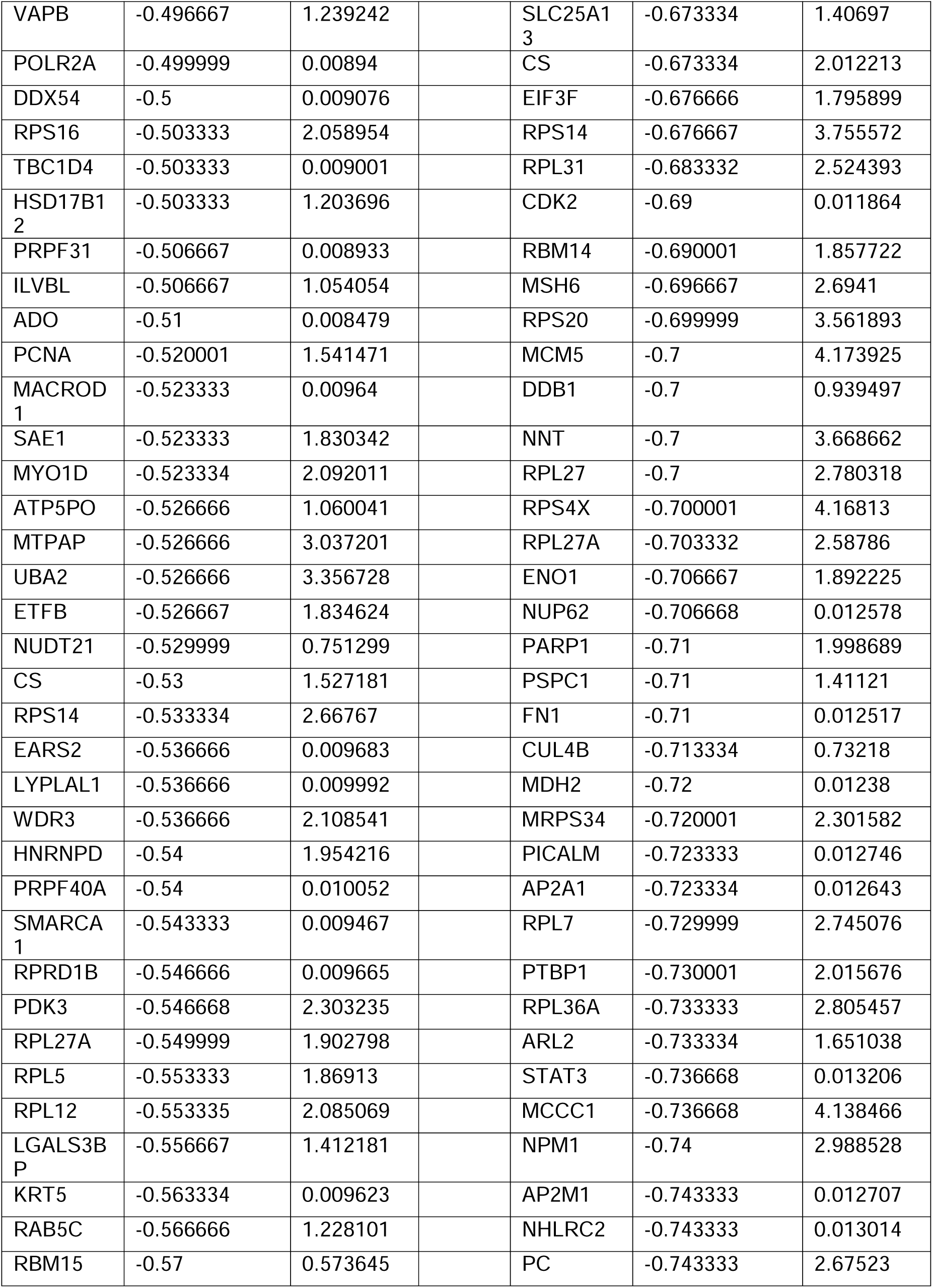
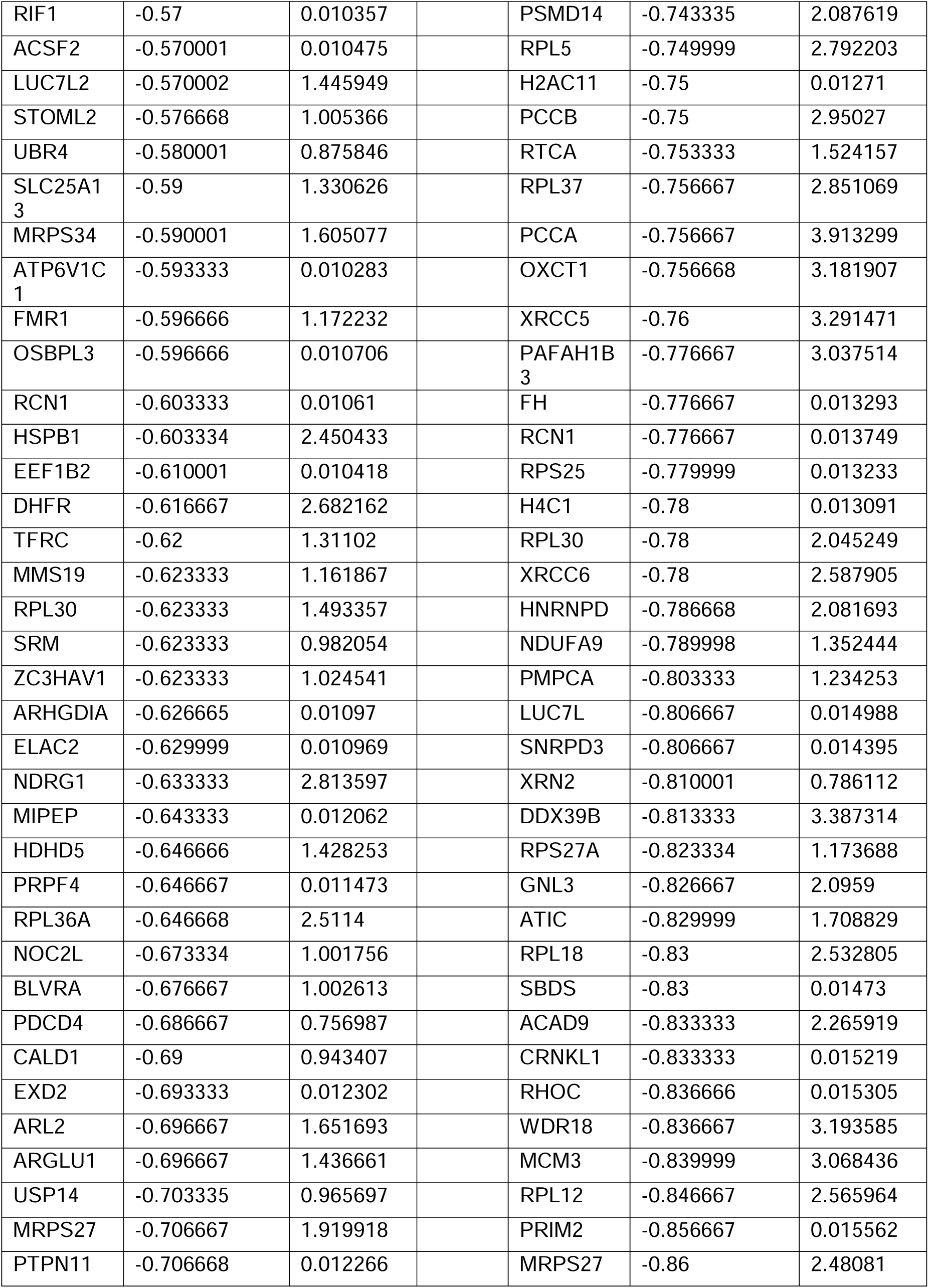
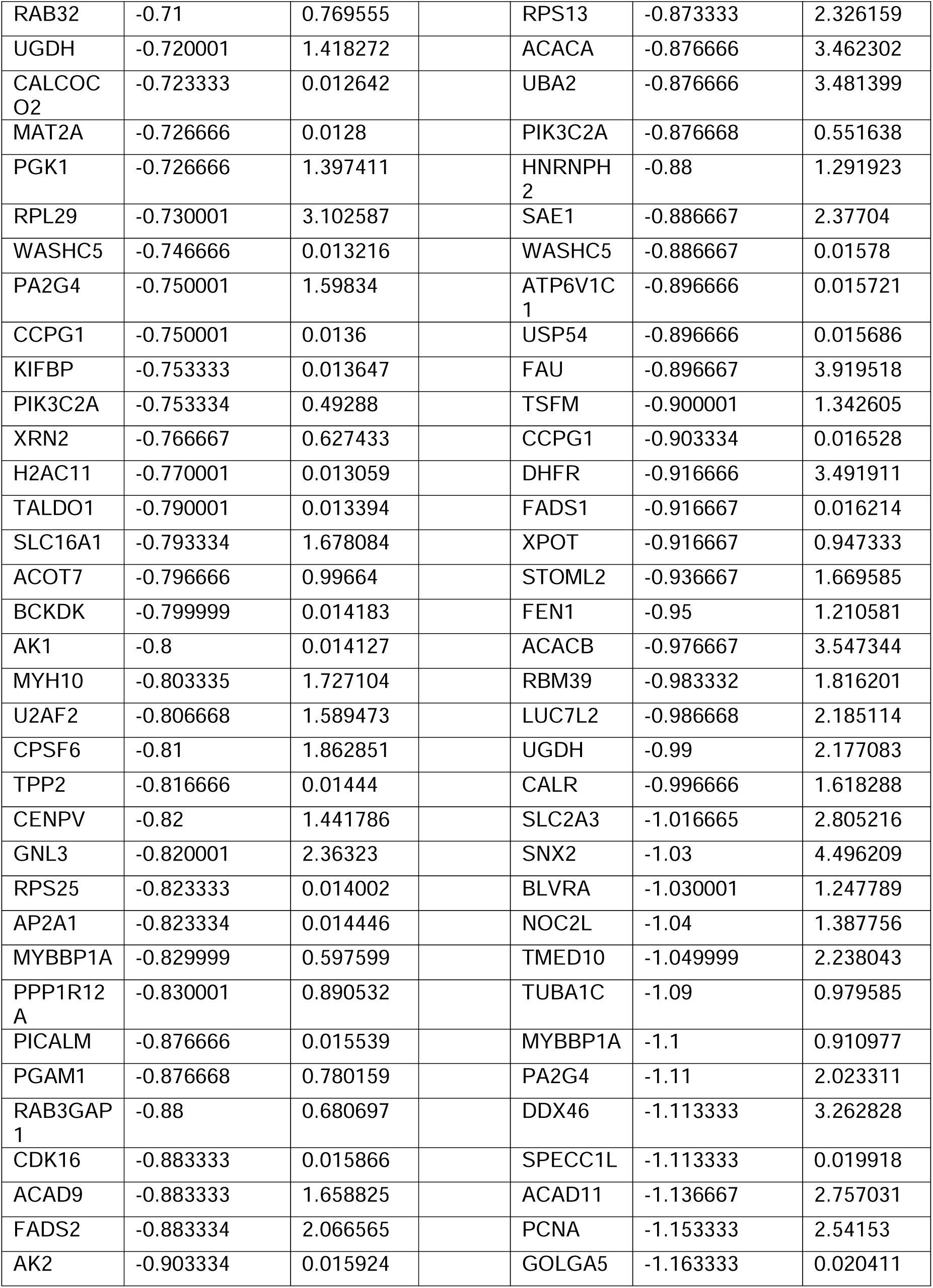
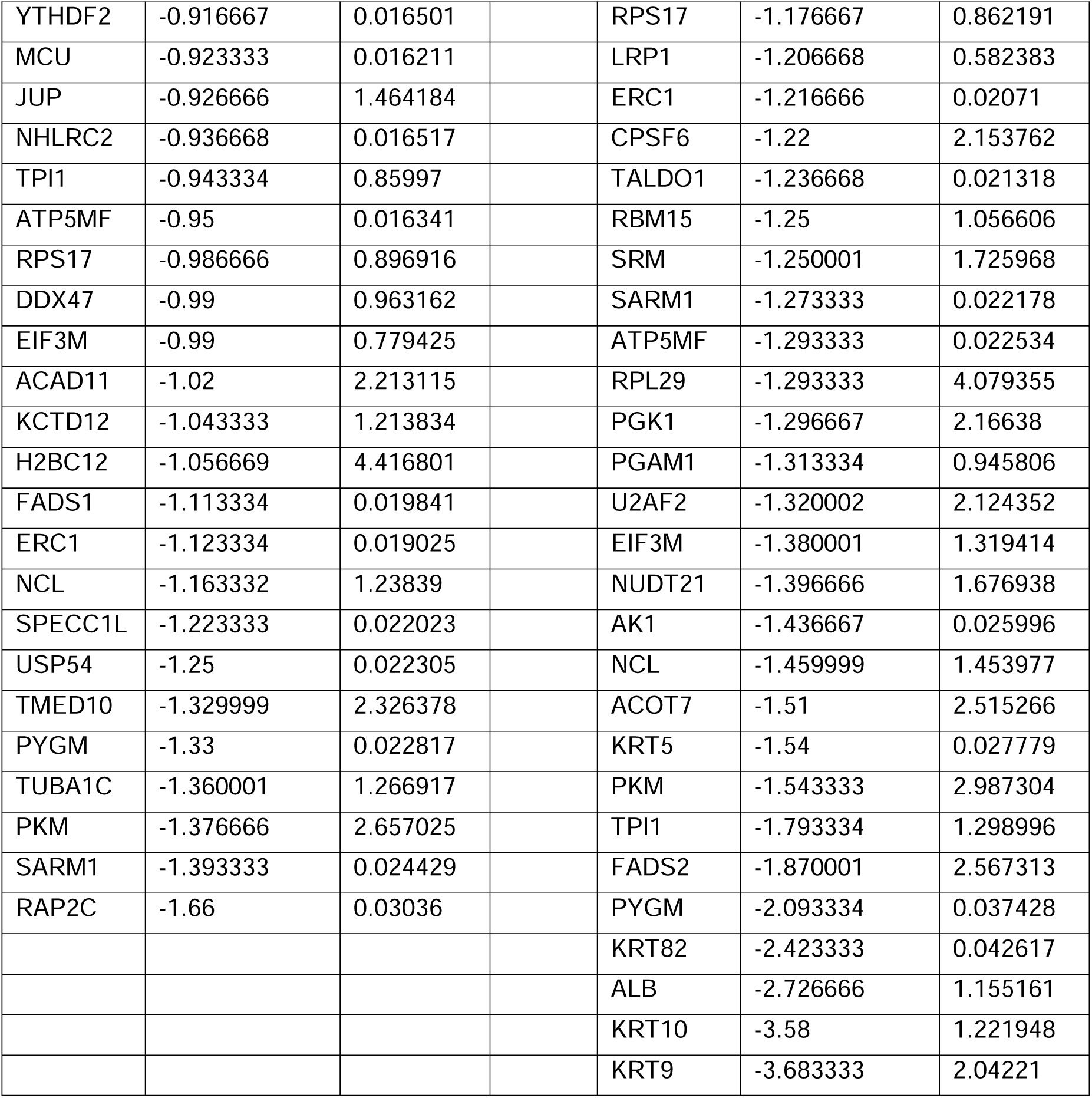
Mass Spectrometry Data.

